# Transgenic menstruating mice reveal the spatial logic of pre-menstrual uterine differentiation

**DOI:** 10.1101/2025.10.08.681007

**Authors:** Çağrı Çevrim, Nicholas J. Hilgert, Aellah M. Kaage, Andrew J. C. Russell, Allison E. Goldstein, Claire J. Ang, Patricia L. Murphy, Enya Miguel Whelan, Sofia G. Bennetts, Kayla C. Lyons, Alan Y. Baez Vazquez, Daniel H. Kim, Daisy A. Hoagland, Laura E. Bagamery, Ana Breznik, Jaina L.R. Gable, Daniela J. Di Bella, Mustafa Talay, Jingyu Peng, Fátima Sanchís-Calleja, Dorothy K. Sojka, Kathleen E. O’Neill, Ruth A. Franklin, Fei Chen, Sean R. Eddy, Kara L. McKinley

## Abstract

During menstruation, the superficial layer of the endometrium differentiates and sheds, while the underlying basal layer is preserved to support repeated regeneration. How differentiation is restricted to the superficial layer remains unknown. Progress in understanding menstruation has been hindered by the lack of tractable animal models, as mice and rats do not menstruate. Here, we present transgenic mouse models that use chemogenetics to induce key anatomical, functional, and transcriptional features of human menstruation. Using these models, we find that premenstrual differentiation is spatially organized through epithelial–stromal crosstalk, resulting in initiation around the luminal epithelium that progresses towards the basal endometrium. Our findings reveal key principles underlying the spatial patterning of premenstrual endometrial differentiation and establish a transgenic platform for menstruation research.

## Introduction

The uterine lining (endometrium) undergoes hundreds of cycles of differentiation, shedding, and regeneration across the lifespan, a rare example of scarless regeneration in mammals. These events are physically segregated within the tissue: the superficial layer (the functionalis) undergoes differentiation that leads to shedding, whereas the underlying layer (the basalis) remains undifferentiated and is retained during menstruation (Critchley et al., 2020). This partition is important for preserving the cellular reservoirs that drive repair, and disruption of the basalis is associated with endometrial scarring (Yu et al., 2008). Although the bilayered structure of the endometrium was described over a century ago (Sekiba, 1923), the cellular mechanisms that compartmentalize pre-menstrual differentiation to the superficial endometrium remain unclear.

Studies of menstrual biology are limited by the available animal models. Menstruation occurs in fewer than 2% of mammalian species, and ethical and technical constraints limit work in the few species that menstruate naturally (Emera et al., 2012). Laboratory mice (*Mus musculus*) share much of the anatomy of the human reproductive tract and undergo cyclic changes in estrogen and progesterone across their four to five day estrous cycle, but they do not menstruate (Cunha et al., 2019; Liu et al., 2020; Winkler et al., 2024). This difference stems from decidualization, the transformation of endometrial fibroblasts that prepares the uterus for pregnancy. In humans, rising progesterone after ovulation drives decidualization in every cycle; when progesterone falls in the absence of pregnancy, the decidualized tissue breaks down and sheds. In mice, as in most non-menstruating mammals, decidualization occurs only in response to an implanting embryo, so in a non-pregnant cycle there is no decidualized tissue to shed (Ramathal et al., 2010) (Figure 1A).

**Figure 1.**
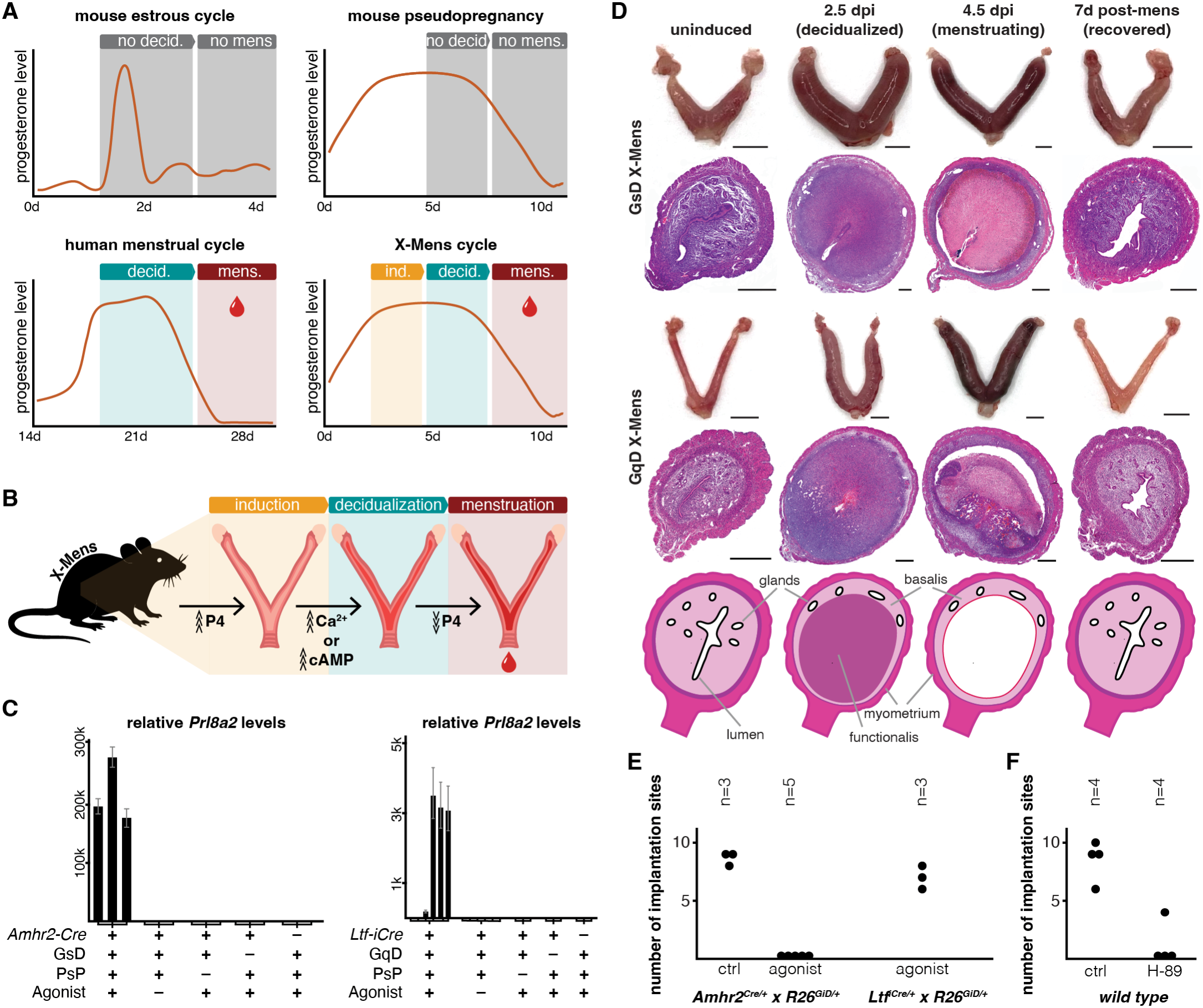
Chemogenetic induction of decidualization and menstruation in mice. **A**. Schematic of progesterone levels and the major events of decidualization and shedding in the wild-type mouse estrous cycle, wild-type mouse pseudopregnancy, the human menstrual cycle, and the X-Mens cycle. Only the human cycle and X-Mens cycle proceed through decidualization and menstruation. d: days; ind: induction; decid: decidualization; mens: menstruation. **B**. Schematic of the X-Mens menstrual cycle, starting with initiation of pseudopregnancy. Decidualization is induced by chemogenetic elevation of calcium (Ca^2+^) in the endometrial epithelium or cyclic AMP (cAMP) in endometrial fibroblasts, and menstruation follows progesterone (P4) decline. **C**. Relative *Prl8a2* mRNA levels in *Amhr2*^*Cre*^ x *R26*^*GsD*^ (left) and *Ltf* ^*iCre*^ x *Tg*^*GqD*^ (right) uteri, together with vehicle-treated, Cre-negative, DREADD-negative, and non-pseudopregnant controls. Each bar is the relative expression for one animal, with error bars showing the propagated standard deviation of technical replicates. PsP: pseudopregnancy. **D**. Representative images of whole-mount uteri and H&E-stained uterus cross sections from *Amhr2*^*Cre*^ x *R26*^*GsD*^ (top) and *Ltf* ^*iCre*^ x *Tg*^*GqD*^ (bottom) mice at sequential stages of the menstruation protocol. Bottom row: schematic illustration of the compartments of X-Mens uteri at each stage. Scale bars are 5 mm for whole-mount uteri images, and 500 µm for uterine cross sections. dpi: days post-induction. **E**. Number of implantation sites at gestational day 6.5 (GD6.5) in *Amhr2*^*Cre*^ x *R26*^*GiD*^ and *Ltf* ^*iCre*^ x *R26*^*GiD*^ mice following agonist or vehicle (ctrl) administration. F. Number of implantation sites at gestational day 6.5 (GD6.5) in wild-type mice following H-89 or vehicle (ctrl) administration. Each dot represents one dam; n indicates the number of dams per group.

In non-menstruating species such as mice, rats, and rabbits, decidualization can be experimentally induced in the absence of an embryo. Classic experiments showed that mechanical trauma to the progesterone-primed uterus, or injection of vegetable oil into the uterine lumen, is sufficient to trigger decidualization (Buxton and Murdoch, 1982; Loeb, 1908), inducing menstruation-like bleeding when progesterone subsequently falls (Finn and Pope, 1984). Many modifications of this procedure have subsequently been developed, with variations in how progesterone is controlled and how oil is delivered (Peterse et al., 2018; Rogers et al., 2025) (see supplementary text). These models recapitulate many aspects of human menstruation and have been instrumental to the field, elucidating roles for hypoxia, immune cells, sex hormones, and inflammation in shedding and regeneration, and enabling work on endometrial stem cells (Armstrong et al., 2017; Brasted et al., 2003; Greaves et al., 2014; Kaitu’u-Lino et al., 2007a; Kaitu’u-Lino et al., 2007b; Kaitu’u-Lino et al., 2010; Kirkwood et al., 2022; Liu et al., 2020; Maybin et al., 2018; Rogers et al., 2025; Shaw et al., 2022). However, these methods rely on non-physiological interventions that act through undefined molecular mechanisms (Liu et al., 2020; Rogers et al., 2025).

We aimed to trigger decidualization and menstruation through targeting a defined molecular pathway. Two second messengers, cyclic AMP (cAMP) and calcium, have been implicated in triggering decidualization. Membrane-permeable cAMP analogs, combined with progesterone, cause cultured human and rodent endometrial stromal cells to adopt hallmarks of decidualization *in vitro* (Brar et al., 1997; Gellersen and Brosens, 2003; Gellersen and Brosens, 2014; Tang et al., 1993; Yee and Kennedy, 1991), and uterine cAMP rises sharply when a decidualization stimulus is applied *in vivo* (Leroy et al., 1974; Rankin et al., 1977). Elevating intrauterine calcium likewise induces decidualization in mice (Buxton and Murdoch, 1982; Sakoff and Murdoch, 1996), and increased intracellular calcium has been reported in human *in vitro* decidualization models (Kawarabayashi et al., 2012). We used chemogenetic control of endometrial cAMP or calcium to enable precise induction of decidualization *in vivo*. Progesterone decline following chemogenetic decidualization triggered hallmarks of menstruation, including vaginal bleeding, shedding of viable endometrial cells, and human-like immune and vascular remodeling, including the formation of early spiral arteries. The endometrium regenerated and preserved reproductive function, and the process could be repeated over successive cycles. Our studies using this system revealed that pre-menstrual differentiation exhibited striking spatial organization despite the elevation of cAMP uniformly across fibroblasts, initiating beneath the epithelium lining the lumen and propagating to the periphery. When the luminal epithelium was ablated, activation of the cAMP cascade in fibroblasts failed to induce decidualization, revealing a requirement for epithelial-stromal crosstalk that constrains where decidualization can initiate. Our data support a model in which licensing of stromal decidualization by the epithelium converts a uniform decidualization cue into a layered tissue, with an inner compartment committed to shedding and an outer compartment preserved for regeneration.

## Results

### Fibroblast cAMP and epithelial calcium drive decidualization in vivo

To place the two second messengers, calcium and cAMP, that have been implicated in decidualization under experimental control, we used Designer Receptors Exclusively Activated by Designer Drugs (DREADDs) (Armbruster et al., 2007). DREADDs are modified G-protein-coupled receptors that elevate specific second messengers upon administration of a small-molecule agonist: the Gs-DREADD (GsD) raises intracellular cAMP and the Gq-DREADD (GqD) raises intracellular calcium. We generated four mouse lines by combining each of the two DREADD alleles (GqD or GsD) with each of two Cre drivers: *Amhr2*^*Cre*^ (Jamin et al., 2002) to target endometrial fibroblasts, or *Ltf* ^*iCre*^ (Daikoku et al., 2014) to target endometrial epithelium (Figure 1B, supplementary text, Figure S1). This design allowed us to ask whether elevation of cAMP or calcium in specific uterine cell types is sufficient to induce decidualization. To model the prolonged progesterone period of the human luteal phase, we induced pseudopregnancy, in which mating with a vasectomized male produces progesterone-producing corpora lutea that persist for approximately eight days (Rudolph et al., 2012) (Figure 1A). We administered DREADD agonists (hereafter agonist; see methods) to pseudopregnant animals, and assessed decidualization 2.5 days post-induction (dpi) by measuring uterine expression of *Prl8a2*, a well-established marker of decidualization (Orwig et al., 1997). Agonist treatment upregulated *Prl8a2* in pseudopregnant animals when cAMP was elevated in endometrial fibro-blasts (*Amhr2*^*Cre*^ x *R26*^*GsD*^) or calcium was elevated in the epithelium (*Ltf* ^*iCre*^ x *Tg*^*GqD*^) (Figure 1C). Animals administered agonist in the absence of pseudopregnancy did not decidualize based on *Prl8a2* expression, nor did genetic controls carrying only one of the two required transgenes (Figure 1C). The reciprocal genetic combinations, elevating cAMP in the epithelium (*Ltf* ^*iCre*^ x *R26*^*GsD*^) or calcium in the fibroblasts (*Amhr2*^*Cre*^ x *Tg*^*GqD*^), also did not induce detectable *Prl8a2* (Figure S2). Thus, specific elevation of epithelial calcium or fibroblast cAMP is sufficient to induce molecular characteristics of decidualization.

*Prl8a2* induction reports the transcriptional onset of decidualization but not the associated tissue remodeling. We therefore asked whether chemogenetically induced tissue acquired the structural hallmarks of decidua. Pseudopregnant *Amhr2*^*Cre*^ x *R26*^*GsD*^ and *Ltf* ^*iCre*^ x *Tg*^*GqD*^ mice treated with agonist underwent pronounced uterine enlargement, which was absent in controls (Figure 1D, Figure S3). For histological analysis, we generated cross-sections through the uterine horns. Histology at 2.5 dpi revealed tissue architecture characteristic of decidualization (Ramathal et al., 2010), including stromal cells with expanded cytoplasm and enlarged nuclei, and reduced intercellular space (Figure 1D, Figure S4). Induced tissue in both strains showed increased vascular density, assessed by CD31 (PECAM-1) staining, together with elevated HIF1A and CA9, markers of the hypoxic environment that accompanies decidual growth (Figure S5). These phenotypes align with observations of mouse decidualization during pregnancy (Kim et al., 2013) and after oil induction (Maybin et al., 2018; Zuo et al., 2015). We observed that the expanding decidua displaced the endometrial glands and luminal epithelium toward the endometrial periphery (Figure S6), as occurs in both pregnancy and oil-induced decidualization (Ang et al., 2026; Yuan et al., 2018). Together, these data reveal that cell type-specific calcium or cAMP elevation is sufficient to reproduce the morphological hallmarks of decidualization *in vivo*.

Finally, we assessed the relevance of our engineered approach to physiological decidualization. To test whether fibroblast cAMP is required for decidualization during normal pregnancy, we used an inhibitory DREADD (GiD) (Zhu et al., 2016) to lower cAMP during the implantation window. We treated *Amhr2*^*Cre*^ x *R26*^*GiD*^ females with agonist during the implantation period (gestational day 3.5 to 4.5) and assessed their uteri on gestational day 6.5 (GD6.5). We did not detect decidual swellings in agonist-treated animals, whereas *Ltf* ^*iCre*^ x *R26*^*GiD*^ females were unaffected (Figure 1E). Inhibiting the principal effector of cAMP, protein kinase A, by systemically delivering H-89 at the same time window, produced a comparable loss of decidual swellings in wild-type females (Figure 1F). The convergence of the pharmacological and chemogenetic results identifies fibroblast cAMP as a physiological driver of decidualization, supporting the use of this node for decidual induction on demand.

### Chemogenetic decidualization enables repeated, human-like menstruation in mice

Approximately 3 dpi with agonist, pseudopregnant *Amhr2*^*Cre*^ x *R26*^*GsD*^ and *Ltf* ^*iCre*^ x *Tg*^*GqD*^ mice exhibited vaginal bleeding (Figure 2A-D). Bleeding began approximately 8 days after the onset of pseudopregnancy, when progesterone declines due to luteolysis (Atkinson and Hooker, 1945; Rudolph et al., 2012), the same hormonal event that triggers human menstruation. Bleeding lasted approximately 4 days, and the protocol could be repeated as early as 4 days after bleeding ceased (Figure 2, C and D). Bleeding occurred in 93% of induced *Amhr2*^*Cre*^ x *R26*^*GsD*^ and 70% of induced *Ltf* ^*iCre*^ x *Tg*^*GqD*^ animals (Figure 2E) and could be induced by multiple DREADD agonist regimens (supplementary text, Figure S7). We name the DREADD-based menstrution approach Chemogenetic Induction of Menstruation (X-Mens) and refer to the strains based on their respective DREADDs: GsD X-Mens and GqD X-Mens. The biological and practical distinctions between the two strains, their limitations, and comparisons to oil-based menstruation models are detailed in the supplementary text.

**Figure 2.**
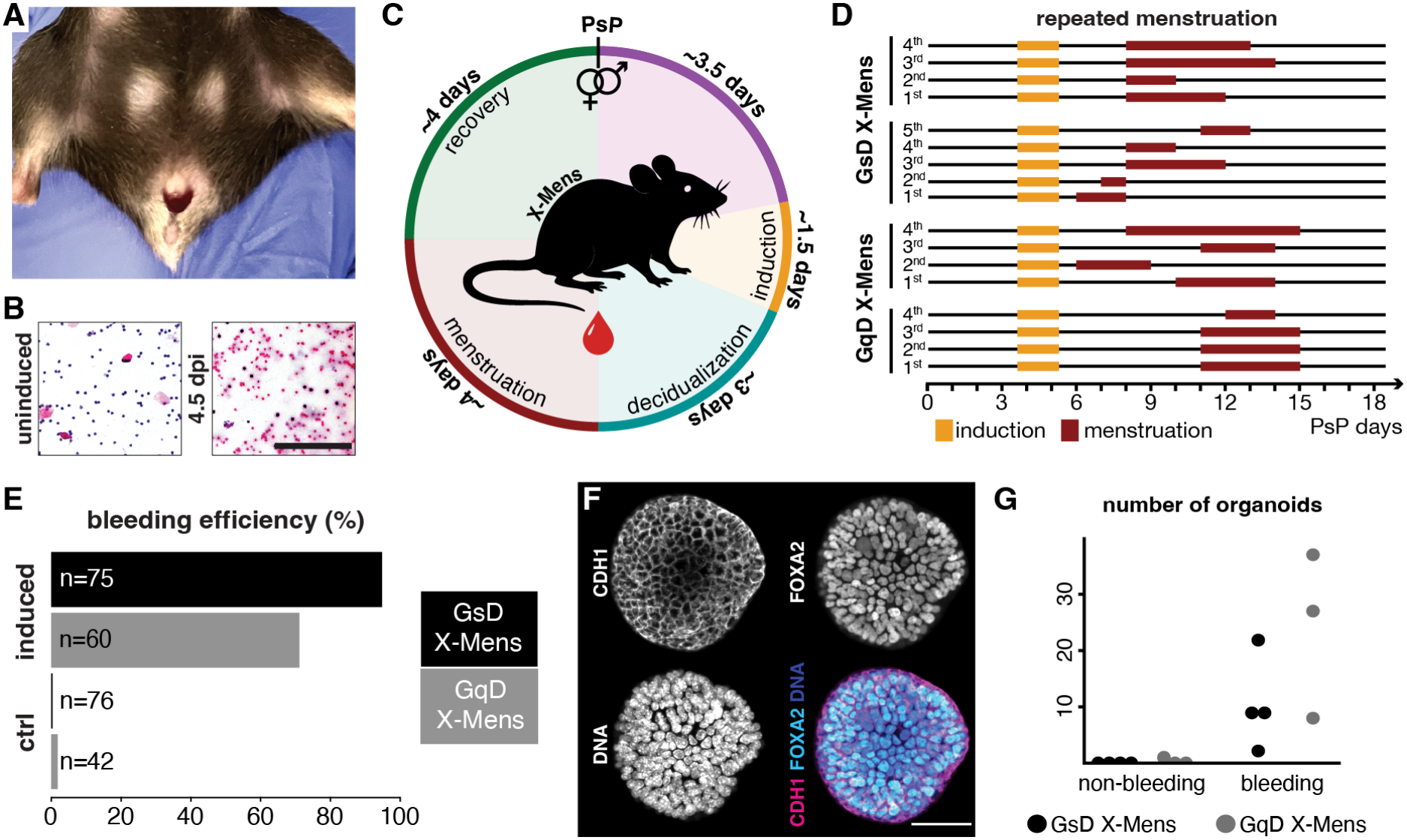
Menstrual bleeding, cyclicity, and tissue shedding in X-Mens mice. **A**. Representative image of an X-Mens mouse exhibiting vaginal bleeding after induction and progesterone decline. **B**. H&E staining of vaginal lavage samples from X-Mens mice before induction (left) and during menstruation at 4.5 days post induction (dpi) (right). Red blood cells are visible in pink. Scale bar, 200 µm, applies to both panels. **C**. Schematic of the X-Mens menstrual cycle, starting with initiation of pseudopregnancy (PsP), with the approximate duration of each phase indicated. **D**. Bleeding episodes in two GsD (top) and two GqD (bottom) X-Mens mice across repeated menstruations. Each bracketed block is one animal; within a block, successive cycles run from bottom to top, and each is aligned to the day of pseudopregnancy initiation for that cycle. Orange bars, induction period; red bars, bleeding. **E**. The percentage of animals showing detectable vaginal bleeding by guaiac test after induction, in GsD and GqD X-Mens mice treated with agonist or vehicle (ctrl). n indicates the number of animals per group; only animals going through the pipeline for the first time are included. The GsD X-Mens data are also presented in Figure S7. **F**. Representative immunofluorescence images of an organoid derived from menstrual effluent of X-Mens mice, stained for epithelial marker CDH1, glandular epithelial marker FOXA2 and DNA (DAPI). Individual channels are shown in grayscale. Scale bar, 50 µm. **G**. Quantification of endometrial organoids derived from vaginal lavages of non-bleeding and bleeding GsD and GqD X-Mens mice. Each dot represents organoids cultured from pooled vaginal lavages of two mice.

Histological analysis of bleeding uteri revealed red blood cells infiltrating the decidual tissue, consistent with localized vascular breakdown (Figure 1D, Figure S4). The superficial decidual layer subsequently detached and shed, leaving behind non-decidual fibroblasts and glandular epithelial cells (Figure 1D, Figure S4 and S8). Human menstruation involves a similar partition, in which the superficial functionalis is shed and the underlying basalis is retained (Cousins et al., 2021; Garry et al., 2009; Salamonsen et al., 2021; Sekiba, 1923). As human menstruation entails shedding of viable cells, we tested whether X-Mens menstrual effluent contained living cells that could be expanded under established endometrial organoid conditions (Turco et al., 2017). Organoids expressing the endometrial epithelial markers CDH1 and FOXA2 developed from vaginal lavages of both X-Mens strains collected during bleeding, but not from lavages collected from the same animals before induction (Figure 2, F and G), indicating that viable endometrial epithelium is shed during menstruation but not in the uninduced state. Chemogenetic decidualization therefore results in vaginal bleeding accompanied by shedding of live superficial endometrium.

We further assessed whether X-Mens mice could under-go functional regeneration after menstruation and menstruate repeatedly. One week after bleeding ceased, uterine morphology was indistinguishable from uninduced animals (Figure 1D, Figure S4). Menstruation did not alter subsequent pregnancy outcomes in either strain: GqD X-Mens females supported pregnancies to term regardless of prior induction, and post-induction GsD X-Mens females supported implantation but not pregnancy maintenance, consistent with the outcomes observed in uninduced animals of this strain (supplementary text, Figure S9). We repeated the menstruation protocol up to five times in individual animals, with no substantial change in uterine morphology across cycles (Figure 2D and Figure S10), although some GsD X-Mens mice exhibited vaginal swelling after four or more cycles, a phenotype not observed in GqD X-Mens mice (see supplementary text). We leveraged the capacity of our models to undergo repeated cycles of menstruation to assess intra-individual variability in bleeding duration, which is a common feature of menstruation in humans (Belsey and Pinol, 1997; Dasharathy et al., 2012). Bleeding duration exhibited both inter-individual variability and cycle-to-cycle variability within the same animal (Figure 2D, Figure S10 and S11). Reasoning that this variation reflected differences in the timing of progesterone decline at the end of pseudopregnancy, we administered the progesterone receptor antagonist mifepristone at 2.5 dpi, an approach used to induce menstruation in classical mouse models (Xu et al., 2007). Mifepristone synchronized and shortened bleeding: all animals initiated bleeding at 3.5 dpi, one day after mifepristone administration, and most bled for a single day (Figure S11), consistent with the onset and duration of bleeding being set by the kinetics of progesterone withdrawal.

### The X-Mens endometrium recapitulates pre-menstrual immune and vascular remodeling

Having established that X-Mens reproduces the anatomy and physiology of menstruation, we asked whether it reproduces the cellular changes that precede shedding. These dynamics have been difficult to study in humans, owing to the scarcity of full-thickness peri-menstrual tissue from healthy donors and to substantial inter-individual variation arising from genetic and life-history differences. X-Mens enables reproducible, precisely timed sampling in isogenic animals raised under uniform conditions with matched reproductive histories.

We collected GsD X-Mens uteri at six time points: uninduced, decidualizing (1.5 dpi early, 1.5 dpi late, 2.5 dpi, 3.5 dpi), and menstrual (4.5 dpi). These and all subsequent samples were collected using the progesterone fluctuations of pseudopregnancy without mifepristone. We generated transverse sections, the equivalent orientation used above for histology, which captures the full thickness of the endometrium, from the lumen at the tissue center to the uterine muscle (myometrium). We subjected these sections to Slide-tags spatially resolved single-nucleus RNA sequencing (Russell et al., 2024). Slide-tags is a spatial transcriptomics method that uses positional barcodes from a patterned bead array to isolate single nuclei from tissue sections while retaining spatial information (Figure 3A-F). Across the six samples we recovered 57,244 nuclei, with a median of 3,131 transcripts and 1,758 genes per nucleus (Figure S12; see methods). Cell types were distinguished based on Leiden clustering, an unsupervised method that partitions cells by transcriptional similarity (Traag et al., 2019), and annotated through established marker genes (Figure 3, A to D, and Figure S12).

**Figure 3.**
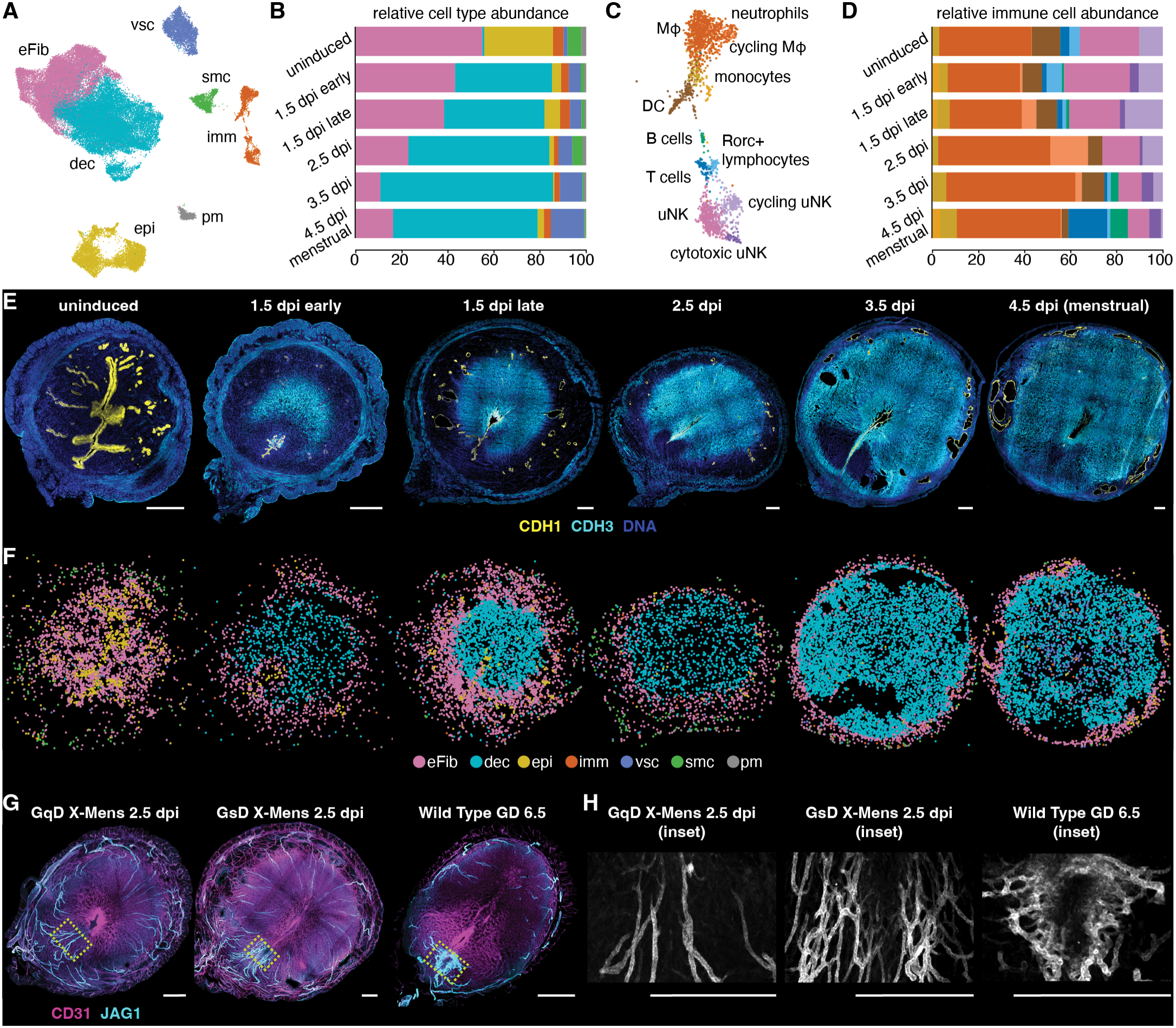
Cellular dynamics and spatial organization of the X-Mens endometrium across the menstrual cycle. **A**. UMAP visualization of single-nucleus transcriptomes from GsD X-Mens Slide-tags samples from all time points, colored by cell type. eFib: endometrial fibroblasts; dec: decidual cells; epi: epithelial cells; imm: immune cells; vsc: vasculature cells; smc: smooth muscle cells; pm: perimetrium cells. **B**. Abundance of each cell type as a fraction of all nuclei recovered, at each sampled stage. Colors as in panel A. dpi: days post-induction. **C**. UMAP visualization of the immune compartment from the same samples, colored by immune cell subtype. Mϕ: macrophages; DC: dendritic cells; uNK: uterine natural killer cells. **D**. Abundance of each immune subtype, as a fraction of all immune nuclei, at each sampled stage. Colors as in panel C. **E**. Immunofluorescence images of GsD X-Mens uterine cross sections at sequential stages of the menstruation protocol, stained for the epithelial marker CDH1 (yellow), the decidual marker CDH3 (cyan) and DNA (DAPI, blue). **F**. Spatial maps of the GsD X-Mens Slide-tags samples, colored by major cell type as in panel A. The immunofluorescence staining in panel E and the Slide-tags analysis were performed on consecutive sections of each sample. **G**. Immunofluorescence images of GqD (left) and GsD (middle) X-Mens uterine cross sections at 2.5 dpi, and of a wild-type uterus at gestational day 6.5 (GD6.5, right), stained for the endothelial marker CD31 (magenta) and the arterial marker JAG1 (cyan). **H**. Enlarged views of the JAG1 channel alone (grayscale)in the yellow dashed boxes in panel G. For all fluorescence images, scale bars: 500 µm. Imaging settings, brightness, and contrast were adjusted independently to enable visualization of key structures.

Cell type proportions shifted over the time course in agreement with our histological observations: expansion of the decidual and vascular populations progressively reduced the representation of epithelial, immune, and fibroblast cells as a fraction of the endometrium (Figure 3B). Spatial mapping of the annotated cell types revealed that the decidua formed a central core surrounded by a persistent rim of endometrial fibroblasts; this partition was apparent from 1.5 dpi and maintained through menstruation (Figure 3F, Figure S13). Therefore, decidualization generates two transcriptionally distinct layers of inner and outer endometrium three days before the onset of tissue shedding.

We first examined the immune populations, which are known to be highly dynamic across decidualization (Gellersen and Brosens, 2014; Wang et al., 2020b; Yang et al., 2026). In humans, uterine natural killer (uNK) cells accumulate through the secretory phase and progressively acquire perforin and granzyme B in the days before menstruation (Yang et al., 2026); their cytotoxic granules have been implicated in initiating the endometrial apoptosis that accompanies shedding (Konno et al., 1999; Strunz et al., 2021). Macrophages follow a parallel trajectory, supporting tissue remodeling and angiogenesis earlier in the cycle before contributing to clearance and repair once shedding begins (Thiruchelvam et al., 2013; Thiruchelvam et al., 2016). We quantified endometrial immune composition across our time course (Figure 3, C and D; Figure S14). uNK cells were abundant among endometrial immune cells before induction, and a substantial proportion acquired a cytotoxic signature during early decidualization. During the progression to menstruation, the uNK fraction of the immune population declined as the macrophage fraction increased, including a proliferating macrophage subset that peaked at 2.5 to 3.5 dpi, consistent with an active role in tissue remodeling during this window. Monocytes and dendritic cells remained at low and relatively stable proportions through-out. At the onset of menstruation, we observed an influx of T and B cells, which were minimal at earlier time points. CD45 staining showed an overall reduction in immune cell density within the decidua itself, consistent with previous observations in mice (Ainslie et al., 2026; Li et al., 2022; Yang et al., 2023). Nonetheless, a subset of immune cells including CD64+ macrophages were detectable within the decidua (Figure S15), as reported in humans (Houser et al., 2011). X-Mens mice therefore undergo extensive immune remodeling ahead of shedding that parallels the human secretory phase.

Pre-menstrual vascular remodeling is a further defining feature of the human menstrual cycle (Bellofiore et al., 2018). Beyond the increased decidual vascular density described above (Figure S5), we found that endometrial arteries marked by JAG1 became progressively branched during early decidualization in GsD-but not in GqD X-Mens (Figure 3, G and H). These vessels were structurally similar to the spiral arteries of mouse pregnancy, resembling arteries at embryonic day (E) 6.5 in both branching morphology and clustered localization (Zwierzynski et al., 2026). The development of spiral arteries has been documented in the naturally menstruating spiny mouse but has not been reported in induced mouse models of menstruation (Bellofiore et al., 2018).

To our knowledge, these are the first spiral artery-like vessels described in a non-pregnant mouse uterus. Notably, GsD X-Mens mice express GsD in fibroblast cells but not the vasculature (Figure S1), indicating that elevation of cAMP drives the emergence of these arteries non-cell autonomously. A candidate mediator is *Cxcl12*, a chemokine important for spiral artery development during pregnancy (Zwierzynski et al., 2026) that is transiently upregulated in endometrial fibroblasts during decidualization (Figure S16). Our data indicate that decidualization driven by fibroblast cAMP induces initial phases of spiral artery development.

### A radial differentiation gradient governs the organization of the pre-menstrual endometrium

The classical functionalis-basalis model predicts a binary partition of the endometrial stroma: an inner population that decidualizes and is shed, and an outer population that resists decidualization and is retained (Sekiba, 1923). To determine if additional substructure was present, we investigated fibroblasts and decidual cells at finer Leiden resolutions and identified five fibroblast and four decidual clusters (Figure S17). Although clustering was agnostic to spatial information, when we mapped each cluster back onto its recorded spatial coordinates, each occupied a discrete territory within the endometrium (Figure 4A and Figure S18).

**Figure 4.**
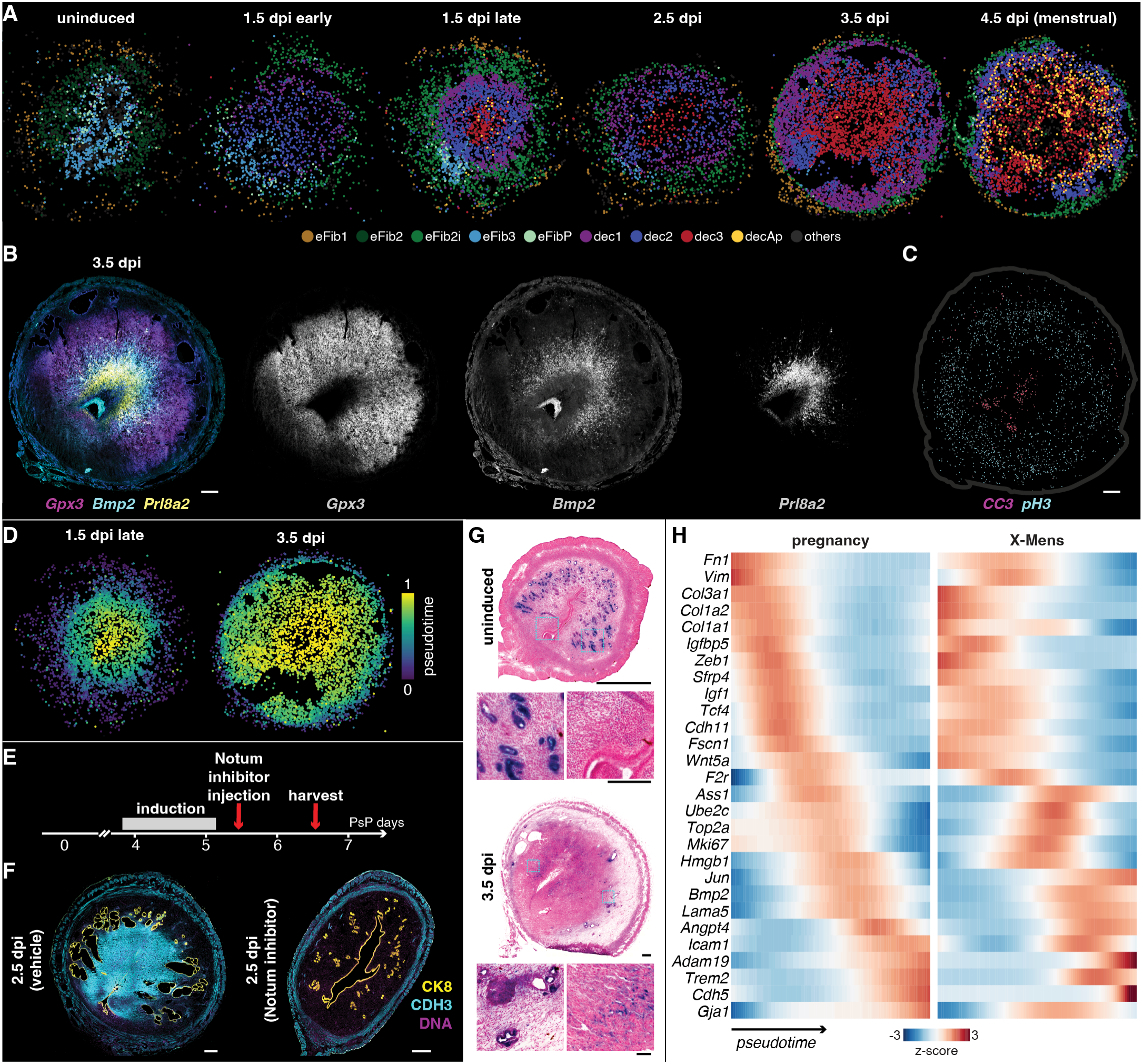
Radial organization of decidual maturation in the X-Mens endometrium. **A**. Spatial maps of GsD X-Mens Slide-tags samples showing endometrial fibroblast (eFib) and decidual (dec) subclusters; all other cell types in gray. dpi: days post-induction; eFib1-3: endometrial fibroblast subclusters 1-3; eFib2i: induced endometrial fibroblasts 2; eFibP: proliferating endometrial fibroblasts; dec1-3: decidual cell subclusters 1-3; decAp: apoptotic decidual cells. **B**. Representative fluorescence in situ hybridization image of a GsD X-Mens uterine cross section at 3.5 dpi. Merge at left; individual channels in grayscale. **C**. Segmented apoptotic (CC3-positive, magenta) and mitotic (pH3-positive; cyan) cells in an adjacent section of the 3.5 dpi sample shown in panel B, plotted at their positions in the tissue. Each dot is one cell; gray outline marks the tissue boundary. These data are reproduced as a panel in Figure S24A. **D**. Spatial maps of Slide-tags samples with eFib and dec cells colored by rank-normalized diffusion pseudotime. **E**. Schematic of the Notum inhibition protocol relative to the day of pseudopregnancy initiation (PsP days). **F**. Representative immunofluorescence images of GsD X-Mens uterine cross sections following administration of vehicle (left) or Notum inhibitor (right), stained for epithelium (CK8, yellow), decidua (CDH3, cyan), and DNA (DAPI, purple). **G**. Senescence-associated β-galactosidase staining of GsD X-Mens uterine cross sections (blue). Nuclei counterstained with nuclear fast red. Insets show the two boxed regions of each overview at higher magnification, in left-to-right order. **H**. Scaled expression of 28 genes along pseudotime in decidualizing cells during mouse pregnancy (left; adapted from Figure 3I of (Yang et al., 2023) with permission from Elsevier) and GsD X-Mens (right). Genes and their row order are taken from the pregnancy panel. Color indicates per-gene z-scores of smoothed expression, clipped at ±3. For all fluorescence and colorimetric images, outset scale bars: 500 µm, insets: 100 µm. Imaging settings, brightness, and contrast were adjusted independently to enable visualization of key structures.

Spatial mapping of the fibroblast clusters revealed a basal fibroblast population (eFib1) that localized to the endometrial-myometrial boundary and persisted through-out decidualization; these cells expressed very low levels of the progesterone receptor *Pgr* (Figure S17C), which may underlie their resistance to decidualization. An initial subluminal population (eFib3), marked by *Rxfp1, Nlgn1* and *Gria4*, diminished as decidualization progressed. A third population (eFib2) localized to the middle region between eFib1 and eFib3 in uninduced endometrium. In induced samples, a distinct but closely related cluster occupied the same position, which we named eFib2-induced (eFib2i; see Methods for the annotation). We also identified a small population expressing proliferation genes such as *Mki67* (eFibP; Figure S17C), present predominantly at early timepoints and overlapping spatially with eFib3 (Figure 4A, Figure S18).

The decidual clusters were positioned at distinct depths within the endometrium; in our transverse sections, this arrangement manifested as the clusters positioned in concentric rings. Upon induction, dec1 formed an outer ring and dec2, which expressed proliferation markers including *Mki67* and *Top2a*, formed an inner ring. At subsequent timepoints, a third cluster, dec3, arose at the center of the dec2 ring. This arrangement persisted, with dec3 expanding and the boundary between decidual and fibroblast territory shifting outward toward the myometrium (Figure 4, A and B, Figure S18). We also detected a fourth cluster of putative apoptotic cells (*Bax*-positive) predominantly in the menstrual sample (decAp; Figure S17C and Figure S18), consistent with reports of apoptosis in earlier mouse models of menstruation (Armstrong et al., 2017). To confirm the transcriptomic data, we performed RNA in situ hybridization of *Gpx3* to mark all decidual cells, *Bmp2* as a marker of dec2 and dec3, and *Prl8a2* for dec3. These experiments confirmed the radial arrangement, with the three transcripts marking nested domains from the outer decidual boundary to the innermost core (Figure 4B, Figure S19). Notably, this concentric patterning could not be explained by spatial variation in the decidualization stimulus: GsD was expressed uniformly across endometrial fibroblasts (Figure S20). Dec1 to dec3 were similarly organized in GqD X-Mens (Figure S19), indicating that this patterning does not depend on how decidualization is triggered. Together, these data show that the pre-menstrual endometrium is organized into nested layers of decidual cells rather than the two compartments of the classical functionalis-basalis model.

The emergence of the decidual clusters over time suggested a model in which decidualization begins near the uterine lumen and spreads outward, so that maturity decreases with endometrial depth (distance from the tissue center). In this model, cells nearest the lumen would decidualize first and would therefore have reached the most mature state (dec3) by later timepoints, the outermost dec1 cluster would comprise cells that had only recently begun to differentiate, and dec2 would contain cells at an intermediate stage. To test this, we performed diffusion pseudotime analysis on endometrial fibroblasts and decidual cells, which orders cells by transcriptomic similarity (Haghverdi et al., 2016) (Figure S21). Pseudotime orders cells but does not specify the direction of the trajectory, so we rooted it in the fibro-blasts, the precursors of decidual cells. Consistent with this orientation, fibroblasts and decidual cells fell at opposite ends of the trajectory (Figure S21). These data are consistent with a model in which decidual clusters are not distinct cell types but rather successive states along a differentiation trajectory. Along this trajectory, fibroblasts first commit to the decidual fate (dec1), proliferate (dec2), and then exit the cell cycle as they acquire the full decidual gene expression program (dec3). This model is consistent with a previous report that decidualization progresses through multiple stages *in vitro* (Rytkönen et al., 2019). We performed spatial mapping of the pseudotime trajectory, which revealed a strong correlation between pseudotime and endometrial depth (Figure 4D), indicating that the decidualization trajectory is tightly spatially organized, progressing radially from the tissue center to the periphery.

We next examined changes in molecular identity that occurred along the trajectory. Dec1 cells lacked canonical decidualization markers including *Prl8a2* and *Bmp2*, raising the question of whether they represent a bona fide decidual state. Searching our dataset for genes expressed across all decidual clusters, we identified *Cdh3, Gpx3* and *Gja1* as pan-decidual markers labeling dec1, dec2 and dec3 (Figure S17C). Antibody staining confirmed that these markers label the decidua in both X-Mens models and in pregnancy (Figure S22). Dec1 therefore represents an early decidual state that canonical markers fail to detect.

As cells acquired dec2 identity, they began to express increasing levels of *Notum*, a secreted inhibitor of Wnt signaling that acts by deacylating Wnt ligands (Kakugawa et al., 2015) (Figure S23). Wnt signaling is required for decidualization (Franco et al., 2011; Li et al., 2007), so enrichment of a Wnt antagonist within decidualizing cells was unexpected. To test whether Notum activity contributes to decidualization, we administered the small-molecule Notum inhibitor LP-922056 (Brommage et al., 2019) to GsD X-Mens animals during early decidualization. Following Notum inhibition, uteri failed to undergo detectable decidualization (Figure 4, E and F). Together with earlier work implicating the Wnt inhibitor *Dkk1* in decidualization (Li et al., 2013; Macdonald et al., 2011), these results indicate that decidualization requires not only Wnt activation but also Wnt antagonism within the differentiating decidua.

We turned to dec3, which pseudotime analysis placed as the most mature population. These cells displayed a further hallmark of maturity: senescence. In both human and mouse pregnancy, a subset of mature decidual cells becomes senescent, and this process is tightly regulated and consequential for pregnancy outcome (Brighton et al., 2017; Hirota et al., 2010). Senescence-associated β-galactosidase staining identified senescent stromal cells specifically within the dec3 region (Figure 4G), supporting dec3 as the terminal state of the gradient.

Having established the organization of molecular markers of cell state within the tissue, we asked whether cellular processes exhibited similar spatial organization. We performed immunofluorescence staining across the time course for phospho-histone H3 (pH3) to mark dividing cells and cleaved caspase 3 (CC3) for apoptosis (Figure 4C, Figure S24). Segmenting and quantifying the images revealed that mitotic cells were initially enriched at the mid-endometrium and shifted progressively deeper ahead of menstruation, toward the endometrial-myometrial boundary. At menstruation, the number of dividing cells fell sharply, and the few remaining pH3-positive nuclei lay at the outer margin, within the retained basalis-like layer (Figure 4C, Figure S24). In contrast, CC3-positive cells peaked at menstruation and were concentrated around the tissue center, aligned with the dec3 and decAp domains. Proliferation and apoptosis are therefore segregated along the same radial axis that organizes the decidual clusters.

Finally, we asked whether X-Mens resembles the natural decidualization that occurs in early pregnancy. Comparing the pseudotime progression of our X-Mens time course with a published pseudotime trajectory of early mouse pregnancy (Yang et al., 2023), we found that genes were induced in a similar sequence in both, progressing from extracellular matrix and fibroblast identity genes, through proliferation genes, to mature decidual genes (Figure 4H). To assess similarities in spatial organization, we then compared decidual clusters in our 3.5 dpi sample (day 7.5 of pseudopregnancy) with a published spatial transcriptomic dataset of the pregnant mouse uterus at gestational day 7.5 (Li et al., 2022). The pregnant endometrium harbors decidual clusters termed decidual zones. Each X-Mens cluster corresponded strongly to one of these zones: dec3 corresponded most closely to the primary decidual zone (PDZ), dec2 to the transitional decidual zone (TDZ), and dec1 to the secondary decidual zone (SDZ) (Figure S25). The decidual zones and X-Mens clusters exhibited similar spatial arrangements, progressing from dec3/PDZ at the tissue center to dec2/TDZ and finally to dec1/SDZ with increasing endometrial depth (Figure S25). Moreover, projecting the gene signatures of the pregnancy decidual zones onto our 3.5 dpi X-Mens sample recovered the same concentric ring organization (Figure S25). X-Mens decidualization therefore reproduces both the temporal and spatial organization of pregnancy decidualization despite the absence of local signals from an embryo.

### X-Mens mice recapitulate the transcriptional programs and radial patterning of human pre-menstrual endometrium

We next asked whether the decidual organization we identified in X-Mens mice is representative of human menstruation. We performed Slide-tags on full-thickness endometrial samples from two human donors without known reproductive disease, one in the secretory and one in the menstrual phase. All major endometrial cell types were present in both samples and occupied their expected spatial positions (Figure 5A-D, Figure S26 and S27). Subclustering of fibroblast and decidual cells resolved eight populations, which were arranged in concentric layers from the myometrium to the lumen (Figure 5C to F). Myometrial fibroblasts (myoFib) localized to the outer myometrium. These cells share the transcriptomic signature of previously identified (Marečková et al., 2024; Ulrich et al., 2024) *C7*+ stromal cells (see supplementary text, Figure S28). Two clusters of endometrial fibroblasts, eFib-a and eFib-b, occupied the deep endometrium. eFib-c and dec-a, located in progressively more luminal zones, formed the transition from fibroblast to decidual states and expressed both fibroblast and early decidual markers. Dec-b cells were the closest cluster to the uterine lumen. Two populations of decidual cells were detected only at menstruation and were present in patches close to the lumen. Dec-c expressed the immediate-early genes *FOS* and *JUN*. Dec-d additionally expressed high levels of heat-shock genes (*HSPA1A, HSPB1, DNAJB1*), hypoxia-associated (*NDRG1, PLOD2*) genes, and a subset of matrix metalloproteases (MMPs; *MMP1, MMP10* and *MMP14*), consistent with shedding cells (Gaide Chevronnay et al., 2009; Nayak et al., 2000). These data suggest a radial organization of fibroblast and decidual cells in the human endometrium, similar to that of mouse.

**Figure 5.**
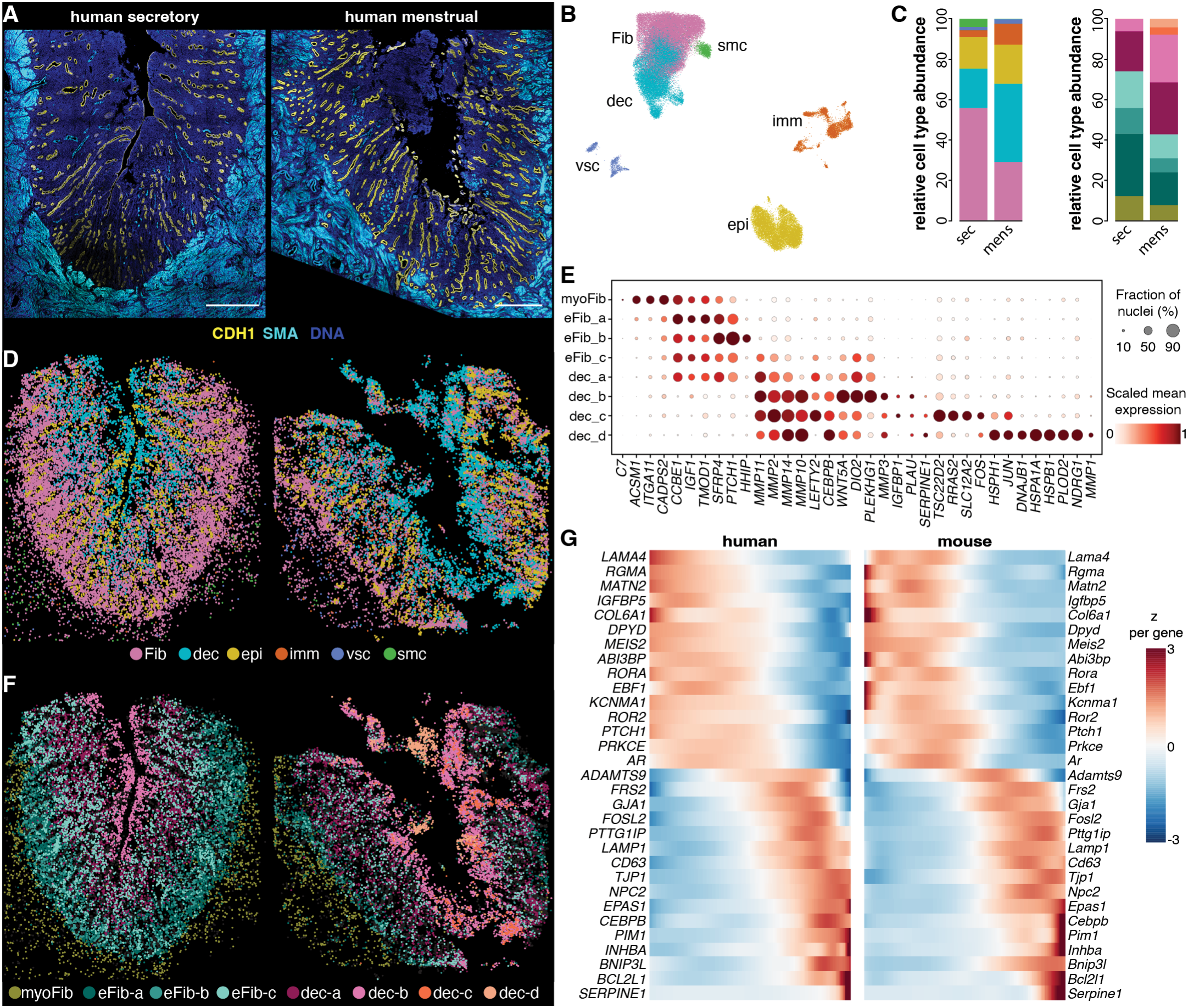
Shared transcriptional programs of pre-menstrual differentiation in human and X-Mens endometrium. **A**. Immunofluorescence images of cross sections of human secretory (left) and menstrual (right) endometrium, stained for the epithelial marker CDH1 (yellow), the smooth muscle marker SMA (cyan), and DNA (DAPI, blue). Scale bars: 1 mm. **B**. UMAP visualization of single-nucleus transcriptomes from both human Slide-tags samples, colored by cell type. The immunofluorescence staining in panel A and the Slide-tags analysis were performed on consecutive sections of each sample. Fib: fibroblasts; dec: decidual cells; epi: epithelial cells; imm: immune cells; vsc: vasculature cells; smc: smooth muscle cells. **C**. Left: abundance of each cell type as a fraction of all nuclei recovered in the human samples, colored as in (B); right: fibroblast and decidual cell subclusters, colored as in panel F. sec: secretory; mens: menstrual. **D**. Spatial maps of all cell types in the secretory (left) and menstrual (right) samples. **E**. Dot plot depicting fraction and normalized mean expression of marker genes for each fibroblast and decidual cell subcluster. myoFib: myometrial fibroblast. **F**. Spatial maps of all fibroblasts and decidual cell subclusters in the secretory (left) and menstrual (right) samples. **G**. Scaled expression of 31 shared pseudotime-associated genes along eFib-to-dec pseudotime in human (left) and X-Mens (right). Genes were drawn from the orthologous pairs ranked in the top 10% by mutual information with pseudotime in both species (379 pairs; Figure S31A), selected to represent the range of dynamics across that set, with preference for genes with known roles in stromal and decidual biology. Genes are grouped by their behavior along pseudotime (falling, switching and rising) and ordered within each group by their position along human pseudotime; the same gene order is used for both species. Color indicates per-gene z-scores computed separately within each species, clipped at ±3.

To compare the transcriptional basis of pre-menstrual differentiation between species, we performed diffusion pseudotime analysis on the human dataset, which produced a trajectory extending from endometrial fibro-blasts to decidual cells (Figure S29), as observed in X-Mens (Figure S21). We identified genes significantly associated with pseudotime progression in each species and partitioned them into those shared between species and those associated with pseudotime in one species only (Figure 5G, Figure S30). Established decidualization markers were among the genes shared between species, including *WNT5A* and *CEBPB* (Supplementary Data S2). Overall, 31% of the top pseudotime-associated genes in humans had X-Mens orthologs in the corresponding mouse set (Figure S31), significantly more than the 12% expected under a null model of independent programs in the two species (hypergeometric test; p ≈ 4.8 × 10^-85^; see Methods). X-Mens mice therefore engage a significant subset of the programs driving human pre-menstrual differentiation.

We next asked whether the human endometrium exhibits the same spatial organization as X-Mens. We co-integrated endometrial fibroblast and decidual cells from the human and X-Mens datasets, using SAMap (Tarashan-sky et al., 2021). SAMap infers cell type homology across species from both sequence similarity and transcriptional nearest-neighbor relationships. The two species mixed thoroughly in the joint embedding, indicating the presence of homologous cell populations between species (Figure 6A, Figure S32). To define the human counter-parts of the X-Mens populations, we assigned each human cell the fibroblast or decidual label and pseudo-time value of its nearest X-Mens neighbors in the cross-species embedding (see Methods). Mapping the transferred cell type identities onto the human tissue positioned decidual cells in the tissue center and fibro-blasts at the periphery, as in X-Mens (Figure 6, B and D). Genes distinguishing outer, basalis-like cells from inner, functionalis-like cells were shared between species (Figure S33), indicating correspondence of both cell identity and location. When we assessed the spatial positions of transferred pseudotime values on the human tissue, pseudotime was strongly correlated with endometrial depth in humans (Figure 6, C and E), as we had observed in X-Mens mice (Figure 4D). As recent work has reported that the human glandular epithelium exhibits a gradient of gene expression along endometrial depth (Lorenzi et al., 2026), we also interrogated the spatial organization of epithelial cells in our human datasets. Leiden clustering of the epithelial cells resolved two glandular populations, one mapping to glands within the decidualized region and the other to glands embedded among fibroblasts that resided deeper in the tissue (Figure S34). Glands within the decidualized region expressed markers of secretory differentiation, including *PAEP, IDO1, MMP7* and *CLDN10*, with little *PGR* or *ESR1*, whereas glands among the surrounding fibroblasts contained cells expressing previously reported markers of basalis epithelium including *AXIN2, IHH, BOC*, and *CDH2* (Fitzgerald et al., 2026). Thus, epithelial and stromal compartments are both stratified along the radial axis and exhibit gene expression boundaries at equivalent depth, raising the possibility that decidualization and epithelial patterning may be coupled. Together, these findings indicate that radial patterning is a core organizing principle of the pre-menstrual endometrium in both species.

**Figure 6.**
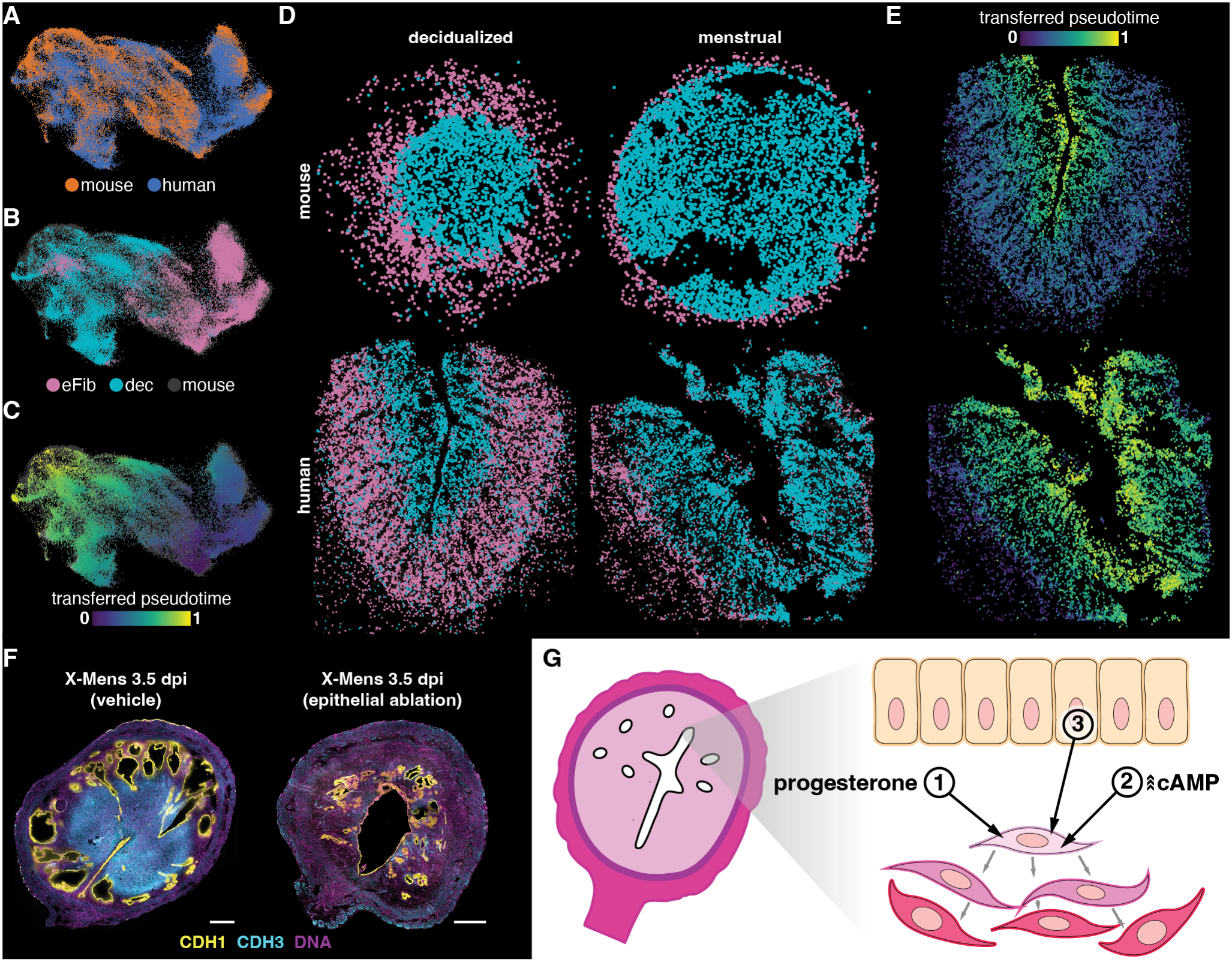
Conserved radial patterning of decidualization and its dependence on the luminal epithelium. **A**. UMAP visualization of human and X-Mens endometrial fibroblasts (eFib) and decidual cells (dec) co-embedded with SAMap, colored by species. **B**. Human cells of the same embedding, colored by eFib and dec labels transferred from X-Mens. **C**. Human cells of the same embedding, colored by rank-normalized diffusion pseudotime transferred from X-Mens. Mouse cells are colored gray. The same data are also displayed in Figure S32B. **D**. Spatial maps of eFib and dec cells in X-Mens (top); and human (bottom) samples, colored as in panel B; human labels are transferred from X-Mens. Columns show decidualized (left; 1.5 dpi late for X-Mens and secretory for human) and menstrual (right; 4.5 dpi for X-Mens) samples. **E**. Spatial maps of human eFib and dec cells from the decidualized (top) and menstrual (bottom) samples, colored by pseudotime transferred from X-Mens through the SAMap embedding. The same data are also displayed in Figure S32F. **F**. Immunofluorescence images of GsD X-Mens uterine cross sections at 3.5 dpi following instillation of vehicle (left) or polidocanol (right) prior to induction. Samples were stained for epithelium (CDH1, yellow), decidua (CDH3, cyan), and DNA (DAPI, purple). Scale bars, 500 µm. Imaging settings, brightness, and contrast were adjusted independently to enable visualization of key structures. **G**. Schematic model of the three signals required for decidualization: (1) systemic progesterone, (2) elevation of cAMP in endometrial fibroblasts, and (3) a permissive signal from the luminal epithelium. Gray arrows and the deepening color indicate the propagation of decidual differentiation away from the lumen.

### Endometrial fibroblasts require an epithelial signal for decidualization *in vivo*

Finally, we asked why decidualization begins adjacent to the uterine lumen. In some decidualization contexts, this origin may be explained by the position of the inducing stimulus. During mouse pregnancy, decidualization begins where the blastocyst contacts the luminal epithelium, and in the GqD model the epithelium likewise initiates the decidualization cascade. In both cases, the signal received by the luminal epithelium must then be relayed to the fibroblasts that manifest the decidual response, and the subluminal fibro-blasts are intuitive first responders based on position. The GsD X-Mens model removes this positional information, as all fibroblasts express the DREADD (Figure S20) and are exposed to circulating progesterone and agonist. In this respect GsD X-Mens resembles the human cycle in which fibroblasts respond to circulating progesterone. Yet in GsD X-Mens, as in the human endometrium, decidualization initiates under the luminal epithelium (Figure 6D and E). We therefore asked whether the epithelium modulates fibroblast responses to decidualization cues. To test this, we induced pseudopregnancy in GsD X-Mens mice and ablated the luminal epithelium shortly before agonist administration using an established protocol (Ang et al., 2026). In this way, cAMP levels were elevated in the fibroblast cells of a de-epithelialized uterus. By 3.5 days after ablation and induction the epithelium had regenerated, yet decidualization was not detectable by tissue morphology or CDH3 staining, whereas sham-ablated animals decidualized robustly (Figure 6F). These data indicate that the luminal epithelium is required for decidualization, even when the decidualization cue is provided directly to the fibroblasts. We therefore propose that decidualization requires the convergence of three inputs in the responding fibroblast: progesterone, elevated cAMP, and a signal from the overlying epithelium (Figure 6G). A short-range epithelial signal would confine the input to the fibroblasts nearest the lumen, potentially accounting for the subluminal origin of the decidual response.

## Discussion

Here, we established X-Mens, a transgenic mouse model that uses defined chemogenetic triggers (cAMP elevation in endometrial fibroblasts or calcium elevation in the endometrial epithelium), to induce decidualization and menstruation on demand. Both the GsD and GqD X-Mens strains exhibit the core anatomical and functional hallmarks of human menstruation: hormonally controlled vaginal bleeding tied to the decline of progesterone, shedding of live endometrial tissue capable of forming organoids, and functional regeneration of the endometrium, including the capacity to support subsequent pregnancies and to undergo repeated cycles of induction. At the cellular and molecular level, we found that X-Mens mice reproduced core transcriptional programs, cell states, and spatial organization of human pre-menstrual differentiation. A further point of similarity with human menstruation is the emergence of arteries in induced GsD X-Mens mice that are structurally similar to the early stages of spiral artery development during mouse pregnancy. The development of spiral arteries outside of pregnancy has long been considered a unique feature of menstruating species (Catalini and Fedder, 2020) that we now show is experimentally inducible. Together, these features establish X-Mens as a robust model of human-like menstruation.

By establishing a tractable, genetically defined, and repeatable model of menstruation in the most experimentally accessible mammalian model organism, X-Mens expands the *in vivo* toolkit available for menstruation research and preclinical development. X-Mens mice combine the precision of cell type-specific genetic control with straightforward, repeatable systemic delivery of agonists to induce physiologically relevant signaling pathways. The capacity to perform repeated cycles makes the system appealing for drug testing as it enables a paired crossover design in which each animal serves as their own control, reducing between-animal variability to increase power and decrease animal usage. The combination of tractability, repeatability, and fidelity to human biology offers new opportunities to dissect menstrual mechanisms within the complex endometrial environment and to understand communication between the menstruating uterus and the rest of the body, advancing our understanding of physiological and pathological processes associated with menstruation.

In addition to serving as a platform for the field, the X-Mens model enabled us to dissect the signaling logic decidualization. Our data support a model for decidualization that requires at least three signals concurrently (Figure 6G). The first signal is elevated systemic progesterone. The second is an induction signal that can be either epithelial calcium or fibroblast cAMP acting through PKA. Our data demonstrate that cAMP is an endogenous mediator of physiological decidualization; whether calcium is an endogenous mediator of decidualization remains to be functionally defined. The third signal required for decidualization is derived from the luminal epithelium. We find that the endometrium fails to decidualize following ablation of the luminal epithelium, even when decidualization is stimulated directly within the fibroblasts. Abrogation of decidualization by luminal epithelium ablation is unlikely to reflect a generalized injury response, as mechanical injury to the progester-one-primed uterus is itself a classical decidualization stimulus (Buxton and Murdoch, 1982; Loeb, 1908; Parkes, 1929). The role of the luminal epithelium in licensing decidualization may underlie the limited transcriptional similarity between decidualization in isolated stromal cultures compared to decidualization *in vivo* (Wang et al., 2020a), and underscores the value of complex co-culture systems that better reconstitute the tissue context (Rawlings et al., 2021). Identifying the molecular nature of this epithelial licensing signal, whether a secreted paracrine factor, a mechanical or adhesive cue, or a local metabolic or oxygen gradient established by the epithelial layer, is an important direction for future work.

We propose that the requirement for the epithelial signal imposes patterning on the decidualization response, consistent with our observations that early decidualization occurs near the luminal epithelium even when the decidualization cue is delivered throughout the fibroblast population in the GsD-X-Mens model. As a result of this outward differentiation process, the pre-menstrual endometrium in both human and mouse is organized as a gradient of decidual states, with progressively less differentiated cells at increasing endometrial depth along the radial axis. Thus, controlled induction of decidualization in X-Mens, combined with validation in human tissue, revealed that the classical bilayered view of the endometrium oversimplifies a radially graded differentiation process. This differentiation gradient was present in all in vivo decidualization systems we examined: GsD-X-Mens, GqD-X-Mens, mouse pregnancy, and human menstruation, supporting this mechanism as a shared patterning principle of decidualization.

Our results enrich the historical view of decidual progression. Noyes, Hertig and Rock’s foundational text for clinical dating of endometrial biopsies describes decidual cells emerging first around the spiral arterioles, and a day later beneath the luminal epithelium (Noyes et al., 1975). We do not view these as competing origins, but as one process observed at two scales. The epithelium may set a permissive zone for decidual initiation along the radial axis; within the responding zone, the first cells to differentiate may be those closest to the vasculature. The vascular-proximal phase may not have been captured in our data because our time course lacked the temporal resolution to capture it, as the mouse completes in two days a progression that unfolds over a week in humans. Future work may reveal still earlier steps of decidual differentiation and further enrich our understanding of the mechanistic underpinnings of these classical histological observations.

A key future frontier is understanding the mechanisms that convert the radial decidual gradient into compartmentalized tissue shedding, a process proposed to involve an array of mediators including matrix metalloproteases and inflammatory signals (Gaide Chevronnay et al., 2009; Wu et al., 2014). Our data regarding the molecular progression of pre-menstrual differentiation and its spatial organization will enable these endeavors. An intriguing possibility is that the extent of propagation of the differentiation wave may contribute to the shedding boundary, potentially resulting in differential extents of shedding in variable cycles while broadly preserving the regenerative capacity of the endometrium across hundreds of reproductive cycles over a lifetime.

## Materials and methods

All material used in this study listed in Table S1.

**Table S1:**
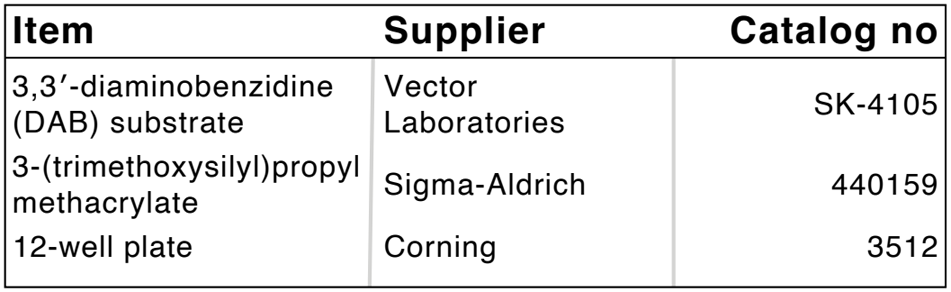
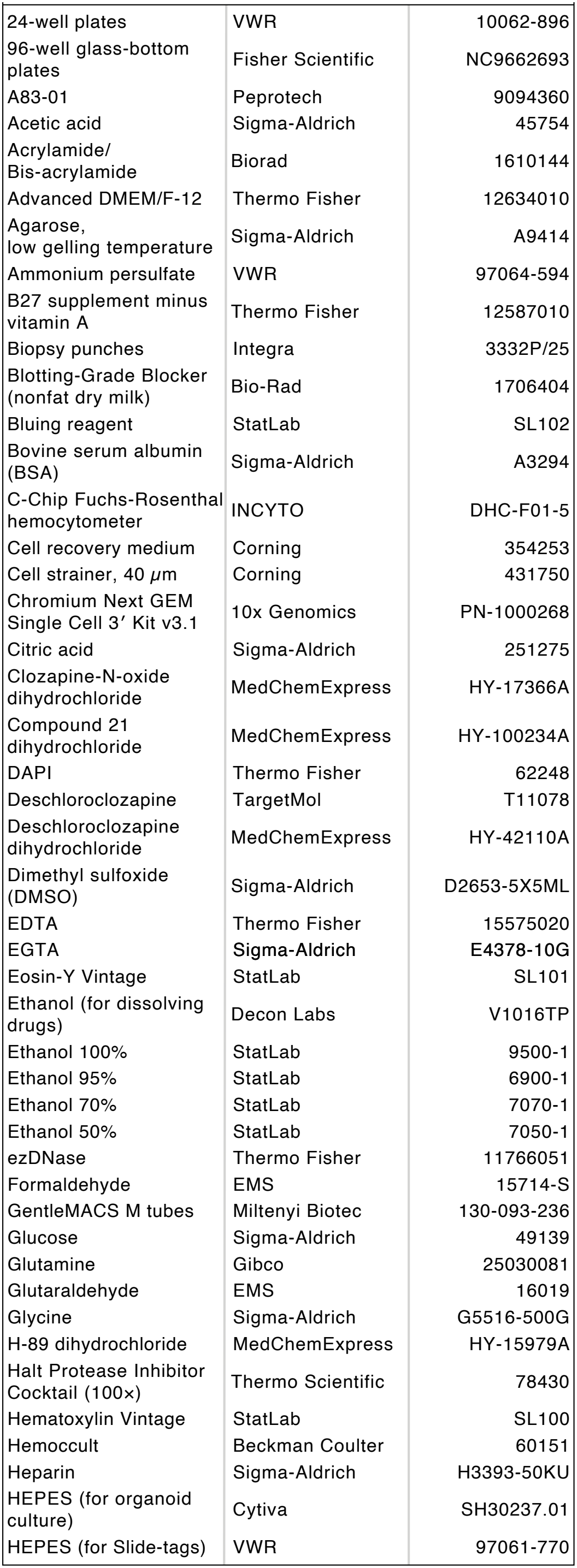
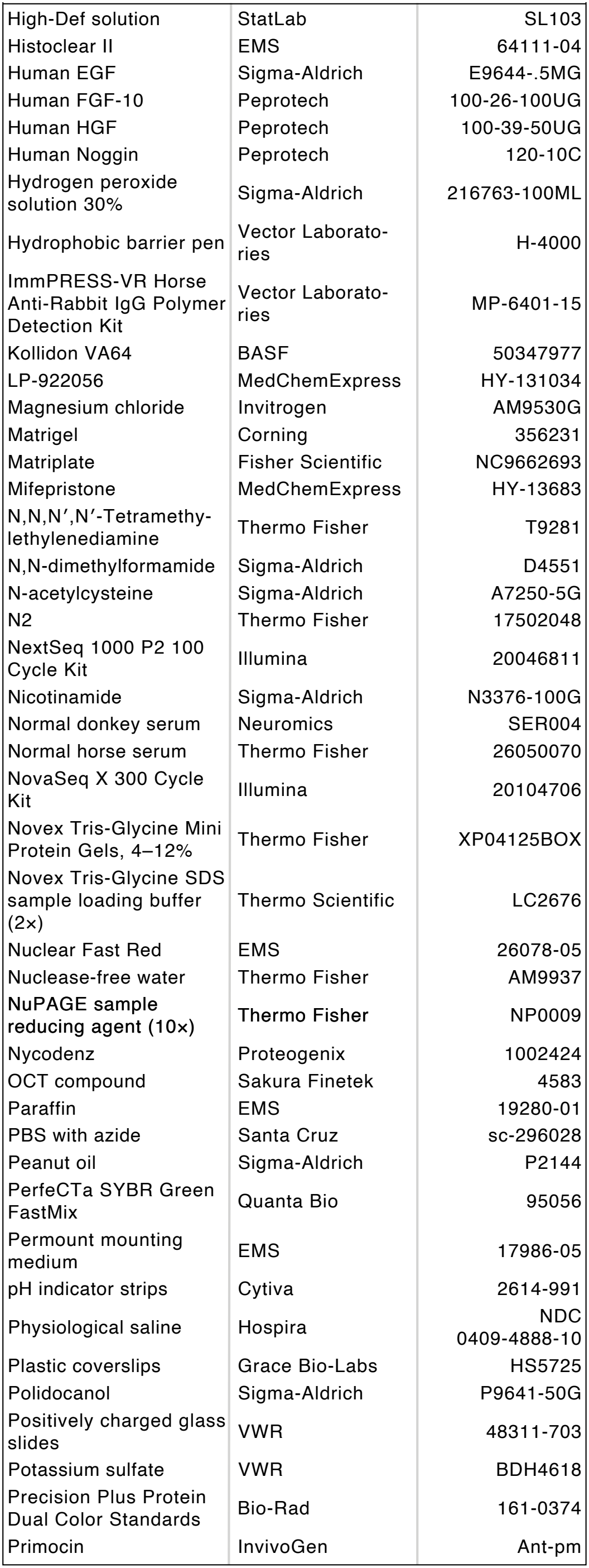
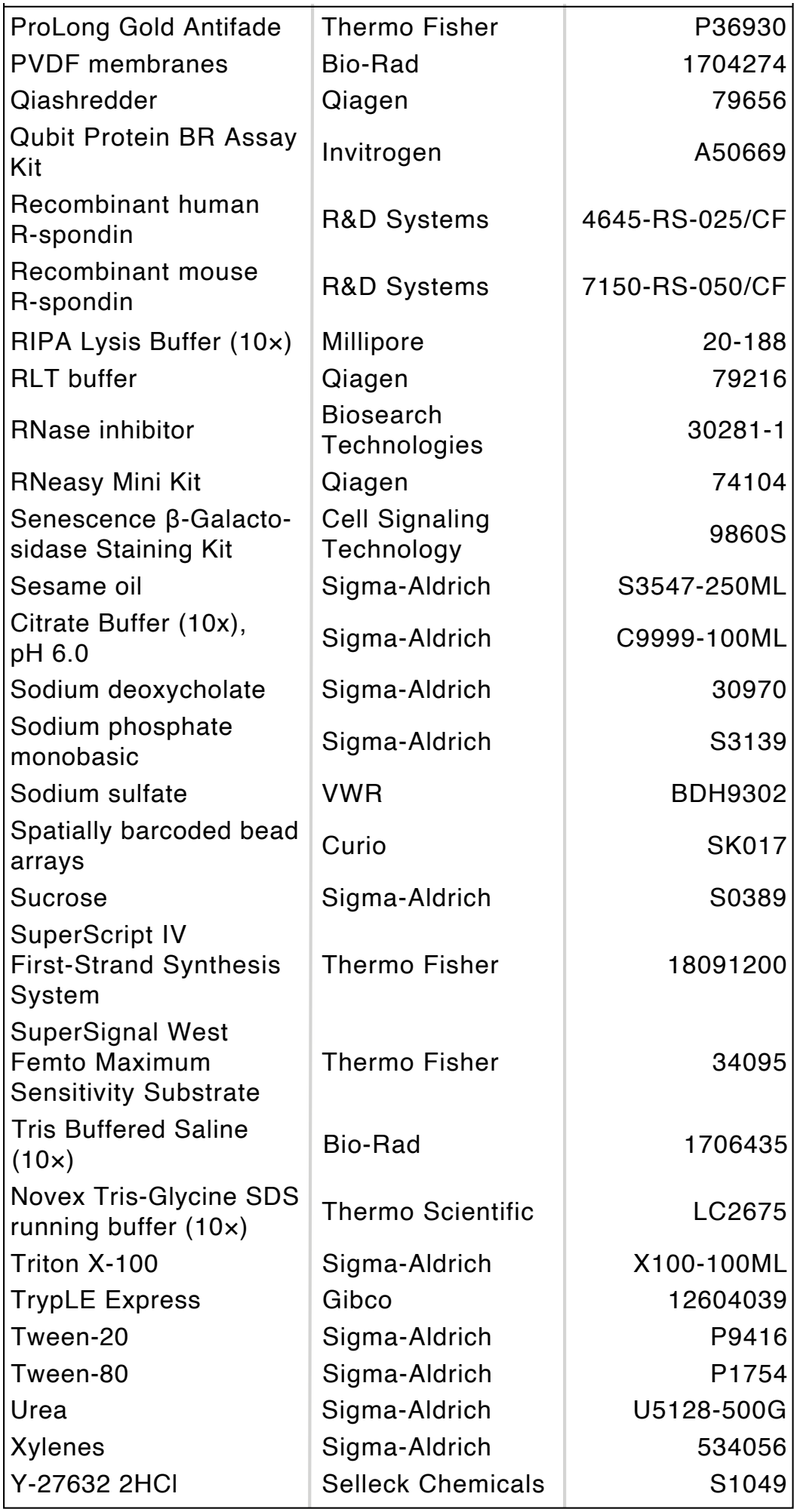
Materials used in this study.

## 1. Animal husbandry

All mice were housed under a 14:10 hour light–dark cycle at 21–25 °C and 30–70% humidity, with ad libitum access to food (irradiated LabDiet Prolab Isopro RMH 3000 5P75; LabDiet, St. Louis, MO) and water. Up to five animals were group-housed per cage. Cages were either polysulfone (dimensions 7.75” W × 12” L × 6.5” H; Allen-town, Allentown, NJ) or PET (dimensions 9.2” W x 14.7” L x 5.5” H; Innovive, San Diego, CA). Bedding was either coarse aspen Sani-chip bedding (PJ Murphy, Ladysmith, WI) or Alpha-Dri (Shepherd Specialty Papers, Watertown TN). All procedures were approved by the Institutional Animal Care and Use Committee of Harvard University and conducted in accordance with relevant ethical regulations and guidelines. Mouse strains used in this study are listed in Table S2. At least three animals were used for each condition unless otherwise noted.

## 2. Induction of decidualization and menstruation in X-Mens mice

### 2.1. Induction of pseudopregnancy

Female mice were paired with vasectomized B6 or CD-1 males at a ratio of one or two females per male after 4 PM. Copulatory plugs were checked the following morning before noon. The day a plug was detected was designated as day 0.5 of pseudopregnancy, after which plugged females were separated from males.

### 2.2. Induction of decidualization and menstruation

On the evening of pseudopregnancy day 3.5 (after 4:00 PM), mice received 48 μg/mL clozapine-N-oxide dihydro-chloride (CNO) in their drinking water. On pseudopregnancy day 4.5, animals received five consecutive intraperitoneal injections of a DREADD agonist cocktail (24 μg/mL CNO, 125 μg/mL deschloroclozapine dihydro-chloride, and 200 μg/mL compound 21 dihydrochloride in physiological saline) at two-hour intervals, starting between 8 and 10am. The day of the injections (pseudo-pregnancy day 4.5) was designated as 0.5 dpi (days post-induction). The following morning (pseudopregnancy day 5.5), CNO-supplemented water was replaced with regular drinking water prior to noon. Control animals received plain drinking water without CNO and were administered intraperitoneal injections of physiological saline on the same schedule as experimental animals. For the 1.5 dpi early timepoint, the animal was not given CNO in the drinking water and was induced only with intraperitoneal injections.

Alternatively, animals received a single subcutaneous injection of deschloroclozapine (2 mg/kg) in 5% ethanol and 95% sesame oil on the evening of pseudopregnancy day 3.5. This method is only used for experiments presented in Figure S7 and Figure 6F.

### 2.3. Assessment of menstruation

Menstruation was monitored by collecting vaginal lavage samples beginning the day after DREADD agonist injections. Animals were gently restrained, and 30–100 μL of physiological saline was pipetted into the vagina and aspirated 5–10 times. A few microliters of the lavage sample were tested for the presence of red blood cells using a fecal occult blood test (Hemoccult). The first day blood was detected in the lavage was designated as bleeding day 0.5. Daily lavages continued until blood was no longer detected or the uterus was collected. For non-menstruating animals, including controls, vaginal lavages were continued daily through pseudopregnancy day 11.5.

**Table S2:** Mouse strains used in this study.

| **Strain** | **Full Name** | **MGI ID** | **Strain No** | **Reference** | **Source** |
|---|---|---|---|---|---|
| B6 | C57BL/6J | – | Jax: 000664 | – | Jax |
| CD-1 | CD-1 IGS | – | Charles River: 022 | – | Charles River |
| *Amhr2<sup>Cre</sup>* | Amhr2<sup>tm3(cre)Bhr</sup> | 3042214 | – | (Jamin et al., 2002) | Ron Chandler |
| *Ltf<sup>iCre</sup>* | Ltf<sup>tm1(icre)Tdku/J</sup> | 5581966 | Jax: 026030 | (Daikoku et al., 2014) | Ron Chandler |
| *R26<sup>GsD</sup>* | Gt(ROSA)26Sor<sup>tm1(CAG-Chrm3*/GFP,cAMPRE-luc)Berd</sup> | 5696731 | – | (Akhmedov et al., 2017) | Bruce Morgan |
| *Tg<sup>GqD</sup>* | B6N;129-Tg(CAG-CHRM3*,-mCitrine)1Ute/J | 5614045 | Jax: 026220 | (Zhu et al., 2016) | Jax |
| *R26<sup>mTmG</sup>* | B6.129(Cg)-Gt(ROSA)26Sor<sup>tm4(ACTB-tdTomato,-EGFP)Luo</sup>/J | 3803814 | Jax: 007676 | (Muzumdar et al., 2007) | Jax |
| *R26<sup>GiD</sup>* | B6.129-Gt(ROSA)26Sor<sup>tm1(CAG-CHRM4*,-mCitrine)Ute</sup>/J | 5614050 | Jax: 026219 | (Zhu et al., 2016) | Jax |

### 2.4. Repeated menstruation

After vaginal bleeding was absent for at least two consecutive days, animals were re-paired with vasectomized males to induce new cycles of pseudopregnancy, decidualization induction, and menstruation assessment.

### 2.5. Mifepristone administration

On the evening of pseudopregnancy day 6.5 (after 4 PM), animals received a subcutaneous injection of 20 mg/kg mifepristone dissolved in 5% ethanol and 95% peanut oil. Following mifepristone administration, menstruation was monitored daily by vaginal lavage as described above. Animals that were already exhibiting vaginal bleeding at the time of mifepristone injection were excluded from analysis.

## 3. Sample collection

### 3.1. Mouse

Mice were euthanized either by exposure to CO_2_ gas for 5 minutes or by cervical dislocation; for Slide-tags and RNA extraction experiments, only cervical dislocation was used. Following euthanasia, the abdominal skin and muscle layers were incised, and the reproductive tract was exposed. The uteri were excised by cutting the ovarian fat pads, mesometrium, and cervix. The mesometrium and ovarian fat pads were trimmed as needed.

Excised uteri were processed in three ways:

1. Imaging and fixation: immediately after dissection, uteri were arranged on filter paper rounds placed inside a 10” × 10” LED photo box under uniform lighting and photographed for documentation. Following imaging, tissues were fixed overnight at 4 °C in 4% formaldehyde in PBS with gentle agitation. Samples were washed three times in PBS (each wash lasting 15 minutes to 2 hours at room temperature, with gentle agitation) and then stored in PBS containing 0.1% sodium azide at 4 °C with gentle agitation.
2. RNA extraction: after dissection, each uterine horn (between the oviduct and the uterine body) was placed individually into sterile 1.5 mL microcentrifuge tubes, and snap frozen in liquid nitrogen. Samples were stored at –80 °C until use.
3. Slide-tags experiments: uterine horns were flash frozen in optimal cutting temperature (OCT) compound and stored at –80 °C until use.

### 3.2. Human

Human uteri were procured from deceased organ donors through the National Disease Research Interchange (NDRI). The protocol was deemed Not Human Subjects Research by the Harvard Institutional Review Board (IRB21-0272). Donors who had received chemotherapy or radiation within the past year, or who had a diagnosis of diabetes, were excluded. Whole uteri were recovered and submerged in chilled University of Wisconsin/SPS-1 solution or histidine-tryptophan-keto-glutarate (HTK) solution, packaged on wet ice, and maintained at 4 °C until shipment. Donor characteristics are listed in Table S3.

The uterus was oriented with the fundus superiorly and the cervix inferiorly, and the cervix was removed. The uterine body was serially sectioned perpendicular to the cervix–fundus axis into sections of approximately 0.5–1 cm with a trimming knife. A central section was selected and bisected along the sagittal midline to generate left and right halves. Each half was submerged in OCT compound and flash frozen. Samples had no visible fibroids or serosal endometriosis on inspection upon receipt, and no adenomyosis on histology. The two samples were staged by a pathologist as secretory and menstrual. Single-nucleus transcriptomes of the secretory sample contained mature decidual cells expressing *IGFBP1* and *PRL*, consistent with the beginning of the late secretory phase.

## 4. In vivo manipulations

### 4.1. Chemical ablation of the luminal epithelium

Polidocanol foam was generated by combining four parts air with one part 5% (w/v) polidocanol in syringes joined by a luer lock adapter and passing the mixture through the adapter 60 times in each direction, as described previously (Ang et al., 2026). On pseudopregnancy day 3.5, polidocanol foam was injected into the uterine lumen from the cervical third of the horn using a 32G needle until the entire horn was visibly filled, and the procedure was repeated for the contralateral horn. Foam that leaked from the injection site was flushed away with sterile saline. The uterus was returned to the abdominal cavity, and the incision was closed with the foam left in place. Four hours after polidocanol injection, animals received a single subcutaneous injection of deschloroclozapine (2 mg/kg) in 5% ethanol and 95% sesame oil (n=3). Control animals received an equivalent volume of vehicle alone (n=3).

### 4.2. Notum inhibition

The Notum inhibitor LP-922056 (Brommage et al., 2019) was dissolved in DMSO at 50 mg/mL and diluted to 2 mg/mL in physiological saline containing 0.1% Tween-80 (final DMSO concentration 4%). GsD X-Mens animals were induced as described above. At 1.5 dpi, animals received two intraperitoneal injections of LP-922056 at 10 mg/kg (5 µL/g body weight), the first between 9:00 and 9:30 AM and the second between 1:30 and 2:00 PM (n=3). Control animals received vehicle (4% DMSO and 0.1% Tween-80 in saline) on the same schedule (n=3). Mice were euthanized by cervical dislocation between 4:30 and 5:00 PM on 2.5 dpi, and uteri were fixed, cryoprotected in 30% sucrose, and embedded in OCT (see Harvesting Uteri, approach 1, and Cryo-sectioning, staining, and imaging).

**Table S3:**
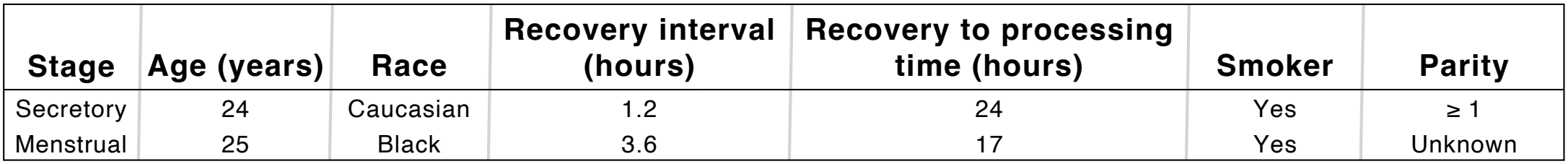
Uterus donors.

| **Stage** | **Age (years)** | **Race** | **Recovery interval (hours)** | **Recovery to processing time (hours)** | **Smoker** | **Parity** |
|---|---|---|---|---|---|---|
| Secretory | 24 | Caucasian | 1.2 | 24 | Yes | ≥ 1 |
| Menstrual | 25 | Black | 3.6 | 17 | Yes | Unknown |

### 4.3. H-89 treatment

The PKA inhibitor H-89 was dissolved in DMSO at 50 mg/mL and diluted to 4 mg/mL in sesame oil (final DMSO concentration 8%). B6 females were paired with intact B6 males after 4 PM, and copulatory plugs were checked the following morning before noon. The day a plug was detected was designated as gestational day 0.5 (GD0.5), after which plugged females were separated from males. Animals received two subcutaneous injections of H-89 at 20 mg/kg (5 µL/g body weight), the first on the evening of GD3.5 between 6:30 and 7:30 PM and the second on the morning of GD4.5 between 9:00 and 10:00 AM. Control animals received an equivalent volume of vehicle (8% DMSO in sesame oil) on the same schedule. Mice were euthanized on the evening of GD6.5 and uteri were excised (see Harvesting Uteri, approach 1) and the number of implantation sites, scored as discrete decidual swellings, was counted across both uterine horns for each animal.

### 4.4. GiD activation during early pregnancy

*Amhr2*^*Cre*^ x *R26*^*GiD*^ and *Ltf* ^*iCre*^ x *R26*^*GiD*^ females were mated with intact B6 males after 4 PM, and copulatory plugs were checked the following morning before noon. The day a plug was detected was designated as gestational day 0.5 (GD0.5), after which plugged females were separated from males. Animals received CNO in their drinking water on the evening of GD3.5 (after 4:00 PM) and, on GD4.5, five consecutive intraperitoneal injections of the DREADD agonist cocktail at two-hour intervals, at the concentrations and on the schedule described for X-Mens induction (see Induction of decidualization and menstruation in X-Mens mice), with gestational days in place of pseudo-pregnancy days. CNO-supplemented water was replaced with regular drinking water on the morning of GD5.5, prior to noon. Control animals received plain drinking water without CNO and intraperitoneal injections of physiological saline on the same schedule. Mice were euthanized on GD6.5 and uteri were excised (see Harvesting Uteri, approach 1) and the number of implantation sites, scored as discrete decidual swellings, was counted across both uterine horns for each animal.

## 5. Organoid culture from menstrual effluent

### 5.1 Sample collection

Menstrual effluent was collected from X-Mens animals that underwent induction as described above and received mifepristone injections to synchronize bleeding (Figure S11A). Organoids were generated from pooled uterine lavage samples collected from pairs of females with copulatory plugs on the same day receiving identical treatments. Uterine lavages were collected at two time points from each animal: (1) pseudopregnancy day 3.5 (prior to agonist administration) and (2) pseudopregnancy day 7.5 (the day after mifepristone administration). For each time point, each mouse was lavaged with 100 µl of sterile saline; lavage fluids from each pair were pooled in transport medium consisting of Advanced DMEM, 10% fetal bovine serum, 10 mM HEPES, and 0.1 mg/mL primocin, and transported on ice.

### 5.2 Establishing organoid culture

Cells were pelleted by centrifugation at 228 x g for 5 min at 4 °C. The pellet was resuspended in 150–250 µl of Matrigel, and 50 µl droplets were plated into individual wells of a 24-well plate. Plates were incubated at 37 °C for 10 min to solidify the Matrigel, after which 750 µl warm endometrial organoid medium (Advanced DMEM supplemented with 0.05 µg/mL human EGF, 0.1 µg/mL human Noggin, 0.1 µg/mL recombinant human or mouse R-spondin, 0.1 µg/mL human FGF, 0.05 µg/mL human HGF, 1:100 N2 supplement, 1:50 B27 supplement minus vitamin A, 0.1 mg/mL Primocin, 1 µM nicotinamide, 2 mM glutamine, 0.5 µM A83-01, 1.25 mM N-acetylcysteine, and 10 µM Y27632), was gently added to each well. Plates were then incubated at 37 °C, 5% CO_2_.

### 5.3 Organoid maintenance

Organoids were maintained at 37 °C in a 5% CO_2_ humidified incubator. Media were replaced every 5 days by removing spent medium and adding 750 µl of fresh media. Organoid medium was prepared in bulk and stored at 4 °C; Y27632 was added freshly immediately before medium changes. The number of organoids per sample was counted manually between days 10–14 after initial plating. Subsequently, media were refreshed every 1–7 days, and organoids were passaged as required.

### 5.4 Organoid passaging

Cultures were harvested by gently washing Matrigel droplets in PBS and releasing organoids with TrypLE Express. The mixture was incubated for 5–10 min at 37 °C and mechanically dissociated by pipetting with P1000 and P200 tips, shearing against the well bottom. Organoid fragments were transferred to 10 ml Advanced DMEM plus 10% FBS and centrifuged at 228 x g for 5 min. The pellet was resuspended in Matrigel and replated in 50 µl droplets in a 24-well plate (1 droplet per well). The plate was incubated at 37 °C for 10 min; 500–750 µl of fresh endometrial organoid medium was added to each well. Standard passaging expanded one Matrigel droplet to four new droplets.

### 5.5 Organoid staining

Matrigel-embedded organoids were released using 1 ml Cell Recovery Medium, incubated on ice for 20 min, and washed with 9 ml Advanced DMEM by centrifugation at 228 x g. Pellets were resuspended in a 1:1 mix of endometrial organoid medium and Matrigel and plated as 50 µl droplets into 96-well glass-bottom plates. Plates were incubated at 37 °C for 10 min, then 500–750 µl endometrial organoid medium was added. After overnight incubation, organoids were fixed in 4% formaldehyde in PBS for 1 h at room temperature, blocked with 3% BSA in PBS with 0.1% Triton X-100, and incubated with primary antibodies overnight at 4 °C. Secondary antibodies were applied for at least 2 h at room temperature. (For details of antibodies and imaging, see below.)

## 6. Tissue processing, imaging, and image analysis

### 6.1. Paraffin sectioning and Hematoxylin & Eosin (H&E) staining

Following fixation and PBS washes, uteri were transferred to 70% ethanol and stored at 4 °C with gentle agitation for at least 24 hours. Tissue dehydration and clearing were performed through a graded ethanol series, followed by xylene and paraffin infiltration. Specifically, samples were placed sequentially in 95% ethanol for 45 minutes and then 30 minutes, 100% ethanol for 45 minutes and then 30 minutes, xylene for 45, 60, and 30 minutes, and finally three changes of paraffin for 60 minutes each. After processing, samples were embedded in paraffin molds and sectioned at 5 μm using a microtome. Sections were mounted onto positively charged glass slides for staining.

For staining, paraffin sections were first deparaffinized in three successive five-minute washes of Histoclear II, followed by rehydration through decreasing concentrations of ethanol and water. Specifically, slides were washed twice in 100% ethanol for 1 minute each, then in 95% ethanol for 30 seconds, 50% ethanol for 30 seconds, and twice in deionized water for 1 minute each. Slides were then stained with hematoxylin for 1 minute, rinsed three times in water for 30 seconds each, and treated with High-def solution for 30 seconds. After a further three water washes of 20 seconds each, slides were placed in bluing reagent for 1 minute, rinsed again three times in water for 20 seconds each, washed in 95% ethanol for 30 seconds, stained with eosin for 45 seconds, dehydrated in three 1-minute changes of 100% ethanol, and cleared in three 1-minute washes of Histoclear II. Stained slides were temporarily stored in Histoclear II prior to mounting. Slides were mounted using approximately 100 μL Permount mounting medium and immediately coverslipped, then allowed to cure for at least 24 hours before imaging.

### 6.2. Immunohistochemistry (IHC)

Paraffin sections were deparaffinized in three successive washes of Histoclear II, each lasting 5 minutes, and rehydrated through a graded ethanol series: twice in 100% ethanol for 3 minutes each, then 70% ethanol for 3 minutes, 50% ethanol for 3 minutes, and twice in water for 3 minutes each. *6*.*2*.*1*.

#### 6.2.1. Antigen retrieval

10 mM sodium citrate buffer (pH 6.0) containing 0.05% Tween-20 was preheated for 30 minutes in a glass Coplin jar placed inside a pressure cooker. Slides were then transferred into the preheated jar and boiled for 20 minutes. The jar was subsequently removed and allowed to cool on the bench for 30 minutes at room temperature. Slides were washed twice in PBS for 3 minutes each.

#### 6.2.2. Quenching and blocking

Endogenous peroxidase activity was quenched by incubation in PBS containing 3% hydrogen peroxide for 10 minutes at room temperature, followed by two 3-minute washes in PBS. Slide perimeters were marked with a hydrophobic barrier pen, and non-specific binding was blocked for 30 minutes at room temperature in a humid chamber using blocking buffer composed of PBS with 5% normal horse serum, 0.5% bovine serum albumin (BSA), 0.1% Triton X-100, and 0.1% sodium deoxycholate.

#### 6.2.3. Antibody incubation and detection

Primary antibodies diluted in blocking buffer composed of PBS with 5% normal horse serum, 0.5% bovine serum albumin (BSA), 0.1% Triton X-100, and 0.1% sodium deoxycholate, were applied to the sections and incubated over-night at 4 °C in a humid chamber. Slides were washed four times for 5 minutes each in PBS containing 0.1% Tween-20. Sections were then incubated for 30 minutes at room temperature in a humid chamber with the ready-to-use ImmPRESS-VR Horse Anti-Rabbit IgG Polymer Detection Kit (Vector Laboratories), after which slides were washed four times for 5 minutes each in PBT. Antibodies used in this study are listed in Table S4. Signal was developed with 3,3’Diaminobenzidine (DAB) substrate under visual inspection on a stereomicroscope, with development times of approximately 5 minutes. A slide processed in parallel without primary antibody was used to monitor background development and to determine the endpoint of the reaction.

#### 6.2.4. Counterstaining and mounting

Following development, slides were rinsed three times in water for 30 seconds each, counterstained with hematoxylin for 1 minute, rinsed three times in water for 30 seconds each, and treated with High-Def solution for 30 seconds. Slides were then placed in bluing reagent for 1 minute, rinsed three times in water for 20 seconds each, and dehydrated through 70% ethanol for 30 seconds, 95% ethanol for 30 seconds, and two changes of 100% ethanol for 30 seconds each. Sections were cleared in three 1-minute washes of Histoclear II, mounted using approximately 100 μL Permount mounting medium, immediately covers-lipped, and allowed to cure overnight before imaging.

### 6.3. Cryosectioning

The fixed tissues (see Harvesting Uteri, approach 1) were PBS-washed and incubated overnight in 30% sucrose in PBS at 4 °C for cryoprotection. Following equilibration, tissues were embedded in OCT compound and snap frozen. OCT blocks were stored at –80 °C until sectioning.

For Slide-tags experiments, sections were generated from uteri fresh-frozen in OCT (see Harvesting Uteri, approach 3). One section was used for nuclei isolation and sequencing; a consecutive section was used for immunofluorescence. See Slide-tags procedure.

Prior to sectioning, OCT blocks were equilibrated within the cryostat chamber at –19 to –21 °C for at least 30 minutes. Tissues were then sectioned at a thickness of 20-30 μm with a Leica CM 1860 cryostat, and the resulting sections were mounted onto positively charged glass slides. Prepared slides were stored at –80 °C until further use.

**Table S4:** Antibodies used in this study.

| Antibody | Host organism | Vendor | Catalog no | RRID | Dilution |
|---|---|---|---|---|---|
| CA9 | rabbit | Abcam | ab184006 | AB_3101886 | 1:1250 |
| CC3 | rabbit | Cell Signaling Technologies | 9661 | AB_2341188 | 1:125 |
| CD31 | goat | R&D Systems | AF3628 | AB_2161028 | 1:100 |
| CD45 | goat | R&D Systems | AF114 | AB_442146 | 1:1000 |
| CD64 | rabbit | Invitrogen | MA5-29706 | AB_2785530 | 1:500 |
| CDH1 | rat | Invitrogen | 13-1900 | AB_2533005 | 1:1000 |
| CDH3 | goat | R&D Systems | AF761 | AB_355581 | 1:200 |
| CK8 | rat | DSHB | TROMA-I | AB_2891089 | 1:20 |
| FOXA2 | rabbit | Cell Signaling Technologies | 8186 | AB_10891055 | 1:200 |
| GAPDH | rabbit | Cell Signaling Technologies | 2118 | AB_561053 | 1:10,000 |
| GFP | chicken | Abcam | ab13970 | AB_300798 | 1:1000 |
| GJA1 | rabbit | Abcam | ab11370 | AB_297976 | 1:100 |
| GPX3 | rabbit | Abcam | ab256470 | AB_3718666 | 1:150 |
| HIF1A | rabbit | Cell Signaling Technologies | 36169 | AB_2799095 | 1:2000 |
| JAG1 | goat | R&D Systems | AF599 | AB_2128257 | 1:500 |
| PDGFRA | rabbit | Abcam | ab203491 | AB_3645435 | 1:200 |
| pH3 | rat | BD Biosciences | 558217 | AB_397065 | 1:100 |
| SMA | mouse | Sigma-Aldrich | C6198 | AB_476856 | 1:1000 |
| tdTomato | goat | Origene | AB8181-200 | AB_3206272 | 1:1000 |

### 6.4. Immunofluorescence staining

Cryosections were brought to 4 °C for 10 minutes to equilibrate in a humid chamber. Fresh frozen sections were fixed in 4% formal-dehyde in PBS for 15 minutes at room temperature, then washed three times for 15 minutes each in PBS containing 0.1% Tween-20 (PBT). For all samples, non-specific binding was blocked for 30–60 minutes at room temperature using blocking buffer composed of PBS with 5% normal donkey serum, 0.5% bovine serum albumin (BSA), 0.1% Triton X-100, and 0.1% sodium deoxycholate. After blocking, slide perimeters were marked with a hydrophobic barrier pen, and primary antibodies diluted in blocking buffer were applied to the tissue sections (approximately 150–300 μl per slide, depending on section size and the number of sections per slide). Slides were incubated with primary antibody either overnight at 4 °C or for 2 hours at room temperature in a humid chamber. After incubation, slides were washed three times for 15 minutes each in PBT. Secondary antibodies, also diluted in blocking buffer, were then applied at the same volume as the primaries, and slides were incubated either overnight at 4 °C or for 2 hours at room temperature in a humid chamber. DAPI (1 µg/ml final concentration) was included in the secondary antibody solution to serve as a nuclear counterstain. After completion of the staining protocol, slides were washed three times for 15 minutes each in PBT. Slides were then mounted using ProLong Gold Antifade mounting medium, coverslipped, and allowed to cure for at least 24 hours at room temperature prior to imaging. Antibodies used in this study are listed in Table S4.

### 6.5. Hybridization Chain Reaction Fluorescence In Situ Hybridization (HCR-FISH)

HCR-FISH was performed with HCR™ Gold RNA-FISH kits (HCR™ HiFi Probes, HCR™ Gold Amplifiers, and the corresponding hybridization, probe wash, amplifier and amplifier wash buffers; Molecular Instruments, Inc., Los Angeles, CA), following the manufacturer’s HCR™ Gold RNA-FISH Protocol: Sample on a Slide and Sample Preparation Protocol for Fixed/Frozen Tissue Sections on a Slide (both version 2026-03-18).

Flash-frozen uterine horns (see Harvesting Uteri, approach 3) were cryosectioned at 20 µm onto positively charged slides and stored at –80 °C. Sections were fixed in ice-cold 4% paraformaldehyde in PBS for 15 min at 4 °C, dehydrated through 50%, 70% and two changes of 100% ethanol (5 min each at room temperature), air-dried, and encircled with a hydrophobic barrier pen before two washes in 1× PBS.

Sections were pre-hybridized in 200 µl HCR™ HiFi Probe Hybridization Buffer for 10 min at 37 °C in a humidified chamber. Probe solution was prepared by adding 2 µl of each HCR™ HiFi Probe to 100 µl of pre-warmed hybridization buffer, applied to the sections (50–100 µl), covered with a plastic coverslip, and incubated at 37 °C overnight. Coverslips were floated off in 1× HCR™ HiFi Probe Wash Buffer at 37 °C and excess probe removed by four 15-min immersions in the same buffer at 37 °C. Sections were pre-amplified in 200 µl HCR™ Gold Amplifier Buffer for 30 min at room temperature. Hairpins h1 and h2 (2 µl each) were snap-cooled separately at 95 °C for 90 s and cooled to room temperature in the dark for 30 min, then combined in 100 µl of HCR™ Gold Amplifier Buffer. Amplifier solution (50–100 µl) was applied, covered with a plastic coverslip, and incubated at room temperature for overnight in the dark. Coverslips were floated off in 1× HCR™ Gold Amplifier Wash Buffer and excess amplifier removed by four 15-min immersions in the same buffer at room temperature in the dark. Sections were mounted in Prolong Gold mounting medium and imaged after 24 h.

Amplifiers were assigned as follows: *Gpx3*, amplifier X3 (Alexa 647, displayed in magenta); *Bmp2*, amplifier X5 (Alexa 405, cyan); *Prl8a2*, amplifier X1 (Alexa 546, yellow). Specificity was assessed using sections processed without probe sets. Imaging was performed as described in section 6.8.

### 6.6. Senescence-associated β-galactosidase (SA-βGal) staining

SA-βGal activity was detected in fresh-frozen cryosections (see Harvesting Uteri, approach 3) using a protocol modified from a previously described method (Idelfonso-García et al., 2024), with the staining solution prepared from a commercial kit according to the manufacturer’s instructions. Comparative experimental groups were stained in parallel using the same batch of all reagents to avoid inter-assay variation.

#### 6.6.1. Solutions

Citric acid–sodium phosphate buffer was prepared by combining 36.85 mL of 100 mM citric acid with 63.15 mL of 200 mM sodium phosphate mono-basic, adjusted to exactly pH 6.0. Fixative solution consisted of 0.5% glutaraldehyde, 1.25 mM EGTA, and 2 mM MgCl_2_ in citric acid–sodium phosphate buffer. β-Galactosidase staining solution was prepared immediately before use from the Senescence β-Galactosidase Staining Kit (Cell Signaling Technology) by combining, per 1 mL, 930 μL 1× staining solution, 10 μL 100× solution A, 10 μL 100× solution B, and 50 μL of 20 mg/mL X-gal in N,N-dimethylformamide; the final pH was confirmed to be 6.0 using pH indicator strips.

#### 6.6.2. Staining

Slides were transferred from −80 °C and equilibrated at 4 °C for 10 minutes in a humid chamber. Sections were fixed for 5 minutes in fixative solution and washed three times for 5 minutes each in PBS. Slide perimeters were marked with a hydrophobic barrier pen, and approximately 250 μL of β-galactosidase staining solution was applied per slide, with the volume adjusted according to section area. The chamber was sealed to prevent evaporation, shielded from light with aluminum foil, and incubated at 37 °C in a dry incubator without CO_2_ for 48 hours, with color development monitored periodically on a stereomicroscope.

#### 6.6.3. Counterstaining and mounting

Sections were washed three times for 2 minutes each in PBS, counter-stained with Nuclear Fast Red for 30–60 seconds, and washed three times for 5 minutes each in PBS. Sections were mounted with Permount mounting medium and allowed to cure overnight before imaging.

### 6.7. Tissue clearing and immunostaining of thick sections

Fixed uterine horns (see Harvesting Uteri, approach 1) were mounted in 2–5% low gelling temperature agarose and sectioned at 200 μm thickness using a compresstome, as described previously (Ang et al., 2026). Sections were stored in PBS containing 0.1% sodium azide at 4 °C until use. Unless otherwise noted, all subsequent steps were carried out at room temperature.

#### 6.7.1. Permeabilization and bleaching

Sections were incubated in permeabilization solution 1 (0.2% Triton X-100, 10% DMSO, 0.2% sodium deoxycholate, 20 EDTA in PBS, pH 8.0) for 24 hours at 37 °C, then transferred to permeabilization solution 2 (0.2% Triton X-100, 10% DMSO, 100 mM glycine in PBS) for 1 hour at 37 °C. Because X-Mens tissues carry endogenous fluorescent proteins, sections were subsequently bleached in 10% hydrogen peroxide in 0.05 M phosphate buffer for 3 hours at 37 °C, washed twice briefly in PTwH (0.2% Tween-20, 10 μg/mL heparin in PBS), and washed three further times in PTwH for 10 minutes each.

#### 6.7.2. Immunostaining

Sections were blocked for 1 hour in PBS containing 0.2% Triton X-100, 10% DMSO, and 5% normal donkey serum, then incubated overnight at 4 °C with primary antibodies diluted in staining solution (0.2% Tween-20, 10 μg/mL heparin, 5% DMSO, and 5% normal donkey serum in PBS). Sections were washed twice briefly in PTwH, followed by three washes of 10 minutes each, and then incubated for 3 hours with donkey secondary antibodies diluted in staining solution containing DAPI (1:1000) as a nuclear counterstain. Sections were washed twice briefly in PTwH, followed by three washes of 10 minutes each. Antibodies used in this study are listed in Table S4.

#### 6.7.3. Optical clearing and mounting

To facilitate imaging through the full section thickness, residual liquid was removed and sections were incubated for at least 1 hour in EZ index solution (80% (w/v) Nycodenz, 42% (w/v) urea, and 0.025% sodium azide in 0.2 M phosphate buffer, pH 7.4). Sections were mounted in EZ index on slides fitted with 200 μm spacers and imaged within 4 days of mounting.

### 6.8. Imaging

#### 6.8.1. Brightfield imaging

H&E, IHC and SA-βGal slides were imaged on a Zeiss Axioscan.Z1 slide scanner equipped with a Plan-Apochromat 20×/0.8 objective and a VIS LED light source under brightfield illumination. Image curation (resizing, cropping, and adjusting white balance and saturation) was performed in Fiji (Schindelin et al., 2012) as needed.

#### 6.8.2. Confocal imaging

Immunofluorescence samples, both cryosections and cleared thick sections, were imaged on a Yokogawa CSU-W1 spinning disk confocal system mounted on an inverted Nikon Ti-2E microscope, using either a CFI60 Plan Apochromat Lambda 20×/0.75 NA objective or a CFI60 Plan Apochromat Lambda D 10×/0.45 NA objective. Large fields were acquired using the Nikon Elements JOBS module, and stitching and maximum intensity projections were generated using the Nikon Elements General Analysis 3 module. Thick-section images are shown as maximum intensity projections. Acquired images were processed and analyzed using Fiji.

### 6.9. Segmentation and quantification of pH3 and CC3

Cryosections (20 µm) underwent immunofluorescence staining for pH3, CC3 and DAPI (see Cryo-sectioning, staining, and imaging). Samples were imaged with the 10× objective (pixel size 0.647 µm; z-step 5 µm) using identical acquisition settings for all samples. One section per stage was analyzed. Maximum intensity projections of the three channels were analyzed in raw intensity units with a custom Python script (Python 3.10.19; NumPy, SciPy, scikit-image, pandas and Cellpose; (Stringer et al., 2021)), applying one set of parameters to every image. The tissue was segmented from the DAPI channel by Gaussian smoothing (σ = 8 px, ≈5 µm), thresholding, filling holes, removing objects smaller than 1 mm^2^, and retaining the largest connected component. Nuclei were segmented from the DAPI channel with the pretrained Cellpose “nuclei” model ((Stringer et al., 2021); expected diameter 12 µm), restricted to the tissue mask, and objects outside 8–400 µm^2^ were discarded. For each of the pH3 and CC3 channels, the image was despeckled with a median filter (disk, radius 2 px), and a Gaussian-blurred copy (σ = 20 px, ≈13 µm) was subtracted to remove background. The result was thresholded (pH3 > 450; CC3 > 2,000). Connected objects were retained if they lay within the tissue mask, had an area of 5–200 µm^2^ and a circularity (4πA/P^2^) of at least 0.30. pH3 objects also had to lie within a segmented nucleus, and CC3 objects within 10 µm of one. Each retained object was counted as one positive cell and reported as a percentage of segmented nuclei and per mm^2^ of tissue. For display, the CC3 and pH3 masks were dilated with a disk of radius 6 px (≈3.9 µm); all counts and positions were calculated from the undilated objects.

The radial position of each object was its distance from the centroid of the tissue mask divided by the radius of a circle with the same area as the tissue (r_equiv), giving 0 at the section center and approximately 1 at the periphery. Radial distributions were visualized as Gaussian kernel density estimates (bandwidth factor 0.25). Original imaging files, segmentation parameters (parameters.-json), undilated masks and per-object coordinates will be available upon publication.

## 7. Molecular biology techniqu

### 7.1. Quantitative PCR analysis

#### 7.1.1. RNA extraction

Cryopreserved uterine horns were processed immediately without thawing by direct dissociation in 1 mL RLT buffer using a gentleMACS dissociator (Miltenyi) with gentleMACS M tubes, following the manufacturer’s instructions. 800 µl of the resulting tissue lysate was then transferred to a Qiashredder column and centrifuged at 17,000 g for 3 minutes at 4 °C to remove tissue debris. Total RNA was isolated from the cleared lysate using the Qiagen RNeasy Mini Kit according to the manufacturer’s protocol. RNA concentration was determined using a NanoDrop spectrophotometer, and RNA samples were stored at −80 °C until further use.

#### 7.1.2. cDNA synthesis

Up to 2 µg of total RNA from each sample was treated with ezDNase to remove genomic DNA contamination, per the manufacturer’s protocol. First-strand cDNA synthesis was performed using the SuperScript IV First-Strand Synthesis System with oligo(dT) primers, following the manufacturer’s instructions. Synthesized cDNA was stored at −20 °C prior to quantitative PCR analysis.

#### 7.1.3. Quantitative PCR (qPCR)

Reactions were prepared in a final volume of 20 µl containing 10 µl PerfeCTa SYBR Green FastMix, 0.5 µl cDNA, 1 µl primer mix (10 µM stock concentration of each primer), and 8.5 µl nuclease-free water. For each sample, qPCR was performed for three genes: two housekeeping genes (*Actb* and *Gapdh*) and the decidualization marker *Prl8a2*. Each gene was measured in triplicate as a technical control, and for each experimental condition, at least three independent uterine samples were analyzed as biological replicates. qPCR was performed on a Quant-Studio 7 real-time PCR system. Relative expression levels of *Prl8a2* were calculated using the ΔΔCt method, with normalization to the uninduced pseudopregnant day 4.5 samples. Primers used for quantitative PCR are listed in Table S5.

**Table S5:**
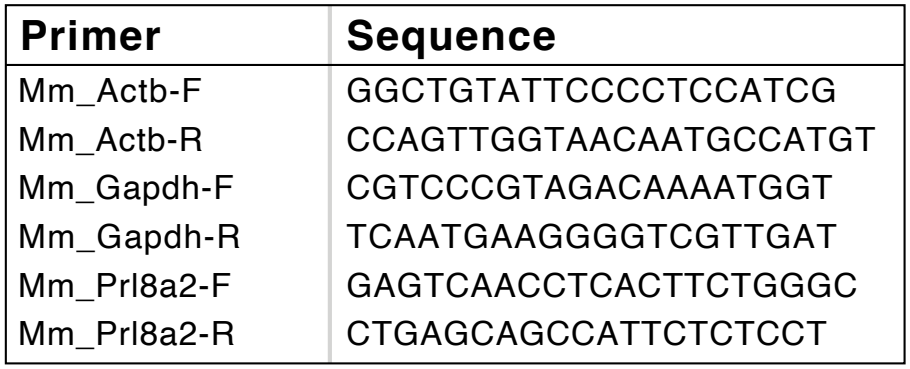
Primers used in this study.

### 7.2. Western Blotting

#### 7.2.1. Sample collection

Mice were euthanized by cervical dislocation and the uterus was harvested as described above (see Harvesting Uteri). Ovaries and adherent fat were removed from one uterine horn, which was then snap frozen in liquid nitrogen and stored at −80 °C until protein extraction.

#### 7.2.2. Protein extraction

Frozen uterine horns were kept on dry ice and weighed. Lysis buffer was prepared fresh in MilliQ water as 1× RIPA lysis buffer, and supplemented with 1× Halt Protease Inhibitor Cocktail and 1× EDTA, and was added at 1 mL per 30 mg of tissue. Tissue was placed in gentleMACS M tubes with cold lysis buffer and dissociated using a gentleMACS dissociator. Lysates were centrifuged at 5,000 g for 5 minutes at 4 °C, and aliquots of the cleared lysate were stored at −80 °C until use.

#### 7.2.3. SDS-PAGE

Protein concentrations were determined using the Qubit Protein Broad Range Assay Kit on a Qubit fluorometer. Each sample was prepared to contain 10 μg protein, 10 μL 2× Novex Tris-Glycine SDS sample loading buffer, and 2 μL 10× NuPAGE sample reducing agent in a final volume of 20 μL made up with MilliQ water, and denatured at 85 °C for 2 minutes. Each sample (18 μL), together with 4 μL Precision Plus Protein Dual Color Standards, was loaded onto Novex 4–12% Tris-Glycine mini protein gels and run at 150 V for 45 minutes in 1× Tris-Glycine running buffer.

#### 7.2.4. Immunoblotting

Proteins were transferred onto PVDF membranes using a Bio-Rad Trans-Blot Turbo Transfer System with the high molecular weight transfer program. Membranes were blocked for 1 hour at room temperature in TBST (1× Tris-buffered saline containing 0.1% Tween-20) supplemented with 5% blotting-grade nonfat dry milk. Membranes were then incubated over-night at 4 °C with primary antibodies against HIF1A and GAPDH, diluted in TBST containing 5% nonfat dry milk. Membranes were washed three times for 5 minutes each in TBST and once for 5 minutes in TBS, incubated for 1 hour at room temperature with HRP-conjugated goat anti-rabbit IgG (H+L) secondary antibody (1:20,000) diluted in TBST containing 5% nonfat dry milk, and washed again as described above. Antibodies used in this study are listed in Table S4.

#### 7.2.5. Detection

Blots were developed using SuperSig-nal West Femto Maximum Sensitivity Substrate for 5 minutes and imaged on a Bio-Rad ChemiDoc Touch imaging system. GAPDH signal was acquired using automatic rapid exposure and HIF1A signal using a 10-second exposure.

## 8. Slide-tags

### 8.1. Samples

We collected a time course of GsD X-Mens uteri (uninduced; 1.5 dpi early; 1.5 dpi late; 2.5 dpi; 3.5 dpi; 4.5 dpi bleeding). We confirmed the progression of decidualization in these samples based on the size of the CDH3 decidual region in immunostaining. One uterine cross-section from each time point was placed on a barcoded bead array, with the exception of the uninduced sample, for which two consecutive sections were placed on the array side-by-side. The mouse samples were processed across nine 10X lanes. For the human dataset, a section from each cycle stage (secretory and menstrual) was placed on a separate barcoded bead array, and samples were processed across three 10X lanes per section, for a total of six 10X lanes.

### 8.2. Slide-tags procedure

Slide-tags assays were performed largely as described in the original publication (Russell et al., 2024), with several modifications.

#### 8.2.1. Preparation of barcoded bead arrays

Barcoded bead arrays were manufactured and sequenced as previously described, using the 3′ feature barcode capture sequence ((TAGS beads): 5′-TTT-PC-GTGACTGGAGTTCAGACGTGTGCTCTTC CGATCTJJJJJJJJTCTTCAGCGTTCCCGAGAJJJJJJJN NNNNNNVVGCTTTAAGGCCGGTCCTAGCAA-3′)

In addition to custom-manufactured arrays, transferred Curio barcoded bead arrays were also used.

To minimize bead loss, beads were transferred from their original substrate to an acrylamide-coated glass coverslip. First, 25 mm square coverslips were thoroughly cleaned and incubated for 15 minutes at room temperature in a solution of 0.2% (v/v) bind silane (3-(trimethoxysilyl)propyl methacrylate) and 1% (v/v) acetic acid in ethanol. After treatment, coverslips were washed three times in 100% ethanol and allowed to dry completely.

A fresh acrylamide solution was prepared on ice consisting of 0.6% (v/v) tetramethylethylenediamine (TEMED), 0.06% (v/v) ammonium persulfate (APS), and 10% (v/v) Acrylamide/Bis-acrylamide (19:1) in water. Approximately 30 μL of this acrylamide solution was applied to the bind silane-treated side of each coverslip. The Curio barcoded array was then placed bead-side down onto the acrylamide solution, ensuring that no bubbles formed between the array and the coverslip. An untreated coverslip was immediately placed on top to generate a thin acrylamide layer. The assembly was incubated undisturbed for 45 minutes at room temperature to allow polymerization.

After incubation, assemblies were gently separated. Successful bead transfer was verified by visual inspection. Excess acrylamide at the coverslip edges was removed, and the coverslips were stored for at least 12 hours at room temperature to ensure the acrylamide layer was completely dry before further use.

#### 8.2.2. Tissue sectioning

Tissue samples were fresh frozen and then cryosectioned to 20 or 30 μm thickness on a Leica CM1950 cryostat at −18 to -21 °C. Excess OCT was removed from sections using biopsy punches before mounting tissue directly onto the barcoded array surface, taking care to avoid tissue folds. Except for the uninduced GsD X-Mens sample (in which two consecutive, small sections were placed side-by-side on a single array), only one cross-section was placed per array (for details, see Figure S12A). Sections were gently melted onto the arrays by warming with a finger placed beneath the barcoded array, then immediately transferred to a glass slide resting on ice. Each tissue section was then covered with 6–10 μL dissociation buffer (82 mM sodium sulfate, 30 mM potassium sulfate, 10 mM glucose, 10 mM HEPES, 5 mM magnesium chloride).

#### 8.2.3. Barcode release and nuclei isolation

Spatial barcode release was initiated by exposure of the barcoded bead array to ultraviolet (365 nm) light (0.42 mW mm^−2^, Thorlabs, M365LP1-C5, LEDD1B) for 1 minute, followed by incubation on ice for 14 minutes to facilitate diffusion of the spatial barcodes. The barcoded array was then transferred to a 12-well plate on ice, to which 2 mL of extraction buffer (dissociation buffer supplemented with 1% Kollidon VA64, 1% Triton X-100, 0.01% BSA, and 666 U mL^−1^ RNase inhibitor) was carefully added away from the tissue “puck.” Tissue was mechanically dissociated using a 200 μL pipette tip, performing 15–20 triturations as assessed by visual inspection, and the process was repeated until complete removal of tissue from the array. Further mechanical dissociation was performed with a 1000 μL pipette tip using 60–100 triturations, also monitored by visual inspection.

The resulting nuclei suspension was carefully aspirated and transferred, and the well was rinsed twice with 1 mL wash buffer (82 mM sodium sulfate, 30 mM potassium sulfate, 10 mM glucose, 10 mM HEPES, 5 mM magnesium chloride, 50 µL RNase inhibitor), and the rinses were pooled with the nuclei suspension to a final volume of 20 mL. Nuclei were pelleted by centrifugation at 600 g for 10 minutes at 4 °C in a pre-cooled swinging bucket centrifuge. After centrifugation, the supernatant was carefully removed, leaving 1 mL, and the pellet was resuspended and filtered through a pre-cooled, pre-wetted 40 μm cell strainer. This suspension was further centrifuged at 300 g for 10 minutes at 4 °C, the supernatant was removed to leave a final volume of 50 μL, and the pellet was resuspended. Nuclei were counted manually using a disposable hemocytometer.

#### 8.2.4. 10X Genomics library preparation

A total of 43.3 μL of counted nuclei was loaded into the 10x Genomics Chromium controller using the Chromium Next GEM Single Cell 3′ Kit v3.1. Subsequent steps were performed using the Chromium Next GEM Single Cell 3′ Reagent Kits v3.1 (Dual Index) with Feature Barcode Technology for Cell Surface Protein (Cell Ranger CG000317) according to the manufacturer’s protocol, with minor modifications. Spatial barcode libraries were prepared according to procedures for cell-surface protein library preparations, employing seven PCR cycles during the index PCR step. For experiments using Curio barcoded bead arrays, 1 μL of 0.329 μM spike-in primer (5′-GTGACTGGAGTTCAGACGT-3′) was included during cDNA amplification, and the Index TT plate was used for the index PCR step.

#### 8.2.5. Slide-tags sequencing

Single-cell RNA-seq and spatial barcode libraries were sequenced using either an Illumina NextSeq 1000 instrument with a P2 100 Cycle Kit or an Illumina NovaSeq X instrument with a 300 Cycle Kit.

#### 8.2.6. Slide-tags data preprocessing

All code for processing Slide-tags data is publicly available at https://github.com/thechenlab/Slide-tags. Binary base call (BCL) files were demultiplexed using 10x Genomics Cell Ranger mkfastq (version 8.0.1) (Zheng et al., 2017). Gene expression reads were mapped to a custom reference genome that included the sequences for the GsD and GqD transgenes. This reference was generated using Cell Ranger mkref (version 7.2.0) (Zheng et al., 2017), and gene expression matrices were produced using Cell Ranger count (version 7.2.0). The resulting gene expression matrices were subsequently processed using CellBender remove-background (version 0.2.2), with the --total-droplets-included parameter set to 50,000 to reduce contamination from ambient RNA.

Reads corresponding to spatial barcode libraries were processed with custom Julia scripts (version 1.10.3) (Bezanson et al., 2017). Puck coordinate files were either acquired from internal databases or obtained through the Curio Bioscience Bead Barcode Whitelist Retrieval tool (https://knowledgebase.curiobioscience.com/bioinformatics/tile-barcode/) and converted to comma-separated values format. The Julia scripts parsed raw FASTQ files and removed reads that contained an ‘N’ in the UMI sequence, did not have a detectable universal primer (UP) site (permitting one deletion or up to two mismatches), or had a spatial barcode that did not match the whitelist CSV (allowing one mismatch or deletion). The scripts produced a dictionary in which each entry was identified by a triplet consisting of cell barcode, UMI, and spatial barcode, with values corresponding to read counts.

These dictionaries were further processed in R (version 4.3.3) using custom R scripts. Reads were further filtered to exclude chimeric sequences, reads with low-quality spatial barcodes, and those that did not correspond to a called cell. The remaining reads were collapsed into UMI counts for each cell and spatial barcode, yielding a spatial barcode count matrix. To assign spatial positions to each nucleus, DBSCAN clustering was used to classify spatial barcodes as either signal or noise. Hyperparameters (ε and minPts) were automatically optimized to achieve maximal accuracy in cell placement. For cells with a single robust signal cluster, the centroid was calculated by weighting nUMI values, and this centroid was assigned as the spatial coordinate for the cell.

## 9. Slide-tags data analysis

### 9.1. Atlas construction

#### 9.1.1. Alignment, quantification and ambient RNA removal

Gene-expression reads were aligned and quantified with the 10x Genomics Cell Ranger count pipeline (version 7.2.0) with default parameters. Human libraries were aligned to the 10x Genomics GRCh38-2024-A reference. Mouse libraries were aligned to a custom reference built with Cell Ranger mkref (7.2.0) from the genome and annotation of the 10x Genomics mm10-2020-A reference, with the Gq-DREADD and Gs-DREADD transgene sequences added as separate contigs, each annotated as a single gene. Ambient RNA was removed from each library’s raw count matrix with CellBender remove-back-ground ((Fleming et al., 2023); version 0.2.2; --total-droplets-included 50,000; otherwise default settings, including 150 epochs), run on NVIDIA A40 GPUs (PyTorch 1.13.1, CUDA 12.9). Spatial positions of nuclei estimated with the Slide-tags pipeline were joined to the gene-expression data by cell barcode.

#### 9.1.2. Quality control

Quality control used Scanpy (version 1.11.1; (Wolf et al., 2018)) and anndata (0.10.8; (Virshup et al., 2024)). Per sample, nuclei with fewer than 500 CellBender-corrected UMIs were excluded. Following the single-cell best practice recommendations (Heumos et al., 2023), nuclei in which mitochondrial genes contributed more than 8% of counts were removed. This affected only 7 nuclei (1 mouse, 6 human). Nuclei whose log10-transformed total counts or log10-transformed number of detected genes lay more than 3 median absolute deviations (MADs) from the sample median were removed (Supplementary Data S1).

Doublets were identified per sample with DoubletDetection (version 4.2; (Gayoso and Shor, 2020)), which performs Leiden clustering and selection of 3,000 highly variable genes internally. A boosting classifier was run for 40 iterations with a p-value threshold of 10−7 and a voter threshold of 0.5, and predicted doublets were removed. Doublet calls were identical in repeated runs on the same processor type (AMD Genoa or Turin) and thread count (8). Of 92,977 mouse and 97,245 human nuclei passing CellBender, the threshold filters removed 12,654 and 12,080 and the doublet classifier 11,016 and 15,016, leaving 69,307 mouse and 70,149 human nuclei for integration.

#### 9.1.3. Dimensionality reduction, clustering and annotation

For each species, nuclei from all samples were combined into a single AnnData object. For each object, 3,000 highly variable genes were selected with scanpy.pp.highly_variable_genes (flavor seurat_v3, on raw counts, sample as batch key, span 0.5) from a candidate pool that excluded mitochondrial and ribosomal genes and genes detected in fewer than 10 nuclei; all genes were retained in the object itself. An scVI variational auto-encoder ((Lopez et al., 2018); scvi-tools 1.4.2, (Gayoso et al., 2022); PyTorch 2.11.0) was trained with two encoder/-decoder layers, 30 latent dimensions and a negative-binomial likelihood on the raw counts of these genes with sample as batch key, for the default number of epochs with early stopping and seed = 0. A k-nearest-neighbor graph (k = 100) was built on the 30-dimensional latent space. Leiden clustering ((Traag et al., 2019) scanpy.tl.leiden with the igraph implementation (Csardi and Nepusz, 2006), random seed 0) was run at resolutions from 0.1 to 3.0.

Clusters were curated iteratively, and in each round, every cluster at the resolution under inspection was assessed on four lines of evidence: expression of a curated panel of canonical marker genes of mouse and human endometrial cell types, examined with scanpy.pl.dotplot on counts normalized to 10,000 per nucleus and log-transformed; the cluster’s top 20 genes from one-versus-rest Wilcoxon rank-sum tests against all remaining clusters (scanpy.tl.rank_genes_groups); its median quality-control metrics (detected genes, total counts, mito-chondrial and ribosomal fractions, and DoubletDetection score); and the contribution of each sample to the cluster. A cluster was removed when this evidence indicated low RNA content, residual ambient or doublet signal, or expression of the marker panels of more than one lineage. Each removal was followed by re-selection of highly variable genes, retraining of the scVI model and re-clustering of the remaining nuclei because the latent space is fit to the nuclei it contains, and the cycle was repeated until no cluster met the removal criteria. Curation was performed at Leiden resolution 3.0 in the whole-tissue objects, deliberately over-clustering relative to the expected number of cell classes so that populations co-expressing markers of more than one class resolved as discrete clusters. In mouse, 6 of 39 clusters were removed in the first round and none in the second; in human, no cluster met the criteria in the whole-tissue object.

Nuclei were then assigned to major cell classes from the marker panels (39 clusters for 6 classes in mouse; 36 clusters for 5 classes in human), and each class was split into its own object and re-integrated with the same scVI procedure, with highly variable genes re-selected on the subset and a 30-nearest-neighbor graph in place of the 100-nearest-neighbor graph used for the whole-ti objects. Three clusters in each species expressed the markers of two classes and were carried into both sub-objects, where the finer embedding resolved them (mouse: epithelial and vasculature, fibroblast and vasculature, immune and perimetrial; human: fibroblast and immune in two clusters, fibroblast and smooth muscle in one). A cluster that on review belonged to a class other than the one it had been assigned was transferred to that object rather than discarded; one vasculature cluster of the human fibroblast object was moved in this way. Curation within each sub-object proceeded as above for one to three rounds at a resolution chosen per object; the rounds, resolutions and clusters removed are listed in Table S6.

Cell types and subtypes were annotated by manual assignment of the surviving clusters from the same marker panels and ranked marker genes, at a resolution chosen per object and lower than the resolution used for curation (0.1 to 2.0 in mouse, 0.1 to 0.6 in human; Table S6). Objects that comprised a single cell type, vascular, smooth muscle and perimetrial cells in mouse, smooth muscle cells in human, were labeled for subtypes. The annotated sub-objects were then merged and re-integrated with scVI as above to give one whole-tissue atlas per species. In human, 48 nuclei that had received both a smooth muscle and an endometrial fibroblast label in their respective sub-objects were removed.

**Table S6:** Data curation cycles.

| Species | Object | Round | Leiden resolution | Clusters removed | Annotation resolution |
|---|---|---|---|---|---|
| mouse | all cells | 1 | 3.0 | 3, 4, 5, 11, 23, 32 | – |
|  | all cells | 2 | 3.0 | none; split by class | – |
|  | epithelial | 1 | 3.0 | 3, 6, 38 | – |
|  | epithelial | 2 | 3.0 | 5 | 0.1 |
|  | fibroblast | 1 | 1.5 | 16 | – |
|  | fibroblast | 2 | 1.5 | 19 | 1.5 |
|  | vasculature | 1 | 1.0 | 0, 2, 4, 11 | – |
|  | vasculature | 2 | 1.0 | 0 | all |
|  | immune | 1 | 3.0 | 0, 4, 19, 30 | – |
|  | immune | 2 | 2.0 | 1, 9, 15 | 2.0 |
|  | smooth muscle | 1 | 0.7 | 4 | all |
|  | perimetrium | 1 | 0.4 | 0, 3 | all |
| human | all cells | 1 | 3.0 | none; split by class | – |
|  | epithelial | 1 | 0.4 | 1 | 0.4 |
|  | fibroblast | 1 | 3.0 | 0, 7, 19*, 20, 35 | – |
|  | fibroblast | 2 | 0.6 | – | 0.6 |
|  | immune | 1 | 3.0 | 0, 1, 3, 4, 9, 10 | – |
|  | immune | 2 | 3.0 | 0, 4, 12 | – |
|  | immune | 3 | 0.4 | – | 0.4 |
|  | vasculature | 1 | 3.0 | 1, 2, 4, 5, 13, 17, 18, 19 | – |
|  | vasculature | 2 | 3.0 | 14 | 0.1 |
|  | smooth muscle | 1 | 3.0 | 3, 4, 9, 11, 13, 19, 20, 21, 23, 24, 26 | – |
|  | smooth muscle | 2 | – | – | all |

\* Transferred to the vascular object rather than discarded.

### 9.2. Cell type proportions and data visualization

#### 9.2.1. Cell type and subtype proportions

Proportions were computed from all annotated nuclei, including those without a spatial position. Within each sample, the nuclei of a compartment were counted by cell type or subtype and expressed as a percentage of that compartment, and the per-sample percentages were then averaged within each timepoint (mouse) or cycle phase (human), so that samples contribute equally regardless of how many nuclei they yielded. Major cell type proportions were computed over all annotated nuclei of a sample; immune subtype proportions over its immune nuclei alone, and endometrial fibroblast and decidual subtype proportions over its fibroblast and decidual nuclei alone, so that each panel sums to 100% within its compartment. Nuclei of a cell type not plotted in a given panel were excluded from both the numerator and the denominator of that panel.

#### 9.2.2. Spatial display

Nuclei are plotted at the coordinates assigned by Slide-tags. For display only, each section was rotated about the center of its bounding box and cropped to a window chosen per section, and human sections were additionally mirrored; the rotations and windows are fixed per section and identical across all figures showing that section. Where a quantity is shown, the color range is clipped to the 1st and 99th percentiles of the nuclei of that section carrying a value, shared across the panels of a figure unless stated otherwise, and nuclei are drawn in order of increasing value so that the highest values are visible.

### 9.3. Gene module scoring

#### 9.3.1. Decidual zone module scoring

Gene sets for the uterine microenvironments defined by (Li et al., 2022) at pregnancy day 7.5 were taken verbatim from their Table 1, which lists the top 20 marker genes per microenvironment, ranked by fold change and identified with Seurat FindAllMarkers on Harmony-integrated Visium profiles of four D7.5 uterine cross sections. We scored the primary, transitional and secondary decidual zones (PDZ, TDZ, SDZ), the vascular sinus zone (VSZ), the mesometrial decidua (MD) and undifferentiated stroma (udStr). Each set was scored per nucleus with scanpy.tl.score_genes (Satija et al., 2015), which subtracts from the mean expression of the set the mean expression of a control set of expression-matched genes (ctrl_size 50, random seed 0), on counts normalized to 10,000 per nucleus and log-transformed. Scoring was restricted to the fibroblast and decidual nuclei of the 3.5 dpi section, which is day 7.5 of pseudopregnancy and therefore matched in days to the pregnancy dataset. Genes detected in fewer than 1% of the nuclei of every group were dropped from their set, and a set was scored only where at least four genes remained; all six sets passed. Genes listed under more than one microenvironment were kept in each, because the published sets overlap by design and removing the shared genes would strip the decidual markers common to adjacent zones. The five endometrial fibroblast subtypes were pooled as eFib and compared with dec1, dec2 and dec3; apoptotic decidual nuclei (decAp) were not assigned to a group and carry no score. For the heat-map, scores were averaged within each group and z-scored across the four groups within each module, so that the color shows which group scores highest for a module rather than the absolute score, and are displayed from −1.8 to 1.8. In spatial maps each nucleus is colored by its score, with the range clipped to the 1st and 99th percentiles of the scored nuclei in view and shared across the six modules; the highest-scoring nuclei are drawn last, and stromal nuclei without a score are drawn underneath in dark gray.

#### 9.3.2. Stro-7 module scoring

To position our human fibro-blast and decidual subtypes relative to the stromal subtypes of the (Ulrich et al., 2024) atlas, each nucleus was scored for the 50 markers with the largest average log2 fold change of each stromal subtype (Dataset S6, sheet 6C) with scanpy.tl.score_genes as above (ctrl_size 50, random seed 0, on counts normalized to 10,000 per nucleus and log-transformed). Scores were averaged within each of the eight fibroblast and decidual subtypes (myoFib, eFib-a to eFib-c, dec-a to dec-d), pooling both donors, and z-scored across the eight subtypes within each module, displayed from −1.8 to 1.8. The C7-Fibro-blast module (Stro-7) was mapped onto the human sections, and the values transferred to X-Mens nuclei through the SAMap graph (see Cross-species alignment and transfer of categorical and continuous values) were mapped onto the mouse sections, with the color range clipped to the 1st and 99th percentiles of scored nuclei and shared across sections.

### 9.4. Pseudotime analysis

#### 9.4.1 Pseudotime

We ordered endometrial fibroblasts and decidual cells by diffusion pseudotime (DPT; (Hagh-verdi et al., 2016)) computed on a time-informed transition matrix, using CellRank 2 (v2.0.7; (Weiler and Theis, 2026)). Apoptotic nuclei were excluded from the pseudotime computation and carry no pseudotime value (1,175 decAp nuclei in mouse; 1,869 dec-c and dec-d nuclei in human). An scVI latent space was used for the remaining nuclei, fit on 3,000 highly variable genes (seurat_v3 on raw counts, after removing mitochondrial and ribosomal genes), with two layers, 30 latent dimensions, negative-binomial likelihood, sample as batch key, seed 0. Timepoints were encoded as days (mouse: uninduced 0, 1.5 dpi early 1, 1.5 dpi late 1.5, 2.5 dpi 2, 3.5 dpi 3, 4.5 dpi menstrual 4; human: secretory 0, menstrual 1). Transitions between consecutive timepoints were estimated by unbalanced entropic optimal transport with moscot (v0.5.0; (Klein et al., 2025); TemporalProblem), using squared Euclidean cost scaled by its mean in the scVI latent space. Entropic regularization was parameterized as ε = 10^−3^, and marginals were set as τa = τb = 0.95. Both marginals were relaxed because the sampled populations are not closed: each timepoint is a separate tissue sample, stromal cells proliferate and die during decidualization, and apoptotic decidual nuclei were excluded. Prior growth rates came from moscot’s proliferation and apoptosis gene sets. All couplings converged. The couplings were combined in a CellRank RealTimeKernel with the kernel settings of the CellRank protocol ((Weiler and Theis, 2026), Procedure 4: self_transitions = “all”, conn_weight = 0.2, threshold = “auto”), with within-time-point connectivities from 15-nearest-neighbor graphs built on principal components of each timepoint’s nuclei. Following the real-time-informed pseudotime (Weiler et al., 2024), we symmetrized the transition matrix ((T + T^T^)/2, rows renormalized) and computed a 15-component diffusion map (Coifman et al., 2005). In mouse, the root was chosen as the outer fibroblast (eFib1) nucleus with the lowest DC1 value, which was also the nucleus with the lowest DC1 value overall. In human, the DC4-DC1 embedding revealed two distinct endpoints at low DC1 that represented plausible roots, one within the myoFib cluster and one within eFib-a. We set the root within myoFib, corresponding to the outermost fibroblasts, as this positioning is analogous to the mouse root. The choice of endpoint changes the ordering of fibroblasts but not of decidual cells. DPT was computed with scanpy.tl.dpt on all 15 components and rank-normalized (average rank / N) for all downstream analyses (Figure 4D and Figure S21).

#### 9.4.2. Pseudotime-associated genes and cross-species conservation

Genes with expression associated with pseudotime were identified with a mutual information (MI)-based approach adapted from previous studies of the menstrual and mouse estrous cycles (Wang et al., 2020b; Winkler et al., 2024). All genes detected in at least 1% of pseudotime-carrying nuclei were tested (14,016 mouse and 15,354 human genes). For each such gene, MI between log-normalized expression and pseudotime was estimated on all nuclei with the k-nearest-neighbor estimator implemented in sklearn.feature_selection.mutual_info_regression with default settings (n_neighbors = 3) and a fixed random state (scikit-learn 1.8.0; (Krask et al., 2004; Pedregosa et al., 2011)). Significance was assessed by permuting pseudotime 1,000 times. Genes were processed in parallel chunks (200 in mouse, 1,500 in human). All genes in a chunk shared the same 1,000 permutations. The one-sided empirical p-value was (number of permutations with MI ≥ observed + 1)/(1,000 + 1). P-values were adjusted with the Benjamini-Hoc - berg procedure ((Benjamini and Hochberg, 1995); stats-models 0.14.6), and genes with q < 0.05 were considered significant (14,016 mouse and 15,187 human genes). Significance served as a floor excluding estimator noise rather than a strict selective criterion. Top pseudotime-associated genes were defined as those whose MI exceeded the 90th percentile among significant genes in each species: 1,402 mouse genes and 1,519 human genes (Figure S31).

Strict one-to-one human-mouse orthologs were obtained from the Ensembl Compara 116 human-mouse homologies of protein-coding genes ((Dyer et al., 2025); homology type ortholog_one2one). Pairs were matched to the genes of each object by Ensembl gene ID. For IDs absent from an object, the gene symbol was used when it matched a single gene, leaving a total of 16,246 pairs.

Of note, decidual prolactin, a classical marker of decidualization in both species, has no one-to-one mouse–human ortholog and is therefore absent from the list. Humans have a single prolactin gene (*PRL*), which is expressed in the decidua in addition to its regulated pituitary expression; mice have 23 prolactin-like genes, of which *Prl8a2* is the functional equivalent in terms of decidua-specific expression.

The background for comparing the top pseudotime-associated genes between species comprised the 9,800 one-to-one pairs in which both genes were detected in at least 1% of fibroblasts and decidual cells and significant for MI (q < 0.05) in their species, and the top gene sets were restricted to this background (1,153 mouse genes, K; 1,231 human genes, n). Their overlap (k = 379 pairs) was tested with a hypergeometric test (scipy.stats.hypergeom in SciPy, (Virtanen et al., 2020); N = 9,800, computed on the log scale with hypergeom.logsf): the expected overlap was K × n / N = 144.8 genes (12% of the human top set) and the observed overlap 31% (2.62-fold enrichment; p = 4.84 × 10−85). To check that the result did not depend on the 90th-percentile threshold, the test was repeated at every percentile cutoff from the 40th to the 99th, recording -log10 p, fold enrichment, and the 2.5th-97.5th percentile range of the hypergeometric null (Figure S30; Supplementary Data S2).

#### 9.4.3. Pseudotime expression heatmaps

To compare gene expression changes during induced decidualization to pregnancy in (Yang et al., 2023), mouse nuclei subsetted from induced timepoints were re-ranked by their pseudotime values. Smoothed expression profiles of the 28 panel genes (Figure 4H) were computed with Monocle 2 (v2.34.0; (Qiu et al., 2017; Trapnell et al., 2014)), using the calculation of its plot_pseudotime_heatmap function: each gene’s size-factor-normalized UMI counts were fitted with a negative-binomial generalized linear model (VGAM; (Yee, 2015)) on a natural cubic spline of pseudo-time with 3 degrees of freedom. Size factors and the dispersion-mean trend that supplies each gene’s dispersion were estimated with Monocle’s default settings on all mouse nuclei with a pseudotime. The fitted curves, evaluated at 100 evenly spaced pseudotime points, were log10(x + 1)-transformed, z-scored per gene and clipped to [-3, 3]. Rows follow the gene order of (Yang et al., 2023).

For the heatmaps of all top pseudotime-associated genes, the genes of each species were its top genes within the background of the overlap test above. Smoothed profiles were computed with the Monocle 2 procedure above on all nuclei with a pseudotime. Rows were ordered by pseudotime centroid, the mean position along pseudotime weighted by the positive part of the z-scored profile, separately in each species (Figure S31B).

#### 9.4.4 Notum expression along pseudotime

*Notum* expression in each mouse fibroblast and decidual nucleus with a pseudotime value was computed from log-normalized UMI counts (Figure S23B). Nuclei were plotted against rank-normalized pseudotime and colored by group: all fibroblast subtypes together, then dec1, dec2, and dec3 separately. The trend line is the mean expression within 25 equal-width pseudotime bins. The 95% bootstrap confidence intervals of bin means over 500 resamples are drawn, but are narrower than the line in most bins.

### 9.5. Cross-species alignment and transfer of categorical and continuous values (SAMap)

SAMap was used to integrate mouse and human Slide-tags datasets. SAMap first builds a cross-species gene graph from all homologous gene pairs rather than from a list of orthologs: links between sequence-similar genes, including paralogs and one-to-many relationships, are initialized from sequence similarity and reweighted by expression correlation as the alignment is refined (Tarashansky et al., 2021). Human and mouse proteomes (Ensembl release 113; GRCh38, GRCm39) were searched reciprocally with phmmer (HMMER 3.4; (Eddy, 2011)) at E ≤ 10^−5^. Because the proteomes include all annotated isoforms, hits were collapsed to genes, keeping the highest-scoring isoform pair for each pair of genes. Proteins lacking a gene symbol were excluded. SAMap matches genes between the count matrices and the homology tables by name, and some genes are named differently in the two because they come from different annotation versions. Gene names in the homology tables were then matched to those of the count matrices by Ensembl gene ID, since the two come from different annotation releases. When building its homology graph, SAMap retained only pairs with E ≤ 10^−6^ (its default threshold).

Fibroblast and decidual nuclei from all timepoints were supplied to SAMap (sc-samap 3.0.1) as raw integer counts indexed by the count matrices’ gene names. Per-species SAM manifolds were built with SAMap’s default settings and stitched over the homology tables for three iterations, keeping up to 20 mutual cross-species neighbors per nucleus, with edge weights rescaled by cross-species expression correlation. The fit was made reproducible by fixing numpy’s random seed (0) and restricting nearest-neighbor index construction (hnswlib 0.8.0) to a single thread.

Values were transferred between species with a single kernel-weighted average, using the SAMap cross-species edge weights of each receiving nucleus as the kernel (a Nadaraya-Watson-type linear smoother; (Hastie et al., 2009; Nadaraya, 1964)). For a receiving nucleus i in one species, with w_ij_ the edge weight to nucleus j of the other species, the normalized weights are 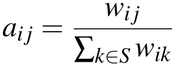 where S is the set of cross-species neighbors of i that carry the quantity being transferred. A continuous value v is transferred as its weighted mean, 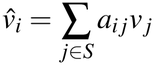

A categorical value y is transferred as the weighted share of each category c, 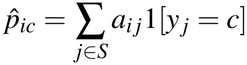

The shares of a nucleus sum to 1, and where a single label is needed it is the category with the largest share, 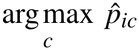

The weights are non-negative and derived from expression and sequence homology alone, independent of every transferred quantity. A nucleus with no cross-species neighbor carrying the quantity received no value.

From mouse to human, we transferred cell identity (categorical: decidual cell or endometrial fibroblast) and pseudotime (continuous: mouse pseudotime). Every mouse nucleus carries a label, whereas the apoptotic decidual nuclei carry no pseudotime. Nine human nuclei had no mouse neighbor and received neither value, and a further 384 had only apoptotic mouse neighbors and received no pseudotime. Transferred labels were compared with the human annotation, and transferred pseudotime with human pseudotime by Spearman correlation over all nuclei and within decidual cells and endometrial fibroblasts (Figure 6E and Figure S32). To summarize how the two cell types correspond in the SAMap graph, the cross-species edge weights of all nuclei of a cell type (endometrial fibroblast or decidual) were summed, and the share going to each compartment of the other species was computed, in both directions (Figure S32).

To test whether the transferred pseudotime gradient could arise from the SAMap graph or the averaging alone, mouse pseudotime values were permuted 1,000 times among mouse nuclei before transfer, with the graph and edge weights held fixed, either across all mouse nuclei with a pseudotime value or only among nuclei of the same mouse cell type (endometrial fibroblast or decidual). The latter preserves the difference between compartments but removes the order within them. For each permutation, we computed the Spearman correlation between transferred and human pseudotime over all nuclei and within each compartment, and the spatial autocorrelation of transferred values over all nuclei and within each compartment. Spatial autocorrelation was measured as Moran’s I with row-standardized weights on the 10 nearest spatial neighbors, computed separately for each of the two tissue sections and averaged. Within a compartment, neighbors were drawn from that compartment only (Figure S32).

### 9.6. Differential gene expression

#### 9.6.1. Identification of differentially expressed genes in X-Mens uteri

Differentially expressed genes between fibroblasts and decidual cells in X-Mens uteri were identified for comparison with a published dataset comparing oil-stimulated and unstimulated uterine horns on day 8.5 of pseudopregnancy (Wang et al., 2020a). These analyses were restricted to the 3.5 dpi and 4.5 dpi X-Mens samples as the most closely matched timepoints. Apoptotic decidual nuclei (decAp) were excluded, leaving 15,749 nuclei. Raw counts were summed within each animal and cell type, pooling the 10X lanes of an animal into a single profile so that lanes were never treated as replicates. Profiles of fewer than 20 nuclei were discarded and only animals contributing both cell types were retained, giving four profiles from two animals. Genes with at least 10 counts in at least two profiles were tested (17,195 genes). Differential expression was computed on pseudobulk profiles with PyDESeq2 ((version 0.5.4; (Muzellec et al., 2023)), a Python implementation of DESeq2 (Love et al., 2014). The model was fit with the design ∼animal + condition and Cook’s distance refitting, and decidual cells were contrasted against endometrial fibroblasts, so that a positive log2 fold change indicates higher expression in decidual cells. Genes were called differentially expressed at a Benjamini-Hochberg adjusted (Benjamini and Hochberg, 1995) p-value of 0.01 or below, an absolute fold change of 2 or above, and a mean above 3 counts per million in at least one of the two cell types, the fold-change and false discovery rate thresholds of (Wang et al., 2020a); 1,438 genes were higher in decidual cells and 1,499 in endometrial fibroblasts. With one animal per timepoint the comparison rests on two biological replicates, leaving fewer than three residual degrees of freedom, so dispersions are estimated imprecisely and the adjusted p-values are better read as a ranking than as calibrated error rates.

#### 9.6.2. Comparison with oil-induced artificial decidualization

We used the differentially expressed genes reported by (Wang et al., 2020a) for oil-stimulated against unstimulated uterine horns on day 8.5 of pseudopregnancy (their Supplementary Table S3; 4,294 genes, 1,127 increased and 3,167 decreased), measured by bulk RNA sequencing of whole horns. Each was matched to an X-Mens gene by Ensembl identifier and, for identifiers absent from our annotation, by current gene symbol (3,752 matched by identifier, 3 by symbol, 539 not present in the X-Mens data); where several entries mapped to one X-Mens gene, one was retained. Genes called differentially expressed in both datasets were classified as shared up where both increased, shared down where both decreased, and divergent where the directions differed. Of the 1,350 genes called in both, 555 (41%) were shared up, 683 (51%) shared down and 112 (8%) divergent. The published table lists only genes that reached significance, so a gene absent from it cannot be distinguished from one that was tested and not called; the comparison is therefore confined to genes called in both datasets, and these proportions describe that set rather than either dataset as a whole. The two contrasts are also not equivalent in kind: the oil-induction contrast is between whole horns and reflects both expression changes within cells and shifts in cell composition, whereas the X-Mens contrast is between annotated cell populations. Gene-level results are given in Supplementary Data S3.

### 9.2. Code and data availability

All datasets generated for this study will be publicly available upon publication. Code is available at github.com/McKinleyLab/xmens under Git tag v2.0.

## Acknowledgments

We thank Len Zon, Iain Cheeseman, Ya-Chieh Hsu, Michalis Averof, Paola Arlotta, Ophir Klein, Ron Vale, Mathilde Paris, and Martin Hemberg for their feedback on the manuscript. We are grateful to the National Disease Research Interchange and donors and their families for donations of human uteri for this work and to Surya Nagaraja and Taylor Skokan for processing the tissues. We thank Jonathan Hecht for human endometrial staging, Rebecca Berdeaux and Bruce Morgan for the GsD mouse, Ron Chandler for the Ltf-iCre and Amhr2-Cre mice, and Ute Hochgeschwender for sharing the GqD transgene sequence. We thank Ângela Gonçalves, Sue Hammoud, Peng Du and Min Yang for sharing data. We thank Kristy Red-Horse, James Zwierzynski, and Philipp Weiler for helpful discussions. We thank Kathleen Pritchett-Corning, Eny Maldonado, and the Harvard Office of Animal Resources for animal care, Timothy Fenn and Lepka Gagoska for administrative support, the Harvard Faculty of Arts and Sciences Research Computing Core for computational resources, the HSCRB histology core, and the Harvard Center for Biological Imaging.

## Funding

Soozin White Research Fund at Harvard Medical School (ÇÇ)

National Science Foundation Graduate Research Fellowship Award Program (CJA)

Human Frontiers Science Program (FSC)

National Institutes of Health grant DP2HD111708 (KLM)

National Institutes of Health grant R00HD101021 (KLM)

New York Stem Cell Foundation (NYSCF)

Robertson Stem Cell Investigator (KLM)

Howard Hughes Medical Institute (KLM)

Charles H. Hood Foundation (KLM)

David and Lucile Packard Foundation (KLM)

Searle Scholars Program (KLM)

Damon Runyon Cancer Research Foundation (KLM)

Massachusetts Life Sciences Center (KLM)

Smith Family Foundation (KLM)

## Author contributions

Conceptualization: ÇÇ, KLM

Methodology: ÇÇ, NJH, AJCR, MT

Investigation: ÇÇ, NJH, AMK, AJCR, AEG, CJA, PLM, EMW, KCL, AYBV, SGB, DHK, DAH, LEB, AB, JLRG, DJDB, MT, JP, FSC, DKS, KLM

Visualization: ÇÇ, NJH, CJA, PLM

Funding acquisition: ÇÇ, KEO, RAF, FC, SRE, KLM

Project administration: ÇÇ, KLM

Supervision: ÇÇ, CJA, RAF, SRE, FC, KLM

Writing – original draft: ÇÇ

Writing – review & editing: ÇÇ, NJH, AJCR, AEG, CJA, PLM, EMW, SGB, KCL, AYBV, DHK, DAH, LEB, AB, DJDB, MT, JP, FSC, DKS, KEO, RAF, SRE, KLM

Software: ÇÇ, NJH, LEB

Formal analysis: ÇÇ, NJH, LEB

Data curation: ÇÇ, NJH, AJCR, LEB

## Competing interests

The authors declare no competing interests.

## Data and materials availability

Requests for materials should be directed to the corresponding author. Code is available at github.com/McKinleyLab/xmens.

## Notes

### Competing Interest Statement

The authors have declared no competing interest.

### Summary of Updates

Full manuscript and supplemental materials revised.

## References

Ainslie, R. J., Kirkwood, P. M., Li, H., Gerasimou, S., Saunders, P. T. K., Simitsidellis, I. and Gibson, D. A. (2026). Multimodal profiling of repair-associated immune cells in a mouse model of menstruation. J. Reprod. Immunol. 178, 104981.

Akhmedov, D., Mendoza-Rodriguez, M. G., Rajendran, K., Rossi, M., Wess, J. and Berdeaux, R. (2017). Gs-DREADD Knock-In Mice for Tissue-Specific, Temporal Stimulation of Cyclic AMP Signaling. Mol. Cell. Biol. 37, e00584–16.

Ang, C. J., Gable, J. L. R., Lyons, K. C., Whelan, E. M., Çevrim, Ç., Skokan, T. D., Bennetts, S. G., Manetta, B. D., Kaage, A. M., Mopure, D., et al. (2026). Luminal epithelium remodeling underlies endometrial regeneration during menstruation and pregnancy. bioRxivorg 2026.03.08.710375.

Armbruster, B. N., Li, X., Pausch, M. H., Herlitze, S. and Roth, B. L. (2007). Evolving the lock to fit the key to create a family of G protein-coupled receptors potently activated by an inert ligand. Proc. Natl. Acad. Sci. U. S. A. 104, 5163–5168.

Armstrong, G. M., Maybin, J. A., Murray, A. A., Nicol, M., Walker, C., Saunders, P. T. K., Rossi, A. G. and Critchley, H. O. D. (2017). Endometrial apoptosis and neutrophil infiltration during menstruation exhibits spatial and temporal dynamics that are recapitulated in a mouse model. Sci. Rep. 7, 17416.

Atkinson, W. B. and Hooker, C. W. (1945). The day to day level of estrogen and progestin throughout pregnancy and pseudopregnancy in the mouse. Anat. Rec. 93, 75–95.

Bellofiore, N., Rana, S., Dickinson, H., Temple-Smith, P. and Evans, J. (2018). Characterization of human-like menstruation in the spiny mouse: comparative studies with the human and induced mouse model. Hum. Reprod. 33, 1715–1726.

Belsey, E. M. and Pinol, A. P. (1997). Menstrual bleeding patterns in untreated women. Task Force on Long-Acting Systemic Agents for Fertility Regulation. Contraception 55, 57–65.

Benjamini, Y. and Hochberg, Y. (1995). Controlling the false discovery rate: A practical and powerful approach to multiple testing. J. R. Stat. Soc. Series B Stat. Methodol. 57, 289–300.

Bezanson, J., Edelman, A., Karpinski, S. and Shah, V. B. (2017). Julia: A fresh approach to numerical computing. SIAM Rev. Soc. Ind. Appl. Math. 59, 65–98.

Brar, A. K., Frank, G. R., Kessler, C. A., Cedars, M. I. and Handwerger, S. (1997). Progesterone-dependent decidualization of the human endometrium is mediated by cAMP. Endocrine 6, 301–307.

Brasted, M., White, C. A., Kennedy, T. G. and Salamonsen, L. A. (2003). Mimicking the events of menstruation in the murine uterus. Biol. Reprod. 69, 1273–1280.

Brighton, P. J., Maruyama, Y., Fishwick, K., Vrljicak, P., Tewary, S., Fujihara, R., Muter, J., Lucas, E. S., Yamada, T., Woods, L., et al. (2017). Clearance of senescent decidual cells by uterine natural killer cells in cycling human endometrium. Elife 6, e31274.

Brommage, R., Liu, J., Vogel, P., Mseeh, F., Thompson, A. Y., Potter, D. G., Shadoan, M. K., Hansen, G. M., Jeter-Jones, S., Cui, J., et al. (2019). NOTUM inhibition increases endocortical bone formation and bone strength. Bone Res. 7, 2.

Buxton, L. E. and Murdoch, R. N. (1982). Lectins, calcium ionophore A23187 and peanut oil as deciduogenic agents in the uterus of pseudopregnant mice: Effects of tranylcypromine, indomethacin, iproniazid and propanolol. Aust. J. Biol. Sci. 35, 63.

Catalini, L. and Fedder, J. (2020). Characteristics of the endometrium in menstruating species: lessons learned from the animal kingdom. Biol. Reprod. 102, 1160–1169.

Coifman, R. R., Lafon, S., Lee, A. B., Maggioni, M., Nadler, B., Warner, F. and Zucker, S. W. (2005). Geometric diffusions as a tool for harmonic analysis and structure definition of data: diffusion maps. Proc. Natl. Acad. Sci. U. S. A. 102, 7426–7431.

Cousins, F. L., Filby, C. E. and Gargett, C. E. (2021). Endometrial stem/progenitor cells-their role in endometrial repair and regeneration. Front. Reprod. Health 3, 811537.

Critchley, H. O. D., Maybin, J. A., Armstrong, G. M. and Williams, A. R. W. (2020). Physiology of the endometrium and regulation of menstruation. Physiol. Rev. 100, 1149–1179.

Csardi, G. and Nepusz, T. (2006). The igraph software package for complex network research. InterJournal. Complex Systems 1695, 1–9.

Cunha, G. R., Sinclair, A., Ricke, W. A., Robboy, S. J., Cao, M. and Baskin, L. S. (2019). Reproductive tract biology: Of mice and men. Differentiation 110, 49–63.

Daikoku, T., Ogawa, Y., Terakawa, J., Ogawa, A., DeFalco, T. and Dey, S. K. (2014). Lactoferrin-iCre: a new mouse line to study uterine epithelial gene function. Endocrinology 155, 2718–2724.

Dasharathy, S. S., Mumford, S. L., Pollack, A. Z., Perkins, N. J., Mattison, D. R., Wactawski-Wende, J. and Schisterman, E. F. (2012). Menstrual bleeding patterns among regularly menstruating women. Am. J. Epidemiol. 175, 536–545.

Dyer, S. C., Austine-Orimoloye, O., Azov, A. G., Barba, M., Barnes, I., Barrera-Enriquez, V. P., Becker, A., Bennett, R., Beracochea, M., Berry, A., et al. (2025). Ensembl 2025. Nucleic Acids Res. 53, D948–D957.

Eddy, S. R. (2011). Accelerated profile HMM searches. PLoS Comput. Biol. 7, e1002195.

Emera, D., Romero, R. and Wagner, G. (2012). The evolution of menstruation: a new model for genetic assimilation: explaining molecular origins of maternal responses to fetal invasiveness. Bioessays 34, 26–35.

Finn, C. A. and Pope, M. (1984). Vascular and cellular changes in the decidualized endometrium of the ovariectomized mouse following cessation of hormone treatment: a possible model for menstruation. J. Endocrinol. 100, 295–300.

Fitzgerald, H. C., Mortlock, S., Filby, C. E., Cousins, F. L., Marquez-Garcia, E., Wyatt, K. A., McKinnon, B., Rombauts, L., Tsaltas, J., Montgomery, G. W., et al. (2026). Molecular signature of human endometrial stem/progenitor cells at the single-cell level. Stem Cell Reports 21, 102952.

Fleming, S. J., Chaffin, M. D., Arduini, A., Akkad, A.-D., Banks, E., Marioni, J. C., Philippakis, A. A., Ellinor, P. T. and Babadi, M. (2023). Unsupervised removal of systematic background noise from drop-let-based single-cell experiments using CellBender. Nat. Methods 20, 1323–1335.

Franco, H. L., Dai, D., Lee, K. Y., Rubel, C. A., Roop, D., Boerboom, D., Jeong, J.-W., Lydon, J. P., Bagchi, I. C., Bagchi, M. K., et al. (2011). WNT4 is a key regulator of normal postnatal uterine development and progesterone signaling during embryo implantation and decidualization in the mouse. FASEB J. 25, 1176–1187.

Gaide Chevronnay, H. P., Galant, C., Lemoine, P., Courtoy, P. J., Marbaix, E. and Henriet, P. (2009). Spatiotemporal coupling of focal extracellular matrix degradation and reconstruction in the menstrual human endometrium. Endocrinology 150, 5094–5105.

Garry, R., Hart, R., Karthigasu, K. A. and Burke, C. (2009). A re-appraisal of the morphological changes within the endometrium during menstruation: a hysteroscopic, histological and scanning electron microscopic study. Hum. Reprod. 24, 1393–1401.

Gayoso, A. and Shor, J. (2020). JonathanShor/Doublet-Detection: doubletdetection v3.0. Zenodo.

Gayoso, A., Lopez, R., Xing, G., Boyeau, P., Valiollah Pour Amiri, V., Hong, J., Wu, K., Jayasuriya, M., Mehlman, E., Langevin, M., et al. (2022). A Python library for probabilistic analysis of single-cell omics data. Nat. Biotechnol. 40, 163–166.

Gellersen, B. and Brosens, J. (2003). Cyclic AMP and progesterone receptor cross-talk in human endometrium: a decidualizing affair. J. Endocrinol. 178, 357–372.

Gellersen, B. and Brosens, J. J. (2014). Cyclic decidualization of the human endometrium in reproductive health and failure. Endocr. Rev. 35, 851–905.

Greaves, E., Cousins, F. L., Murray, A., Esnal-Zufiaurre, A., Fassbender, A., Horne, A. W. and Saunders, P. T. K. (2014). A novel mouse model of endometriosis mimics human phenotype and reveals insights into the inflammatory contribution of shed endometrium. Am. J. Pathol. 184, 1930–1939.

Haghverdi, L., Büttner, M., Wolf, F. A., Buettner, F. and Theis, F. J. (2016). Diffusion pseudotime robustly reconstructs lineage branching. Nat. Methods 13, 845–848.

Hastie, T., Tibshirani, R. and Friedman, J. (2009). The elements of statistical learning: Data mining, inference, and prediction, second edition. 2nd ed. New York, NY: Springer.

Heumos, L., Schaar, A. C., Lance, C., Litinetskaya, A., Drost, F., Zappia, L., Lücken, M. D., Strobl, D. C., Henao, J., Curion, F., et al. (2023). Best practices for single-cell analysis across modalities. Nat. Rev. Genet. 24, 550–572.

Hirota, Y., Daikoku, T., Tranguch, S., Xie, H., Bradshaw, H. B. and Dey, S. K. (2010). Uterine-specific p53 deficiency confers premature uterine senescence and promotes preterm birth in mice. J. Clin. Invest. 120, 803–815.

Houser, B. L., Tilburgs, T., Hill, J., Nicotra, M. L. and Strominger, J. L. (2011). Two unique human decidual macrophage populations. J. Immunol. 186, 2633–2642.

Idelfonso-García, O. G., Pacheco-Rivera, R., Alarcón-Sánchez, B. R., Serrano-Luna, J., Baltiérrez-Hoyos, R., Vásquez-Garzón, V. R., Muriel, P., Villa-Treviño, S., Pérez-Carreón, J. I. and Arellanes-Robledo, J. (2024). Protocol to detect senescence-associated β-galactosidase and immunoperoxidase activity in fresh-frozen murine tissues. STAR Protoc. 5, 103009.

Jamin, S. P., Arango, N. A., Mishina, Y., Hanks, M. C. and Behringer, R. R. (2002). Requirement of Bmpr1a for Müllerian duct regression during male sexual development. Nat. Genet. 32, 408–410.

Kaitu’u-Lino, T. J., Morison, N. B. and Salamonsen, L. A. (2007a). Neutrophil depletion retards endometrial repair in a mouse model. Cell Tissue Res. 328, 197–206.

Kaitu’u-Lino, T. J., Morison, N. B. and Salamonsen, L. A. (2007b). Estrogen is not essential for full endometrial restoration after breakdown: lessons from a mouse model. Endocrinology 148, 5105–5111.

Kaitu’u-Lino, T. J., Ye, L. and Gargett, C. E. (2010). Reepithelialization of the uterine surface arises from endometrial glands: evidence from a functional mouse model of breakdown and repair. Endocrinology 151, 3386–3395.

Kakugawa, S., Langton, P. F., Zebisch, M., Howell, S., Chang, T.-H., Liu, Y., Feizi, T., Bineva, G., O’Reilly, N., Snijders, A. P., et al. (2015). Notum deacylates Wnt proteins to suppress signalling activity. Nature 519, 187–192.

Kawarabayashi, Y., Hai, L., Honda, A., Horiuchi, S., Tsujioka, H., Ichikawa, J. and Inoue, R. (2012). Critical role of TRPC1-mediated Ca2+ entry in decidualization of human endometrial stromal cells. Mol. Endocrinol. 26, 846–858.

Kim, M., Park, H. J., Seol, J. W., Jang, J. Y., Cho, Y.-S., Kim, K. R., Choi, Y., Lydon, J. P., Demayo, F. J., Shibuya, M., et al. (2013). VEGF-A regulated by progesterone governs uterine angiogenesis and vascular remodelling during pregnancy: Uterine angiogenesis and vascular remodelling. EMBO Mol. Med. 5, 1415–1430.

Kirkwood, P. M., Gibson, D. A., Shaw, I., Dobie, R., Kelepouri, O., Henderson, N. C. and Saunders, P. T. K. (2022). Single-cell RNA sequencing and lineage tracing confirm mesenchyme to epithelial transformation (MET) contributes to repair of the endometrium at menstruation. Elife 11:e77663.

Klein, D., Palla, G., Lange, M., Klein, M., Piran, Z., Gander, M., Meng-Papaxanthos, L., Sterr, M., Saber, L., Jing, C., et al. (2025). Mapping cells through time and space with moscot. Nature 638, 1065–1075.

Konno, R., Igarashi, T., Okamoto, S., Sato, S., Moriya, T., Sasano, H. and Yajima, A. (1999). Apoptosis of human endometrium mediated by perforin and granzyme B of NK cells and cytotoxic T lymphocytes. Tohoku J. Exp. Med. 187, 149–155.

Kraskov, A., Stögbauer, H. and Grassberger, P. (2004). Estimating mutual information. Phys. Rev. E Stat. Nonlin. Soft Matter Phys. 69, 066138.

Leroy, F., Vansande, J., Shetgen, G. and Brasseur, D. (1974). Cyclic AMP and the triggering of the decidual reaction. J. Reprod. Fertil. 39, 207–211.

Li, Q., Kannan, A., Wang, W., Demayo, F. J., Taylor, R. N., Bagchi, M. K. and Bagchi, I. C. (2007). Bone morphogenetic protein 2 functions via a conserved signaling pathway involving Wnt4 to regulate uterine decidualization in the mouse and the human. J. Biol. Chem. 282, 31725–31732.

Li, Q., Kannan, A., Das, A., Demayo, F. J., Hornsby, P. J., Young, S. L., Taylor, R. N., Bagchi, M. K. and Bagchi, I. C. (2013). WNT4 acts downstream of BMP2 and functions via β-catenin signaling pathway to regulate human endometrial stromal cell differentiation. Endocrinology 154, 446–457.

Li, R., Wang, T.-Y., Xu, X., Emery, O. M., Yi, M., Wu, S.-P. and DeMayo, F. J. (2022). Spatial transcriptomic profiles of mouse uterine microenvironments at pregnancy day 7.5. Biol. Reprod. 107, 529–545.

Liu, T., Shi, F., Ying, Y., Chen, Q., Tang, Z. and Lin, H. (2020). Mouse model of menstruation: An indispensable tool to investigate the mechanisms of menstruation and gynaecological diseases (Review). Mol. Med. Rep. 22, 4463–4474.

Loeb, L. (1908). The experimental production of the maternal part of the placenta in the rabbit. Exp. Biol. Med. (Maywood) 5, 102–104.

Lopez, R., Regier, J., Cole, M. B., Jordan, M. I. and Yosef, N. (2018). Deep generative modeling for single-cell transcriptomics. Nat. Methods 15, 1053–1058.

Lorenzi, V., Moullet, M., Icoresi-Mazzeo, C., Colligan, R., Danks, M. R., Cohen, C. E., Parraga, A., Kim, C. E., Sancho-Serra, C., Sakata, R., et al. (2026). Full-thickness spatial transcriptomics of the human uterus reveals basalis niche architecture and regeneration gradients during menstrual breakdown. bioRxiv 2026.06.08.730878.

Love, M. I., Huber, W. and Anders, S. (2014). Moderated estimation of fold change and dispersion for RNA-seq data with DESeq2. Genome Biol. 15, 550.

Macdonald, L. J., Sales, K. J., Grant, V., Brown, P., Jabbour, H. N. and Catalano, R. D. (2011). Prokineticin 1 induces Dickkopf 1 expression and regulates cell proliferation and decidualization in the human endometrium. Mol. Hum. Reprod. 17, 626–636.

Marečková, M., Garcia-Alonso, L., Moullet, M., Lorenzi, V., Petryszak, R., Sancho-Serra, C., Oszlanczi, A., Icoresi Mazzeo, C., Wong, F. C. K., Kelava, I., et al. (2024). An integrated single-cell reference atlas of the human endometrium. Nat. Genet. 56, 1925–1937.

Maybin, J. A., Murray, A. A., Saunders, P. T. K., Hirani, N., Carmeliet, P. and Critchley, H. O. D. (2018). Hypoxia and hypoxia inducible factor-1α are required for normal endometrial repair during menstruation. Nat. Commun. 9, 295.

Muzellec, B., Teleńczuk, M., Cabeli, V. and Andreux, M. (2023). PyDESeq2: a python package for bulk RNA-seq differential expression analysis. Bioinformatics 39, btad547.

Muzumdar, M. D., Tasic, B., Miyamichi, K., Li, L. and Luo, L. (2007). A global double-fluorescent Cre reporter mouse. Genesis 45, 593–605.

Nadaraya, E. A. (1964). On estimating regression. Theory Probab. Appl. 9, 141–142.

Nayak, N. R., Critchley, H. O., Slayden, O. D., Menrad, A., Chwalisz, K., Baird, D. T. and Brenner, R. M. (2000). Progesterone withdrawal up-regulates vascular endothelial growth factor receptor type 2 in the superficial zone stroma of the human and macaque endometrium: potential relevance to menstruation. J. Clin. Endocrinol. Metab. 85, 3442–3452.

Noyes, R. W., Hertig, A. T. and Rock, J. (1975). Dating the endometrial biopsy. Am. J. Obstet. Gynecol. 122, 262–263.

Orwig, K. E., Ishimura, R., Müller, H., Liu, B. and Soares, M. J. (1997). Identification and characterization of a mouse homolog for decidual/trophoblast prolactin-related protein. Endocrinology 138, 5511–5517.

Parkes, A. S. (1929). The functions of the corpus luteum. II.—The experimental production of placentomata in the mouse. Proc. R. Soc. Lond. B Biol. Sci. 104, 183–188.

Pedregosa, F., Varoquaux, G., Gramfort, A., Michel, V., Thirion, B., Grisel, O., Blondel, M., Louppe, G., Prettenhofer, P., Weiss, R., et al. (2011). Scikit-learn: Machine Learning in Python. J. Mach. Learn. Res. 12, 2825–2830.

Peterse, D., Clercq, K. D., Goossens, C., Binda, M. M., F O, D., Saunders, P., Vriens, J., Fassbender, A. and D’Hooghe, T. M. (2018). Optimization of endometrial decidualization in the menstruating mouse model for preclinical endometriosis research. Reprod. Sci. 25, 1577–1588.

Qiu, X., Mao, Q., Tang, Y., Wang, L., Chawla, R., Pliner, H. A. and Trapnell, C. (2017). Reversed graph embedding resolves complex single-cell trajectories. Nat. Methods 14, 979–982.

Ramathal, C. Y., Bagchi, I. C., Taylor, R. N. and Bagchi, M. K. (2010). Endometrial decidualization: of mice and men. Semin. Reprod. Med. 28, 17–26.

Rankin, J. C., Ledford, B. E. and Baggett, B. (1977). Early involvement of cyclic nucleotides in the artificially stimulated decidual cell reaction in the mouse uterus. Biol. Reprod. 17, 549–554.

Rawlings, T. M., Makwana, K., Taylor, D. M., Molè, M. A., Fishwick, K. J., Tryfonos, M., Odendaal, J., Hawkes, A., Zernicka-Goetz, M., Hartshorne, G. M., et al. (2021). Modelling the impact of decidual senescence on embryo implantation in human endometrial assembloids. Elife 10:e69603.

Rogers, L. M., Lash, G. E., Anderson, G. M. and Girling, J. E. (2025). Modelling menstruation in the common mouse: a narrative review. Reprod. Fertil. Dev. 37, RD25055.

Rudolph, M., Döcke, W.-D., Müller, A., Menning, A., Röse, L., Zollner, T. M. and Gashaw, I. (2012). Induction of overt menstruation in intact mice. PLoS One 7, e32922.

Russell, A. J. C., Weir, J. A., Nadaf, N. M., Shabet, M., Kumar, V., Kambhampati, S., Raichur, R., Marrero, G. J., Liu, S., Balderrama, K. S., et al. (2024). Slide-tags enables single-nucleus barcoding for multimodal spatial genomics. Nature 625, 101–109.

Rytkönen, K. T., Erkenbrack, E. M., Poutanen, M., Elo, L. L., Pavlic, M. and Wagner, G. P. (2019). Decidualization of human endometrial stromal fibroblasts is a multiphasic process involving distinct transcriptional programs. Reprod. Sci. 26, 323–336.

Sakoff, J. A. and Murdoch, R. N. (1996). The role of calcium in the artificially induced decidual cell reaction in pseudopregnant mice. Biochem. Mol. Med. 57, 81–90.

Salamonsen, L. A., Hutchison, J. C. and Gargett, C. E. (2021). Cyclical endometrial repair and regeneration. Development 148, 17.

Satija, R., Farrell, J. A., Gennert, D., Schier, A. F. and Regev, A. (2015). Spatial reconstruction of single-cell gene expression data. Nat. Biotechnol. 33, 495–502.

Schindelin, J., Arganda-Carreras, I., Frise, E., Kaynig, V., Longair, M., Pietzsch, T., Preibisch, S., Rueden, C., Saal-feld, S., Schmid, B., et al. (2012). Fiji: an open-source platform for biological-image analysis. Nat. Methods 9, 676–682.

Sekiba, D. (1923). Zur Morphologie und Histologie des Menstruationszyklus. Arch. Gynak. 121, 36–60.

Shaw, I. W., Kirkwood, P. M., Rebourcet, D., Cousins, F. L., Ainslie, R. J., Livingstone, D. E. W., Smith, L. B., Saunders, P. T. K. and Gibson, D. A. (2022). A role for steroid 5 alpha-reductase 1 in vascular remodeling during endometrial decidualization. Front. Endocrinol. (Lausanne) 13, 1027164.

Stringer, C., Wang, T., Michaelos, M. and Pachitariu, M. (2021). Cellpose: a generalist algorithm for cellular segmentation. Nat. Methods 18, 100–106.

Strunz, B., Bister, J., Jönsson, H., Filipovic, I., Crona-Guterstam, Y., Kvedaraite, E., Sleiers, N., Dumitrescu, B., Brännström, M., Lentini, A., et al. (2021). Continuous human uterine NK cell differentiation in response to endometrial regeneration and pregnancy. Sci. Immunol. 6, eabb7800.

Tang, B., Guller, S. and Gurpide, E. (1993). Cyclic adenosine 3’,5’-monophosphate induces prolactin expression in stromal cells isolated from human proliferative endometrium. Endocrinology 133, 2197–2203.

Tarashansky, A. J., Musser, J. M., Khariton, M., Li, P., Arendt, D., Quake, S. R. and Wang, B. (2021). Mapping single-cell atlases throughout Metazoa unravels cell type evolution. Elife 10:e66747.

Thiruchelvam, U., Dransfield, I., Saunders, P. T. K. and Critchley, H. O. D. (2013). The importance of the macro-phage within the human endometrium. J. Leukoc. Biol. 93, 217–225.

Thiruchelvam, U., Maybin, J. A., Armstrong, G. M., Greaves, E., Saunders, P. T. K. and Critchley, H. O. D. (2016). Cortisol regulates the paracrine action of macro-phages by inducing vasoactive gene expression in endometrial cells. J. Leukoc. Biol. 99, 1165–1171.

Traag, V. A., Waltman, L. and van Eck, N. J. (2019). From Louvain to Leiden: guaranteeing well-connected communities. Sci. Rep. 9, 5233.

Trapnell, C., Cacchiarelli, D., Grimsby, J., Pokharel, P., Li, S., Morse, M., Lennon, N. J., Livak, K. J., Mikkelsen, T. S. and Rinn, J. L. (2014). The dynamics and regulators of cell fate decisions are revealed by pseudotemporal ordering of single cells. Nat. Biotechnol. 32, 381–386.

Turco, M. Y., Gardner, L., Hughes, J., Cindrova-Davies, T., Gomez, M. J., Farrell, L., Hollinshead, M., Marsh, S. G. E., Brosens, J. J., Critchley, H. O., et al. (2017). Long-term, hormone-responsive organoid cultures of human endometrium in a chemically defined medium. Nat. Cell Biol. 19, 568–577.

Ulrich, N. D., Vargo, A., Ma, Q., Shen, Y.-C., Bazzano, D., Hannum, D. F., Gurczynski, S. J., Moore, B. B., Schon, S., Lieberman, R., et al. (2024). Cellular heterogeneity and dynamics of the human uterus in healthy premenopausal women. Proc. Natl. Acad. Sci. U. S. A. 121, e2404775121.

Virshup, I., Rybakov, S., Theis, F. J., Angerer, P. and Wolf, F. A. (2024). anndata: Access and store annotated data matrices. J. Open Source Softw. 9, 4371.

Virtanen, P., Gommers, R., Oliphant, T. E., Haberland, M., Reddy, T., Cournapeau, D., Burovski, E., Peterson, P., Weckesser, W., Bright, J., et al. (2020). SciPy 1.0: fundamental algorithms for scientific computing in Python. Nat. Methods 17, 261–272.

Wang, C., Zhao, M., Zhang, W.-Q., Huang, M.-Y., Zhu, C., He, J.-P. and Liu, J.-L. (2020a). Comparative analysis of mouse decidualization models at the molecular level. Genes (Basel) 11, 935.

Wang, W., Vilella, F., Alama, P., Moreno, I., Mignardi, M., Isakova, A., Pan, W., Simon, C. and Quake, S. R. (2020b). Single-cell transcriptomic atlas of the human endometrium during the menstrual cycle. Nat. Med. 26, 1644–1653.

Weiler, P. and Theis, F. J. (2026). CellRank: consistent and data view agnostic fate mapping for single-cell genomics. Nat. Protoc. 21, 3552–3579.

Weiler, P., Lange, M., Klein, M., Pe’er, D. and Theis, F. (2024). CellRank 2: unified fate mapping in multiview single-cell data. Nat. Methods 21, 1196–1205.

Winkler, I., Tolkachov, A., Lammers, F., Lacour, P., Daugelaite, K., Schneider, N., Koch, M.-L., Panten, J., Grünschläger, F., Poth, T., et al. (2024). The cycling and aging mouse female reproductive tract at single-cell resolution. Cell 187, 981–998.e25.

Wolf, F. A., Angerer, P. and Theis, F. J. (2018). SCANPY: large-scale single-cell gene expression data analysis. Genome Biol. 19, 15.

Wu, B., Chen, X., He, B., Liu, S., Li, Y., Wang, Q., Gao, H., Wang, S., Liu, J., Zhang, S., et al. (2014). ROS are critical for endometrial breakdown via NF-κB-COX-2 signaling in a female mouse menstrual-like model. Endocrinology 155, 3638–3648.

Xu, X. B., He, B. and Wang, J. D. (2007). Menstrual-like changes in mice are provoked through the pharmacologic withdrawal of progesterone using mifepristone following induction of decidualization. Hum. Reprod. 22, 3184–3191.

Yang, M., Ong, J., Meng, F., Zhang, F., Shen, H., Kitt, K., Liu, T., Tao, W. and Du, P. (2023). Spatiotemporal insight into early pregnancy governed by immune-featured stromal cells. Cell 186, 4271–4288.e24.

Yang, L., Yang, J., Li, Y., Su, Y., Ma, H., Cai, S., Mo, M., Ding, L., Huang, C., Liu, F., et al. (2026). An endometrial immune atlas across the female menstrual cycle reveals lymphoid aggregates compromise pregnancy. Nat. Commun. 17, 9981.

Yee, T. W. (2015). Vector generalized linear and additive models: With an implementation in R. 1st ed. New York, NY: Springer.

Yee, G. M. and Kennedy, T. G. (1991). Role of cyclic adenosine 3’,5’-monophosphate in mediating the effect of prostaglandin E2 on decidualization in vitro. Biol. Reprod. 45, 163–171.

Yu, D., Wong, Y.-M., Cheong, Y., Xia, E. and Li, T.-C. (2008). Asherman syndrome--one century later. Fertil. Steril. 89, 759–779.

Yuan, J., Deng, W., Cha, J., Sun, X., Borg, J.-P. and Dey, S. K. (2018). Tridimensional visualization reveals direct communication between the embryo and glands critical for implantation. Nat. Commun. 9, 603.

Zheng, G. X. Y., Terry, J. M., Belgrader, P., Ryvkin, P., Bent, Z. W., Wilson, R., Ziraldo, S. B., Wheeler, T. D., McDermott, G. P., Zhu, J., et al. (2017). Massively parallel digital transcriptional profiling of single cells. Nat. Commun. 8, 14049.

Zhu, H., Aryal, D. K., Olsen, R. H. J., Urban, D. J., Swearingen, A., Forbes, S., Roth, B. L. and Hochgeschwender, U. (2016). Cre-dependent DREADD (designer receptors exclusively activated by designer drugs) mice. Genesis 54, 439–446.

Zuo, R.-J., Gu, X.-W., Qi, Q.-R., Wang, T.-S., Zhao, X.-Y., Liu, J.-L. and Yang, Z.-M. (2015). Warburg-like glycolysis and lactate shuttle in mouse decidua during early pregnancy. J. Biol. Chem. 290, 21280–21291.

Zwierzynski, J. B., Moufarrej, M. N. and Red-Horse, K. (2026). 3D imaging of the pregnant uterus reveals an extensively invasive mouse placenta and early CXCL12-CXCR4 requirement. bioRxivorg 2026.04.06.716785.

