## Supplementary materials for "Transgenic menstruating mice reveal the spatial logic of pre-menstrual uterine differentiation"

*1 Department of Stem Cell and Regenerative Biology, Harvard University, Cambridge MA, 02138, USA*
*2 Department of Molecular and Cellular Biology, Harvard University, Cambridge MA, 02138, USA*
*3 Broad Institute, Cambridge MA 02142, USA*
*4 MRC Laboratory of Medical Sciences, London, UK*
*5 Biological and Biomedical Sciences Program, Harvard University, Cambridge MA, 02138, USA*
*6 Howard Hughes Medical Institute, Cambridge MA, 02138, USA*
*7 Department of Immunology, Harvard Medical School, Boston, MA, 02115, USA*
*8 Department of Systems Biology, Harvard University, Boston MA 02115, USA*
*9 Fundación Instituto Leloir, CABA, Argentina*
*10 Department of Microbiology and Immunology, Loyola University Chicago, Chicago IL, 60153, USA*
*11 Department of Obstetrics and Gynecology, University of Pennsylvania, Philadelphia PA 19104, USA*
*12 Harvard Stem Cell Institute, Harvard University, Cambridge MA 02138, USA*

### Supplementary Material

This file includes:
- Supplementary Text
- Figures S1 to S37

Other Supplementary Materials include:
- Supplementary Data S1-3

#### Supplementary Text

**Comparison of GqD and GsD X-Mens strains:**

***Cell types expressing the DREADDs:*** The GsD X-Mens strain combines the Cre-inducible $R26^{GsD}$ line (Akhmedov et al., 2017), together with the $Amhr2^{Cre}$ line (Jamin et al., 2002) to target endometrial fibroblast cells. Cre expression in these animals initiates in the genital ridges at gestational day 11.5. We used both $Amhr2^{Cre/+}$ x $R26^{GsD/+}$ and $Amhr2^{Cre/+}$ x $R26^{GsD/GsD}$ genotypes for our studies. The expression pattern of Amhr2 and the resulting recombination induced by this Cre line have been subject to ongoing debate (Ang et al., 2023). In our experiments, we consistently observed recombination in the endometrial fibroblasts, myometrium, and occasionally the endometrial epithelium (Figure S1 and Figure S20). To determine whether this occasional epithelial recombination could account for GsD-induced decidualization, we expressed GsD in the endometrial epithelium using the $Ltf^{iCre}$ line (Daikoku et al., 2014). These mice did not decidualize following pseudopregnancy and agonist treatment (Figure S2), indicating that elevation of epithelial cAMP is not sufficient to drive decidualization and that the epithelial component of $Amhr2^{Cre}$ recombination is therefore unlikely to underlie the GsD phenotype. This is distinct from the requirement for the luminal epithelium in licensing fibroblast decidualization, which we address in the main text (Figure 6F).

The GqD X-Mens strain combines the Cre-inducible $Tg^{GqD}$ (Zhu et al., 2016) with the $Ltf^{iCre}$ line. Cre expression initiates at puberty in these animals. $Ltf^{iCre/+}$ x $Tg^{GqD/0}$ animals were used in these experiments. $Ltf^{iCre}$ is reported to be active in the endometrial epithelium and in neutrophils (Daikoku et al., 2014), an expression pattern we confirmed in our analysis (Figure S1). Whether the neutrophil population contributes to the GqD phenotype was not directly tested.

In our experiments, expressing DREADDs with other drivers, including $Pgr^{Cre}$ (Yang et al., 2013), $Pdgfra^{Cre}$ (Roesch et al., 2008), $Pax8^{Cre}$ (Bouchard et al., 2002), $Cxcl15^{Cre}$ (Kelleher et al., 2025) and $Foxl2^{Cre}$ (Nicol et al., 2026) rendered animals moribund, likely reflecting DREADD expression in other organs; we did not investigate this further. Future development of alternative approaches, using different Cre lines, AAV-mediated DREADD delivery, or engineering improved DREADD proteins may further enhance the versatility and precision of this method.

***Biological considerations:*** GsD X-Mens initiate decidualization within the fibroblast compartment, paralleling the human cycle, in which a systemically delivered decidualization cue is received by the fibroblasts themselves. GqD X-Mens initiate the cascade in the epithelium, more closely mirroring mouse pregnancy, in which the signal from the implanting blastocyst is received by the luminal epithelium and relayed to the underlying fibroblasts (Ramathal et al., 2010). The two strains also differ in the second messenger that triggers the response. We show here that fibroblast cAMP, acting through PKA, is required for decidualization during pregnancy (Figure 1, E and F), establishing that the GsD trigger intersects the endogenous decidualization pathway. Whether calcium is likewise an endogenous mediator remains to be tested, as we did not perform an equivalent loss-of-function experiment.

GsD X-Mens mice do not produce live litters, undergoing pregnancy loss in late gestation regardless of exposure to agonist. This is potentially due to basal activity of the GsD receptor in the absence of agonist. Although we use a GFP-fused GsD with lower basal activation than the original GsD, activation is not completely absent (Nakajima et al., 2016). Implantation and early gestation are unaffected by menstruation: quantification of implantation sites at gestational day 9.5 showed comparable numbers in previously menstruated and uninduced GsD X-Mens animals (Figure S9), indicating that menstruation does not impede subsequent embryo implantation or early pregnancy.

Additionally, GsD X-Mens mice develop vaginal swelling after repeated rounds of menstruation (11 of 14 animals), which predominantly begins at the fourth induction cycle. This phenotype does not appear to affect the overall health of the animals and also occurred in animals given agonist repeatedly without pseudopregnancy (2 of 4 animals), indicating that it does not require decidualization or menstruation. Amhr2Cre is active throughout the Müllerian duct mesenchyme, which contributes to the cervix and vagina in addition to the uterus (Cunha et al., 2019; Jamin et al., 2002), so the phenotype may reflect GsD activation outside the uterine body; the small size of the agonist-only group precludes a firm conclusion.

GqD X-Mens mice are fertile regardless of exposure to agonist (Figure S9) and do not develop vaginal swelling after repeated induction, making them the preferred model for studies of fertility and repeated menstruation, notwithstanding their lower bleeding efficiency (see below).

***Phenotypic differences:*** GsD X-Mens mice exhibit a cystic morphology of their endometrial epithelial glands during decidualization that we did not observe in GqD X-Mens mice and that, to our knowledge, has not been reported in the oil-induced menstruation model (Figure S6). GsD X-Mens also exhibits more extensive remodeling of arterial architecture, beginning to resemble the early branching stages of spiral artery development in pregnancy (Figure 3G and H). GqD X-Mens mice undergo many of the same decidualization-associated arterial changes but do not progress to the same extent of branching. GqD X-Mens mice occasionally display a small amount of vaginal bleeding detectable by guaiac test shortly after induction (5 of 42 animals), often followed by a non-bleeding interval before menstruation begins; we did not observe this in GsD X-Mens mice. Finally, decidualization appears to progress more rapidly in GsD than in GqD X-Mens mice, based on changes in uterine volume over time, although we did not quantify this systematically.

***Practical considerations:*** Breeding of GqD X-Mens is enabled by the ability of *Ltf*$^{iCre}$ mice to be maintained as healthy, fertile homozygotes; note that the *Ltf*$^{iCre}$ line is a knockout-knockin, and *Ltf*$^{iCre/iCre}$ animals therefore lack *Ltf* expression. Although GqD is an unmapped transgene, and homozygous animals cannot be distinguished from hemizygotes by standard genotyping, homozygous founders can be identified by test crosses in which all progeny are GqD-positive. Breeding colonies can therefore be established in which 100% of female progeny are experimental. In contrast, *Amhr2*$^{Cre/Cre}$ males are unsuitable as breeders, as they retain Müllerian duct structures and are infertile, reflecting the requirement for *Amhr2* in Müllerian duct regression (Jamin et al., 2002). The most efficient cross for these animals is therefore *Amhr2*$^{Cre/+}$ males to *R26*$^{GsD/GsD}$ females, yielding 50% experimental female progeny.

***Induction protocol:*** Most of our experiments induced decidualization using a combination of water-soluble clozapine N-oxide (CNO) delivered in drinking water and five consecutive intraperitoneal injections of a mixture of three DREADD agonists (water-soluble CNO, deschloroclozapine, and compound 21; see Materials and Methods). This approach robustly induced decidualization in both strains, although GqD X-Mens mice displayed lower bleeding efficiency than GsD X-Mens mice (Figure 2E), potentially due to epithelial mosaicism of the LtfiCre line, which we did not test directly. We observed that delivery via drinking water or intraperitoneal injection alone, or reducing the number of injections, led to decreased bleeding efficiency, although we did not quantify this systematically. We also tested a single subcutaneous administration of deschloroclozapine, which efficiently induced the GsD X-Mens strain (Figure S7) but not the GqD X-Mens strain; further optimization would be required before using this route in GqD X-Mens mice.

**Comparison to other mouse menstruation models:**

All existing mouse models of menstruation combine progesterone fluctuation with a decidualization stimulus. Previous models use intrauterine oil or mechanical scratch as the stimulus (Rogers et al., 2025). Our approach instead uses cell type-specific chemogenetic elevation of cAMP or calcium.

***Progesterone control:*** A variety of methods for controlling progesterone fluctuation have been developed for oil-based menstruation models, reviewed in detail in (Rogers et al., 2025). In brief, one strategy is ovariectomy followed by exogenous hormone supplementation using repeated injections, a subcutaneous progesterone implant, or both. An abrupt progesterone decline is then achieved by stopping the injections or removing the implant. 4 hours from implant withdrawal, vaginal bleeding is detectable in approximately half of treated mice (Cousins et al., 2016). In this model, approximately 90% of mice are bleeding at 12 hours, and bleeding is complete by 24 hours (Cousins et al., 2016). An alternative approach is to elevate endogenous progesterone by mating with a vasectomized male, which induces pseudopregnancy, a state in which the progesterone-producing corpora lutea of the ovary persist for approximately eight days. Natural demise of the corpora lutea (luteolysis) drives a more gradual progesterone decline (Rudolph et al., 2012).

The choice of hormonal control method involves a trade-off between physiological fidelity and precision of timing. In the pseudopregnancy approach used here, progesterone declines through luteolysis, the same mechanism that terminates the human luteal phase. It is also non-surgical and compatible with future pregnancy. However, the timing of luteolysis varies between individuals, producing variation in the onset and duration of bleeding (Figure S10 and S11). We show that this variability can be substantially reduced by administration of the progesterone receptor antagonist mifepristone (Figure S11). However, systemic antagonism of progesterone signaling introduces potential confounds and mifepristone also antagonizes the glucocorticoid receptor (Moguilewsky and Philibert, 1984; Whirledge et al., 2015). Ovariectomy coupled with hormone supplementation, particularly the implant withdrawal method, triggers an abrupt fall in progesterone that yields more reproducible bleeding onset, and may therefore be preferable for studies requiring a defined start time, at the cost of surgery and of subsequent breeding. We anticipate that the range of progesterone control methods deployed in previous mouse models of menstruation will be compatible with X-Mens induction.

***Efficiency:*** Induction efficiencies are not directly comparable across models. Published oil efficiencies are reported as the proportion of animals with at least one decidualized horn, whereas we report the proportion of animals exhibiting detectable vaginal bleeding. In our studies, 93% of GsD X-Mens and 70% of GqD X-Mens animals bled following induction (Figure 2E). For reference, a recent study reported decidualization in at least one horn in 89% of animals after oil delivered by laparotomy, 83% by laparoscopy, and 75% transcervically (Peterse et al., 2018).

***Transcriptomic comparison:*** To compare decidualization between X-Mens and oil induction at the transcriptomic level, we used a published list of differentially expressed genes (DEGs) between oil-stimulated and unstimulated uterine horns of pseudopregnant mice (Wang et al., 2020). We compared these DEGs with those identified between pseudobulked decidual and fibroblast nuclei in X-Mens mice (see Materials and Methods). Because technical differences between datasets, including sequencing depth, can affect which genes are detected as differentially expressed, we restricted the analysis to genes called as DEGs in both datasets (1,350 genes) and asked whether they changed in the same direction. Of these shared DEGs, 1,238 (92%) changed in the same direction in both datasets (555 up and 683 down), and 112 (8%) changed in opposite directions (Figure S35). The two datasets are not strictly equivalent: the oil comparison is between bulk uterine horns, so its DEGs reflect both cell-intrinsic changes and shifts in cell composition, whereas the X-Mens comparison is between annotated cell populations.

***Cell composition:***

<u>Fibroblasts:</u> Two studies from Kirkwood et al. reported five fibroblast populations (F1-5) identified from single-cell RNA sequencing of endometrial stromal cells (Kirkwood et al., 2021; Kirkwood et al., 2022). In these studies, F1-3 were identified in estrus stage animals. F4 and F5 were identified in an oil-induced menstruation model that achieves hormonal control by ovariectomy followed by hormone injections and insertion and removal of a progesterone pellet. Animals were harvested 24 and 48 hours after progesterone implant removal. We examined markers of the F1-F5 populations in our dataset (Figure S36). These analyses identified two putative shared clusters: markers of F3 were expressed in eFib1 in our dataset, and both studies report these as the outermost fibroblast cluster. F5, reported as an apoptotic cluster, may correspond to decAp in our dataset. We did not detect clear correspondence for the remaining clusters.

Several differences between the studies may contribute. First, samples were collected at distinct timepoints. X-Mens samples were collected during pseudopregnancy and during decidualization and early bleeding. Kirkwood et al. sampled animals during estrus and during active bleeding and repair (24 or 48 hours following progesterone withdrawal); correspondingly, decidual cells were not reported in their datasets. Importantly, we performed highly stringent filtering of fibroblast cells during pre-processing, which was designed to yield a high-purity population of fibroblasts and decidual cells to study decidual progression (see Methods). To avoid potentially confounding signals, we deliberately excluded cells that co-expressed markers of fibroblasts and markers of another lineage. Such cells can represent technical artifacts such as persistent doublets, but some represent true cell types. This stringent filtering step likely explains why we do not detect fibroblast population F4, which expresses both fibroblast and epithelial markers. Similarly, we anticipate that our approach would exclude cells that co-express fibroblast and immune markers (Ulrich et al., 2024). Although these populations fall outside the scope of our study, they may be of interest to others, and our raw data are available for further analysis.

<u>Immune cells:</u> Ainslie et al. (Ainslie et al., 2026) performed single cell RNA sequencing and flow cytome-

try of an enriched immune population in cycling animals and in an oil-induced menstruation model that achieved hormonal control through ovariectomy, hormone injections, and insertion and removal of a progesterone pellet. Animals were harvested 0, 12, 24 and 48 hours after progesterone implant removal. Comparing their datasets with the X-Mens dataset we detected some similar patterns, including decreasing proportions of uNK cells during the progression to menstruation. However, the oil-menstruation model is dominated by neutrophils at 12 and 24 hours after progesterone pellet withdrawal based on both single cell transcriptomics and flow cytometry, whereas neutrophils represent a small fraction of our single nucleus transcriptomic dataset at any time point, albeit with a modest increase in the menstrual (4.5 dpi) time point. Technical considerations may contribute to apparent differences, including different timepoints for sample collection and methods of sample processing. Cell composition may also be affected by the introduction of oil within the uterus or DREADD agonist into the circulation.

***Practical considerations:*** X-Mens induces decidualization non-surgically. Non-surgical induction enables repeated induction in the same animal, and avoids the confounding effects of analgesia, which can alter the inflammatory and prostaglandin signaling central to menstruation (Bofill Rodriguez et al., 2019). By contrast, uterine scratch requires abdominal surgery, and intrauterine oil is administered either surgically or transcervically. The route also determines which horns are induced: transcervical oil targets one of the two uterine horns at random, surgical oil can target one or both, and X-Mens induces both. Single-horn induction provides a contralateral control within the same animal, which X-Mens cannot offer; conversely, X-Mens yields induced tissue from both horns and supports paired comparisons across successive cycles in the same animal.

X-Mens is currently limited by its reliance on Cre/lox. Because the Cre driver is committed to DREADD expression, X-Mens cannot readily be combined with other Cre-dependent tools, such as Cre-based lineage tracing. Oil induction carries no such constraint and is readily combined with lineage tracing, an approach we have used for studies of post-menstrual regeneration (Ang et al., 2026). AAV-mediated DREADD delivery, or the use of orthogonal recombinase systems (Dre/rox or Flp/FRT), would expand the genetic toolkit compatible with X-Mens.

Biological considerations: The morphological consequences of decidualization differ between oil induction and X-Mens. Oil injection produces uneven decidualization along the length of the horn ((Kumar et al., 2025; Ledford et al., 1976; Peterse et al., 2018) and references therein), and we and others have observed multiple discrete decidual masses within a single oil-induced horn (Ang et al., 2026). Decidualization in X-Mens horns appears more uniform, although we did not quantify this. Separately, whether any mouse model of menstruation recapitulates the focal, asynchronous shedding characteristic of human menstruation (Garry et al., 2009) remains unclear.

The choice of model also depends on the biological question. X-Mens uses cell type-specific elevation of defined second messengers to induce decidualization. Our study provides an example of how this approach enables biological discovery. Had we only observed decidualization initiating adjacent to the lumen following oil-induction, or in GqD X-Mens, it would not have been informative: in both cases the stimulus is delivered into the lumen, so a luminal origin is expected. In GsD X-Mens, by contrast, GsD is expressed across the fibroblast population and the agonist is delivered systemically, so the source of the stimulus could not be the source of the pattern. This revealed a second spatial constraint acting downstream of cAMP elevation, which we traced to the luminal epithelium (Figure 6F).

**Comparison to human systems:**

Over a century of study of menstruation in humans and other primates has established a detailed description of this process. These studies use biopsies, or uteri recovered from hysterectomies or deceased donors, for histological and, more recently, sequencing-based analyses. This work provides the essential benchmarks against which the X-Mens model can be assessed.

Our analyses indicate strong similarity between X-Mens and humans in cell identity, cell organization, and transcriptional change within the stromal compartment (Figs. 5 and 6). To extend this comparison to previously reported cell clusters, we examined a recently described population of human peripheral fibroblasts marked by the gene *C7* (Garcia-Alonso et al., 2021), defined as subtype Stro-7 in a 20-donor human uterus atlas generated by Ulrich et al. (Ulrich et al., 2024). We generated a module score representing the top 50 marker genes of the Stro-7 cluster and scored each cell in our human dataset. This analysis revealed that the myoFib cluster corresponded most closely with Stro-7. Similarly, the cells with the highest gene expression similarity to Stro-7 localized to the outermost fibroblasts, corresponding predominantly, though not exclusively, to the myometrium (Figure S28). Ulrich et al. similarly noted that Stro-7 cells are enriched in myometrial preparations compared to endometrial preparations. To identify the corresponding cluster in our mouse dataset, we transferred the Stro-7 module score from the human cells to their mouse neighbors in the SAMap embedding. Spatial mapping revealed that these cells corresponded to the most peripheral mouse fibroblasts (Figure S28). Together these data indicate that the identity and position of this peripheral fibroblast population is conserved between species.

We also analyzed the immune compartment in our human samples (Figure S37), which revealed broadly similar populations of uNK cells, macrophage/myeloid cells, and lymphocytes in the two species. Humans had higher proportions of lymphocytes and lower proportions of macrophage/myeloid cells than X-Mens; however, given the two human donors analyzed, these proportional differences should be treated as provisional. Both species contained conventional granzyme B-low uNK cells and acquired a cytotoxic uNK population during menstruation. Surveillance of decidual senescent cells by uNK cells has been implicated in menstrual shedding in humans (Brighton et al., 2017). Our identification of senescent cells within the most mature decidual population (dec3) in X-Mens mice (Figure 4G) raises the possibility of similar mechanisms operating in this model.

#### References


**Ainslie, R. J., Kirkwood, P. M., Li, H., Gerasimou, S., Saunders, P. T. K., Simitsidellis, I. and Gibson, D. A.** (2026). Multimodal profiling of repair-associated immune cells in a mouse model of menstruation. J. Reprod. Immunol. 178, 104981.

**Akhmedov, D., Mendoza-Rodriguez, M. G., Rajendran, K., Rossi, M., Wess, J. and Berdeaux, R.** (2017). Gs-DREADD Knock-In Mice for Tissue-Specific, Temporal Stimulation of Cyclic AMP Signaling. Mol. Cell. Biol. 37,.

**Ang, C. J., Skokan, T. D. and McKinley, K. L.** (2023). Mechanisms of regeneration and fibrosis in the endometrium. Annu. Rev. Cell Dev. Biol. 39, 197–221.

**Ang, C. J., Gable, J. L. R., Lyons, K. C., Whelan, E. M., Çevrim, Ç., Skokan, T. D., Bennetts, S. G., Manetta, B. D., Kaage, A. M., Mopure, D., et al.** (2026). Luminal epithelium remodeling underlies endometrial regeneration during menstruation and pregnancy. bioRxivorg 2026.03.08.710375.

**Bofill Rodriguez, M., Lethaby, A. and Farquhar, C.** (2019). Non-steroidal anti-inflammatory drugs for heavy menstrual bleeding. Cochrane Database Syst. Rev. 9, CD000400.

**Bouchard, M., Souabni, A., Mandler, M., Neubüser, A. and Busslinger, M.** (2002). Nephric lineage specification by Pax2 and Pax8. Genes Dev. 16, 2958–2970.

**Brighton, P. J., Maruyama, Y., Fishwick, K., Vrljicak, P., Tewary, S., Fujihara, R., Muter, J., Lucas, E. S., Yamada, T., Woods, L., et al.** (2017). Clearance of senescent decidual cells by uterine natural killer cells in cycling human endometrium. Elife 6, e31274.

**Cousins, F. L., Kirkwood, P. M., Murray, A. A., Collins, F., Gibson, D. A. and Saunders, P. T. K.** (2016). Androgens regulate scarless repair of the endometrial "wound" in a mouse model of menstruation. FASEB J. 30, 2802–2811.

**Cunha, G. R., Sinclair, A., Ricke, W. A., Robboy, S. J., Cao, M. and Baskin, L. S.** (2019). Reproductive tract biology: Of mice and men. Differentiation 110, 49–63.

**Daikoku, T., Ogawa, Y., Terakawa, J., Ogawa, A., DeFalco, T. and Dey, S. K.** (2014). Lactoferrin-iCre: a new mouse line to study uterine epithelial gene function. Endocrinology 155, 2718–2724.

**Garcia-Alonso, L., Handfield, L.-F., Roberts, K., Nikolakopoulou, K., Fernando, R. C., Gardner, L., Woodhams, B., Arutyunyan, A., Polanski, K., Hoo, R., et al.** (2021). Mapping the temporal and spatial dynamics of the human endometrium in vivo and in vitro. Nat. Genet. 53, 1698–1711.

**Garry, R., Hart, R., Karthigasu, K. A. and Burke, C.** (2009). A re-appraisal of the morphological changes within the endometrium during menstruation: a hysteroscopic, histological and scanning electron microscopic study. Hum. Reprod. 24, 1393–1401.

**Jamin, S. P., Arango, N. A., Mishina, Y., Hanks, M. C. and Behringer, R. R.** (2002). Requirement of Bmpr1a for Müllerian duct regression during male sexual development. Nat. Genet. 32, 408–410.

**Kelleher, A. M., Kim, H. I., Bayammagari, G. S., Davis, D. J. and Spencer, T. E.** (2025). A Cxcl15 Cre recombinase mouse model useful to study gland development in the uterus. Genesis 63,.

**Kirkwood, P. M., Gibson, D. A., Smith, J. R., Wilson-Kanamori, J. R., Kelepouri, O., Esnal-Zufiaurre, A., Dobie, R., Henderson, N. C. and Saunders, P. T. K.** (2021). Single-cell RNA sequencing redefines the mesenchymal cell landscape of mouse endometrium. FASEB J. 35,.

**Kirkwood, P. M., Gibson, D. A., Shaw, I., Dobie, R., Kelepouri, O., Henderson, N. C. and Saunders, P. T. K.** (2022). Single-cell RNA sequencing and lineage tracing confirm mesenchyme to epithelial transformation (MET) contributes to repair of the endometrium at menstruation. Elife 11, e77663.

**Kraskov, A., Stögbauer, H. and Grassberger, P.** (2004). Estimating mutual information. Phys. Rev. E Stat. Nonlin. Soft Matter Phys. 69, 066138.

**Kumar, H. R., Massri, N., Bhurke, A. V., Kapur, A., Gadhiya, P. and Arora, R.** (2025). Murine implantation chamber formation precedes natural and artificial decidualization. bioRxivorg 2025.12.30.697086.

**Ledford, B. E., Rankin, J. C., Markwald, R. R. and Baggett, B.** (1976). Biochemical and morphological changes following artificially stimulated decidualization in the mouse uterus. Biol. Reprod. 15, 529–535.

**Li, R., Wang, T.-Y., Xu, X., Emery, O. M., Yi, M., Wu, S.-P. and DeMayo, F. J.** (2022). Spatial transcriptomic profiles of mouse uterine microenvironments at pregnancy day 7.5. Biol. Reprod. 107, 529–545.

**Moguilewsky, M. and Philibert, D.** (1984). RU 38486: potent antiglucocorticoid activity correlated with strong binding to the cytosolic glucocorticoid receptor followed by an impaired activation. J. Steroid Biochem. 20, 271–276.

**Nakajima, K.-I., Cui, Z., Li, C., Meister, J., Cui, Y., Fu, O., Smith, A. S., Jain, S., Lowell, B. B., Krashes, M. J., et al.** (2016). Gs-coupled GPCR signalling in AgRP neurons triggers sustained increase in food intake. Nat. Commun. 7, 10268.

**Nicol, B., Gruzdev, A., Amato, C., Liu, C., Scott, G. J., Hagler, T. B., Ray, M. K. and Yao, H. H.-C.** (2026). A new constitutive Foxl2-Cre mouse model reveals Foxl2-derived cell fate in reproductive and non-reproductive tissues. Biol. Reprod. 115, 832–849.

**Pedregosa, F., Varoquaux, G., Gramfort, A., Michel, V., Thirion, B., Grisel, O., Blondel, M., Louppe, G., Prettenhofer, P., Weiss, R., et al.** (2011). Scikit-learn: Machine Learning in Python. J. Mach. Learn. Res. 12, 2825–2830.

**Peterse, D., Clercq, K. D., Goossens, C., Binda, M. M., F O, D., Saunders, P., Vriens, J., Fassbender, A. and D'Hooghe, T. M.** (2018). Optimization of endometrial decidualization in the menstruating mouse model for preclinical endometriosis research. Reprod. Sci. 25, 1577–1588.

**Ramathal, C. Y., Bagchi, I. C., Taylor, R. N. and Bagchi, M. K.** (2010). Endometrial decidualization: of mice and men. Semin. Reprod. Med. 28, 17–26.

**Roesch, K., Jadhav, A. P., Trimarchi, J. M., Stadler, M. B., Roska, B., Sun, B. B. and Cepko, C. L.** (2008). The transcriptome of retinal Müller glial cells. J. Comp. Neurol. 509, 225–238.

**Rogers, L. M., Lash, G. E., Anderson, G. M. and Girling, J. E.** (2025). Modelling menstruation in the common mouse: a narrative review. Reprod. Fertil. Dev. 37,.

**Rudolph, M., Döcke, W.-D., Müller, A., Menning, A., Röse, L., Zollner, T. M. and Gashaw, I.** (2012). Induction of overt menstruation in intact mice. PLoS One 7, e32922.

**Ulrich, N. D., Vargo, A., Ma, Q., Shen, Y.-C., Bazzano, D., Hannum, D. F., Gurczynski, S. J., Moore, B. B., Schon, S., Lieberman, R., et al.** (2024). Cellular heterogeneity and dynamics of the human uterus in healthy premenopausal women. Proc. Natl. Acad. Sci. U. S. A. 121, e2404775121.

**Wang, C., Zhao, M., Zhang, W.-Q., Huang, M.-Y., Zhu, C., He, J.-P. and Liu, J.-L.** (2020). Comparative analysis of mouse decidualization models at the molecular level. Genes (Basel) 11, 935.

**Whirledge, S. D., Oakley, R. H., Myers, P. H., Lydon, J. P., DeMayo, F. and Cidlowski, J. A.** (2015). Uterine glucocorticoid receptors are critical for fertility in mice through control of embryo implantation and decidualization. Proc. Natl. Acad. Sci. U. S. A. 112, 15166–15171.

**Yang, C. F., Chiang, M. C., Gray, D. C., Prabhakaran, M., Alvarado, M., Juntti, S. A., Unger, E. K., Wells, J. A. and Shah, N. M.** (2013). Sexually dimorphic neurons in the ventromedial hypothalamus govern mating in both sexes and aggression in males. Cell 153, 896–909.

**Zhu, H., Aryal, D. K., Olsen, R. H. J., Urban, D. J., Swearingen, A., Forbes, S., Roth, B. L. and Hochgeschwender, U.** (2016). Cre-dependent DREADD (designer receptors exclusively activated by designer drugs) mice. Genesis 54, 439–446.

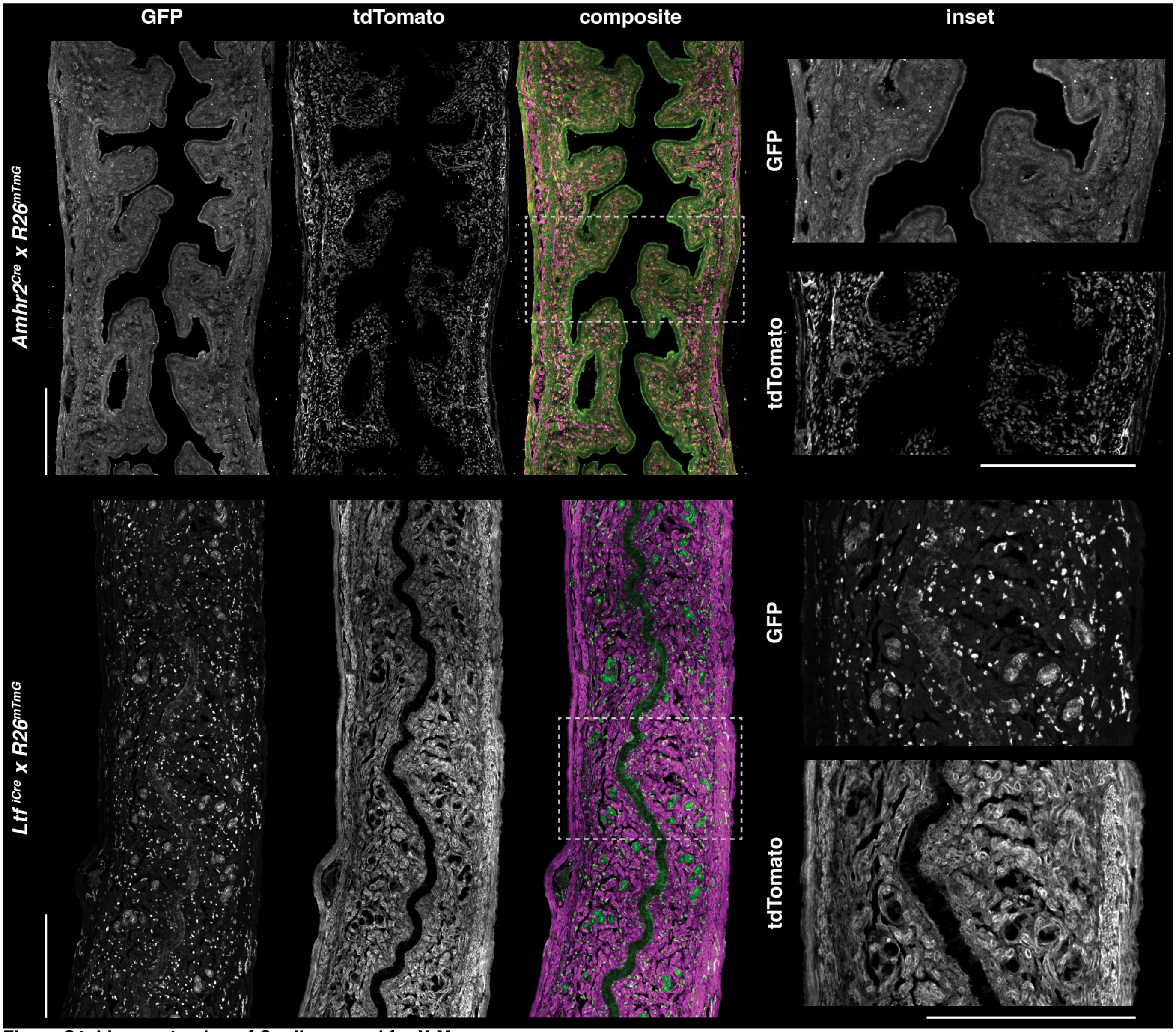


**Figure S1. Lineage tracing of Cre lines used for X-Mens.**
Immunofluorescence images of uterine longitudinal sections from Amhr2Cre (top) and LtfiCre (bottom) mice crossed with R26mTmG reporter mice. Cre-positive cells are labeled with membrane-bound GFP and Cre-negative cells express membrane-bound tdTomato. Individual channels are shown in grayscale, and composite images show the merged channels with GFP in green and tdTomato in magenta. Dashed boxes on the composite images indicate the regions shown at higher magnification in the insets. Scale bars: 500 $\mu$m.

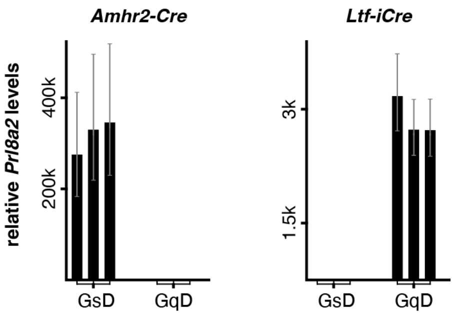


**Figure S2. Compartment specificity of chemogenetic *Prl8a2* induction.**
Relative *Prl8a2* mRNA levels in pseudopregnant *Amhr2*$^{Cre}$ (left) and *Ltf* $^{iCre}$ (right) mice carrying either the Gs-DREADD (GsD) or the Gq-DREADD (GqD), following agonist administration. Levels are normalized to uninduced controls. Each bar is the relative expression for one animal, with error bars showing the propagated standard deviation of technical replicates. For *Ltf* $^{iCre}$ *x Tg* $^{GqD}$, only animals with visible decidualization were analyzed, unlike Figure 1C, which includes all induced animals.

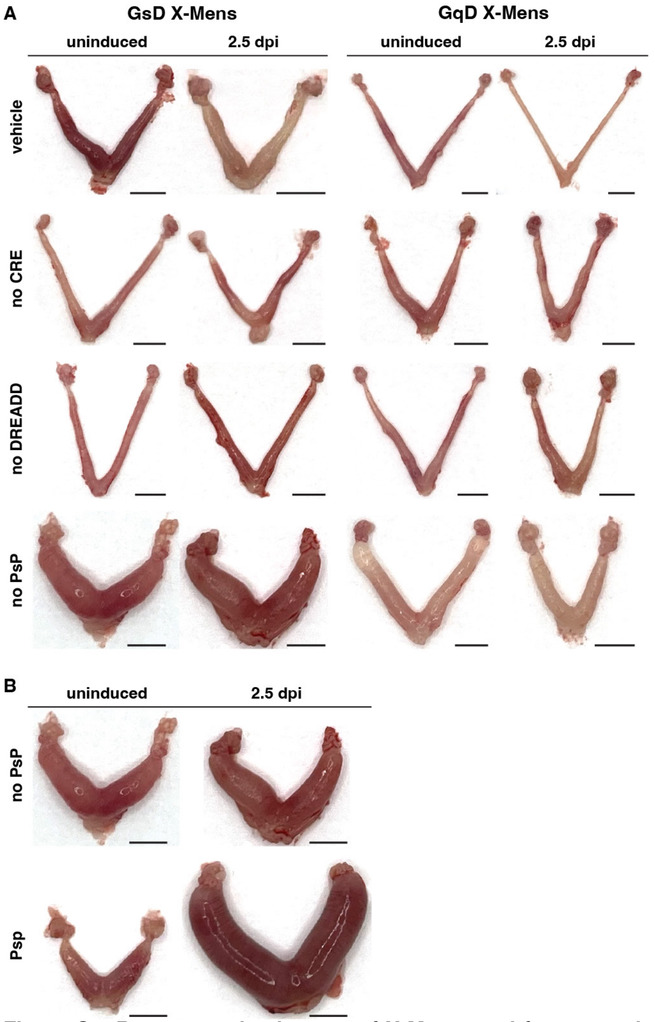


**Figure S3. Representative images of X-Mens uteri from experimental controls.**
**A.** Top row: animals receiving vehicle only; second row: genetic controls lacking the Cre allele; third row: genetic controls lacking the DREADD allele; bottom row: animals carrying both alleles induced with agonist but without pseudopregnancy (PsP). **B.** Uteri from GsD X-Mens animals that received agonist in the absence (top) or presence of pseudopregnancy (bottom), shown to scale to visualize that the size difference upon agonist delivery is unique to pseudopregnant animals. The no Psp uteri are reproduced from (A). The images of the Psp uteri are the same as those shown in Figure 1D. Scale bars: 5 mm. dpi: days post-induction.

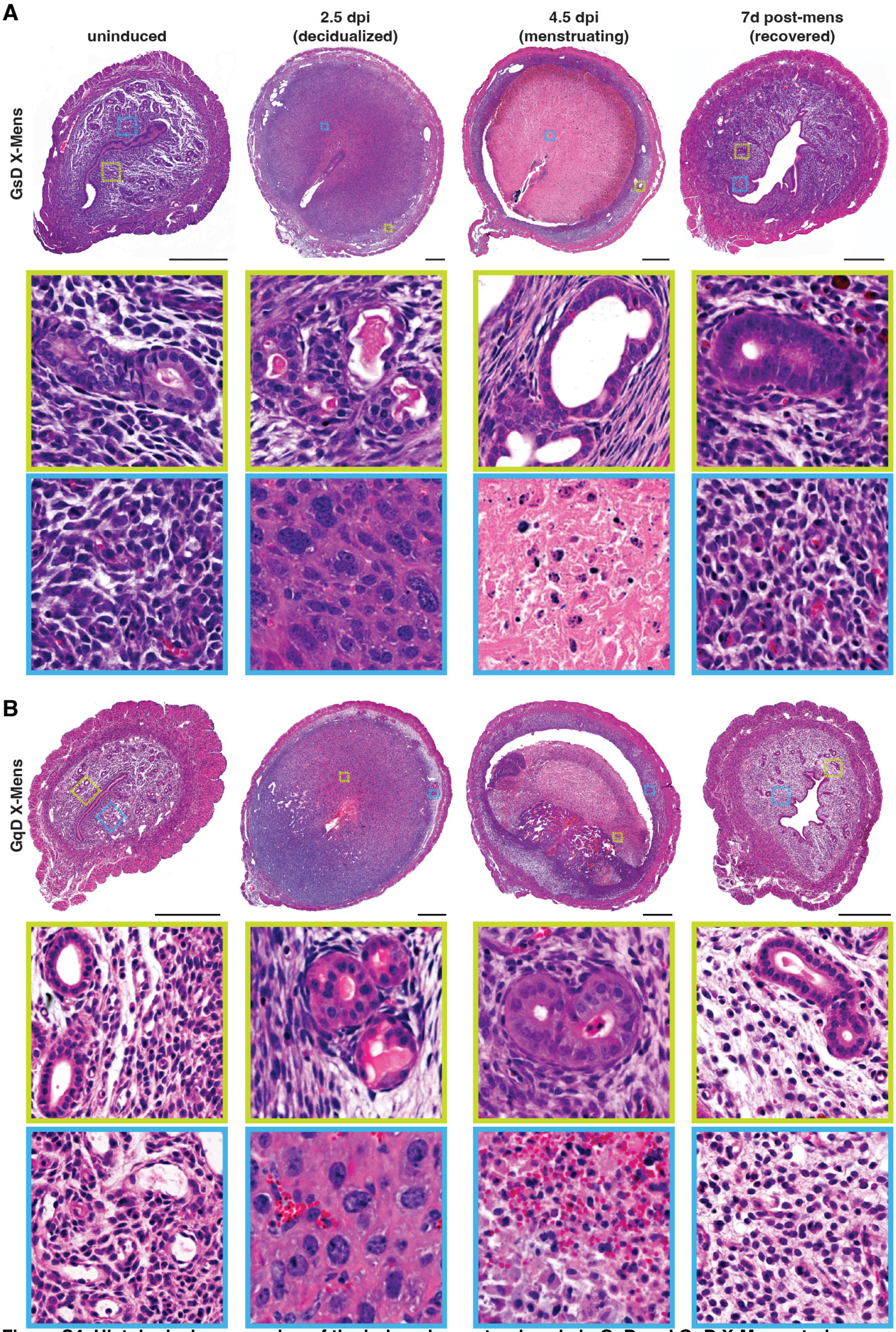


**Figure S4. Histological progression of the induced menstrual cycle in GsD and GqD X-Mens uteri.**
**A.** Hematoxylin and eosin (H&E) staining of GsD X-Mens uterine cross sections at the indicated stages: uninduced, 2.5 dpi (decidualized), 4.5 dpi (menstruating) and 7 days post-menstruation (recovered). The cross-sections in the top row are the same sections shown for GsD X-Mens in Figure 1D. Middle and bottom rows, magnified views of the regions boxed above, showing the glandular compartment (yellow) and the stroma (cyan). Each inset covers a 150 x 150 $\mu$m area. **B.** H&E staining of GqD X-Mens uteri at the stages indicated in panel A. The cross-sections in the top row are the same sections shown for GqD X-Mens in Figure 1D. Scale bars 500 $\mu$m. dpi: days post-induction.

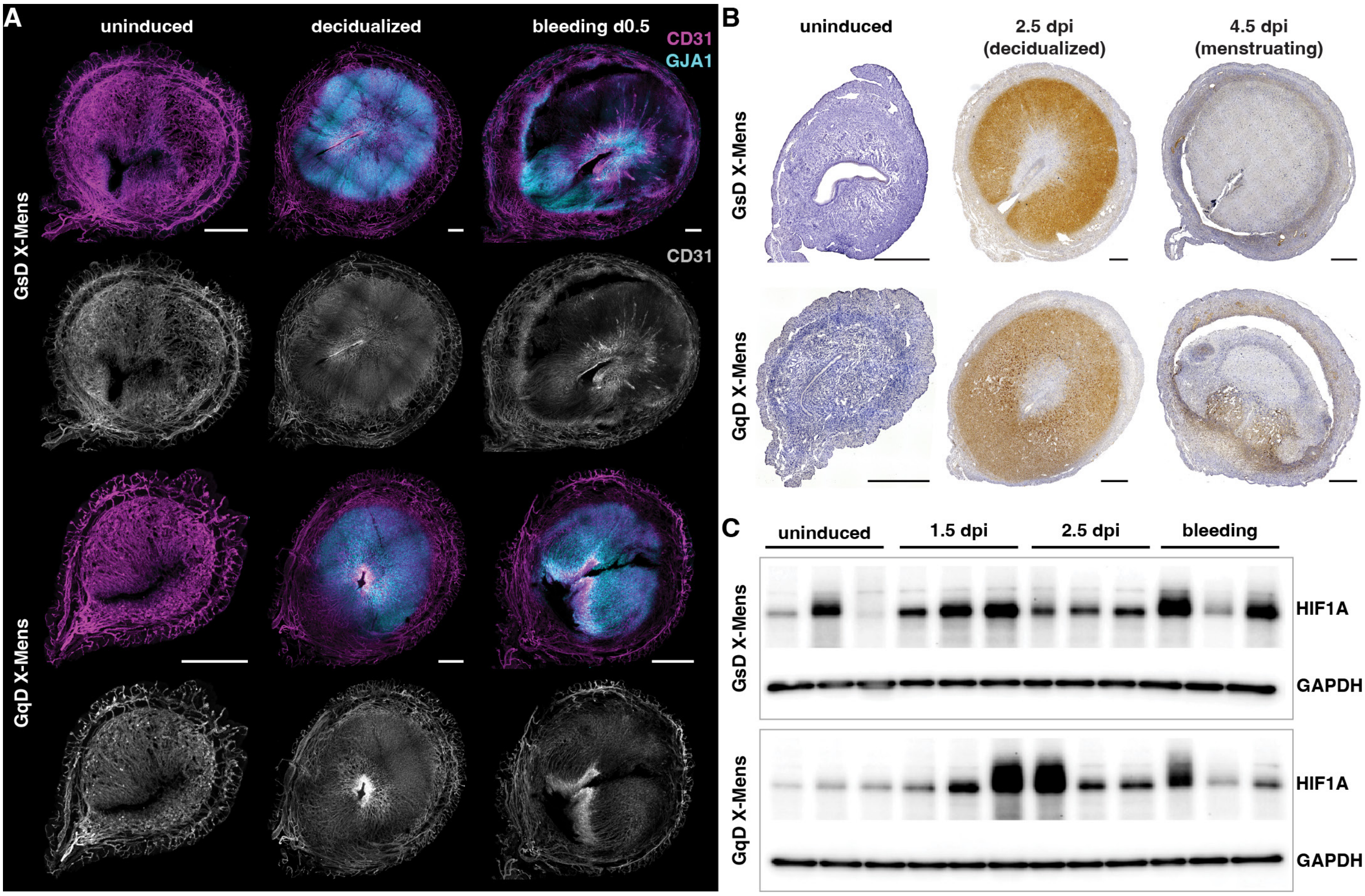


**Figure S5. Vascular and hypoxia markers in the endometrium of menstruating X-Mens mice.**
**A.** Representative immunofluorescence images of uterine cross sections from GsD (top) and GqD (bottom) X-Mens mice at the indicated stages, stained for CD31 (endothelial cells; magenta) and GJA1 (decidual cells; cyan). Grayscale images below each composite show the CD31 channel alone. Scale bars: 500 $\mu$m. Imaging settings, brightness, and contrast were adjusted independently to enable visualization of key structures. **B.** Immunohistochemical staining for carbonic anhydrase 9 (CA9, brown), a Hif1α-responsive gene, in uterine cross sections from GsD (top) and GqD (bottom) X-Mens mice at the indicated stages; nuclei are counterstained with hematoxylin (purple). **C.** Western blot of HIF1A and GAPDH in whole uterine lysates from GsD (top) and GqD (bottom) X-Mens mice at the indicated stages. Each lane represents one animal. dpi: days post-induction.

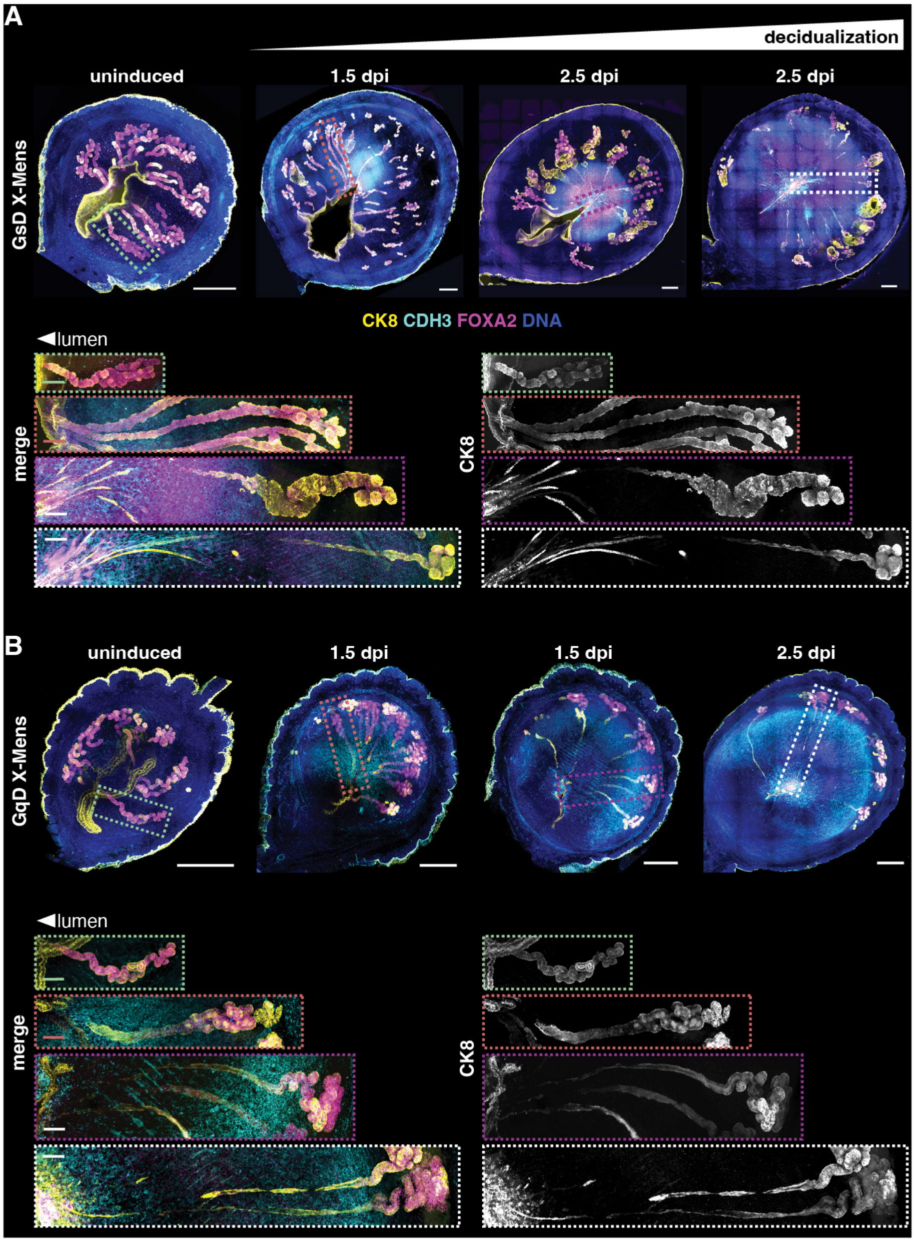


**Figure S6. Displacement of endometrial epithelial glands during decidualization in X-Mens mice.**
Immunofluorescence images of 200 $\mu$m thick uterine cross sections from **A.** GsD and **B.** GqD X-Mens mice, stained for epithelium (CK8, yellow), decidual cells (CDH3, cyan), glandular epithelium (FOXA2, magenta), and DNA (DAPI, blue). Sections are ordered by increasing extent of decidualization based on the size of the CDH3-positive decidual zone. Two 2.5 dpi samples are shown that exhibited differing degrees of decidualization at harvest. Dotted boxes on each section indicate the regions shown below at higher magnification, as merged channels (left) and the CK8 channel alone (right); box colors correspond between the section and its inset. Insets are shown on the same scale, and oriented with the uterine lumen to the left. Scale bars: 500 $\mu$m (sections) and 50 $\mu$m (insets). dpi: days post-induction.

**Figure S7. Comparison of agonist delivery protocols in X-Mens mice.**
Bleeding efficiency in GsD (left) and GqD (right) X-Mens mice following induction with CNO delivered in drinking water combined with five injections of a cocktail of DREADD agonists (black), or as a single injection of deschloroclozapine (Dcz, gray). n indicates the number of animals per group. The bleeding efficiency of animals administered water and five injections (black bars) is reproduced from data also shown in Figure 2E.

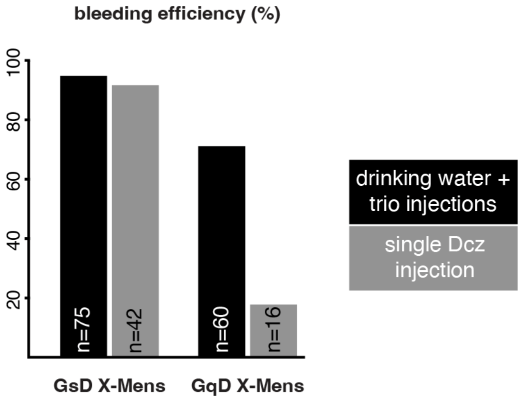

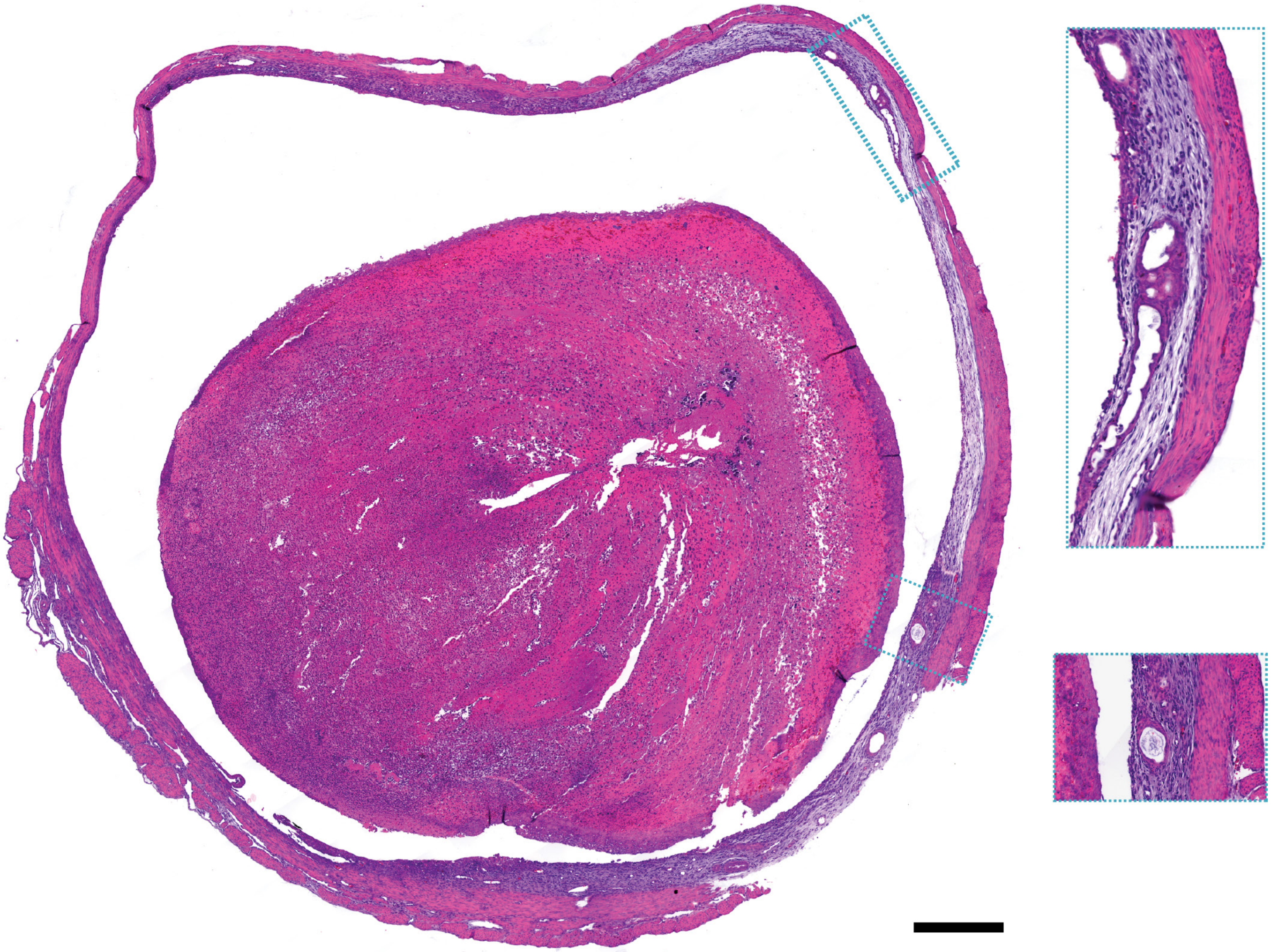

**Figure S8. A basalis-like layer in X-Mens mice contains non-decidual endometrial tissue and glands.**
H&E staining of an X-Mens uterus cross-section, harvested during menstruation. The inner functionalis-like layer is entirely detached from the outer basalis. Two inset images show glands and undifferentiated endometrial tissue residing in the basalis. Scale bar: 500$\mu$m.

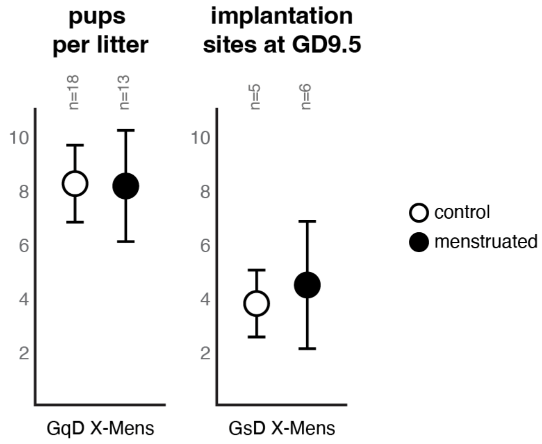


**Figure S9. Pregnancy outcomes in X-Mens mice.**
Left: the number of pups per litter in menstruated and vehicle-treated (control) GqD X-Mens mice. Right: the number of implantation sites at gestational day 9.5 (GD9.5) in menstruated and vehicle-treated (control) GsD X-Mens mice. Implantation sites were quantified because GsD-X-Mens mice experience late-gestation miscarriages regardless of exposure to DREADD agonist, precluding study of litter size (see supplementary text). Data are presented as mean ± 95% confidence interval. n indicates the number of dams per group.

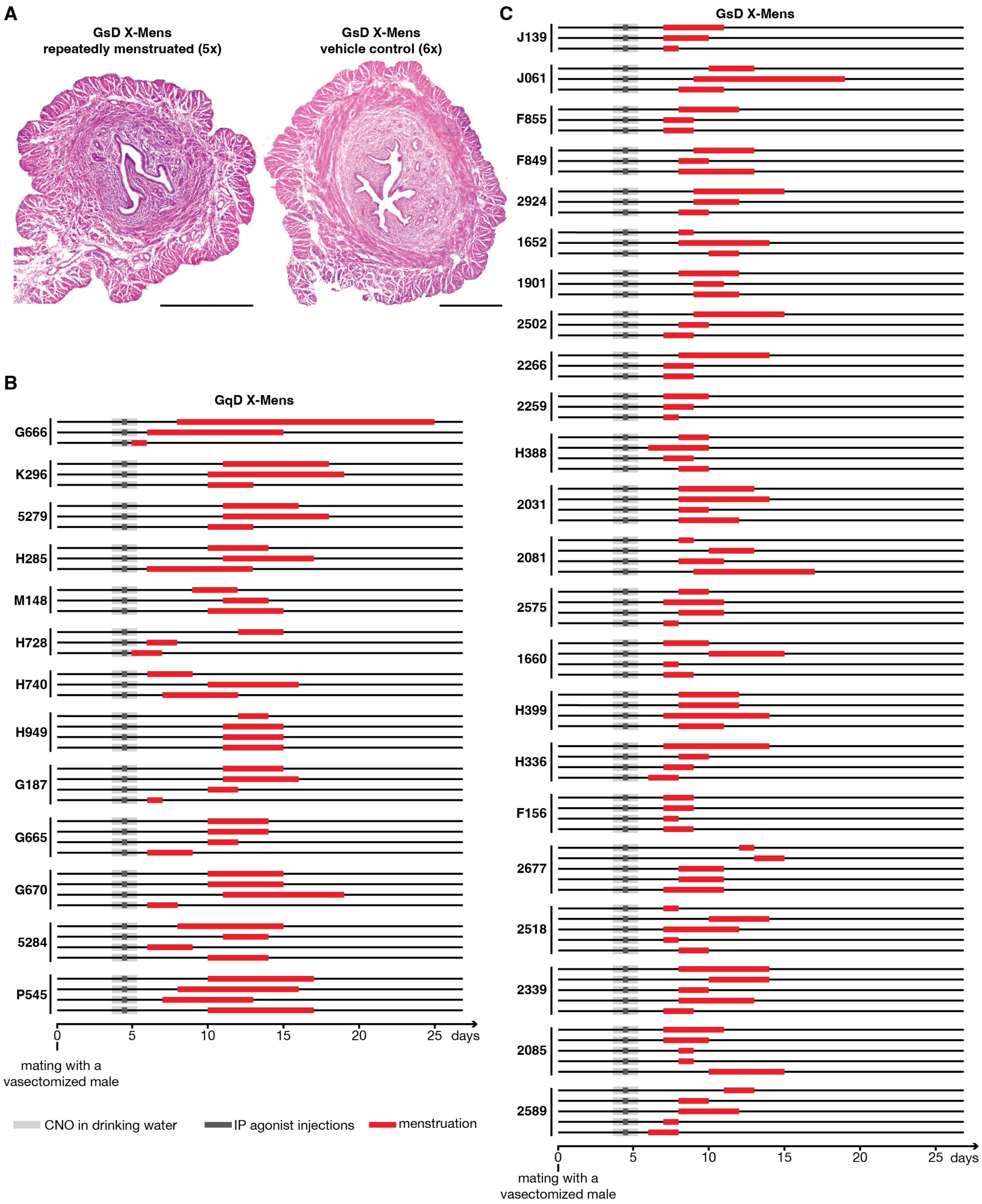


**Figure S10. Repeated menstruation does not alter uterine morphology, and bleeding episodes vary within and between animals.**
**A.** H&E staining of uterine cross sections from a GsD X-Mens mouse that underwent five rounds of menstruation (left) and a GsD X-Mens mouse that underwent six rounds of the induction protocol with vehicle in place of agonist (right). Scale bars: 500 $\mu$m. **B-C.** Individual bleeding episodes for each GqD X-Mens mouse (B; n = 13), and each GsD X-Mens mouse (C; n = 22), all of which underwent repeated menstruation. Animal ID is shown on the left; within each animal, the bottom row is the first menstruation and the top row the last. Day 0 is the day a copulatory plug was found after mating with a vasectomized male. Light gray indicates CNO in the drinking water, dark gray indicates intraperitoneal agonist injections, and red indicates menstrual bleeding. The bleeding trajectory of two animals from each strain are reproduced in Figure 2D.

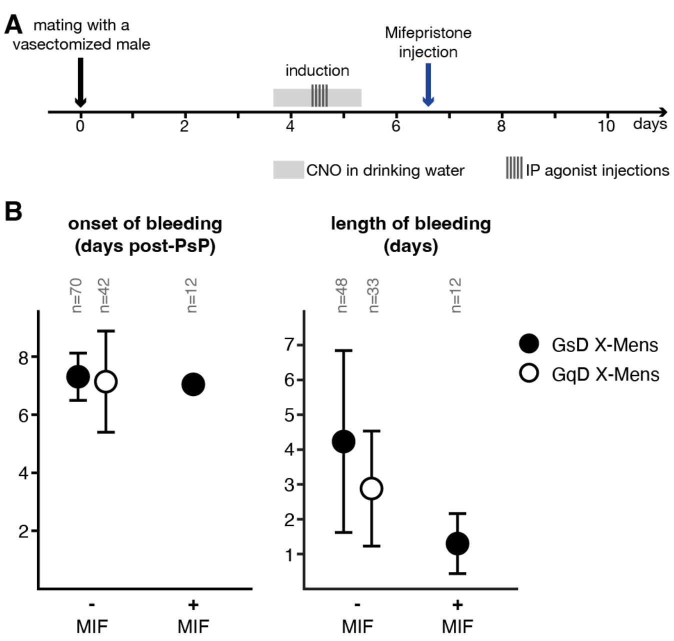


**Figure S11. Induction protocol and variability in onset and duration of bleeding in X-Mens models.**
**A.** Schematic of the X-Mens induction protocol with optional mifepristone injection step. Agonist is delivered as CNO in drinking water (light gray) combined with intraperitoneal (IP) agonist injections (vertical bars; see supplementary text). **B.** Onset (left) and length (right) of bleeding in GsD and GqD X-Mens mice, with and without MIF (mifepristone) injection (mean ± 95% confidence interval). MIF was administered to GsD X-Mens mice only. n indicates the number of bleeding episodes per group. PsP: pseudopregnancy.

**A**

| timepoint | section thickness | #10X lanes | #cells | spatially mapped (%) |
|---|---|---|---|---|
| uninduced | 2x30$\mu$m | 2 | 16,668 | 42.1 |
| 1.5 dpi early | 20$\mu$m | 1 | 3,146 | 55.6 |
| 1.5 dpi late | 20$\mu$m | 2 | 13,272 | 53.2 |
| 2.5 dpi | 20$\mu$m | 1 | 4,169 | 61.5 |
| 3.5 dpi | 20$\mu$m | 2 | 12,074 | 62.2 |
| 4.5 dpi (mens) | 20$\mu$m | 1 | 7,914 | 74.1 |

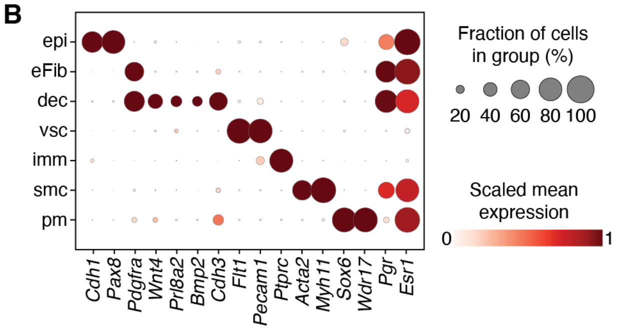


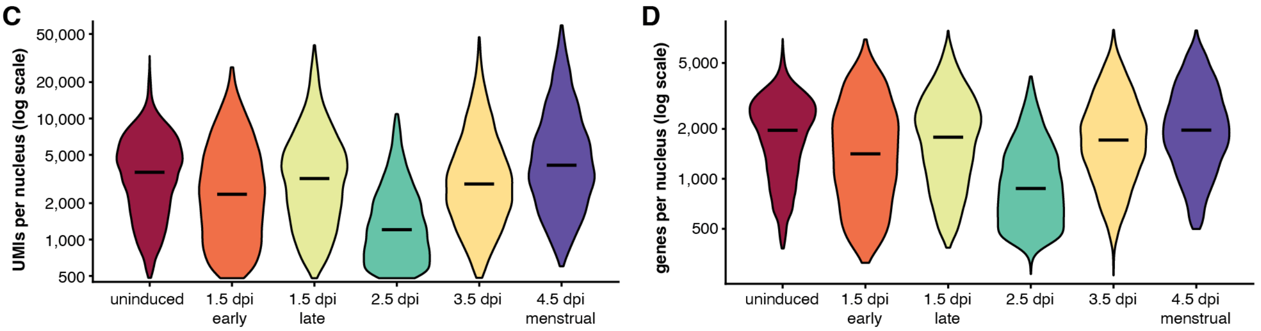


**Figure S12. Quality control and cell type annotation for Slide-tags datasets from GsD X-Mens samples.**
**A.** Table summarizing the timepoint, section thickness, number of 10X lanes, number of cells, and percentage of cells spatially mapped for each Slide-tags sample. **B.** Dot plot of marker gene expression across cell types. Dot size indicates the fraction of cells in which the gene was detected, and color indicates mean log normalized expression scaled from 0 to 1 across cell types for each gene. **C-D**. Violin plots of UMIs (C) and genes (D) detected per nucleus at each time point; y-axes are log-scaled and black lines mark medians. dpi: days post-induction; mens: menstruation; epi: epithelial cells; eFib: endometrial fibroblasts; dec: decidual cells; vsc: vasculature cells; imm: immune cells; smc: smooth muscle cells; pm: perimetrium cells.

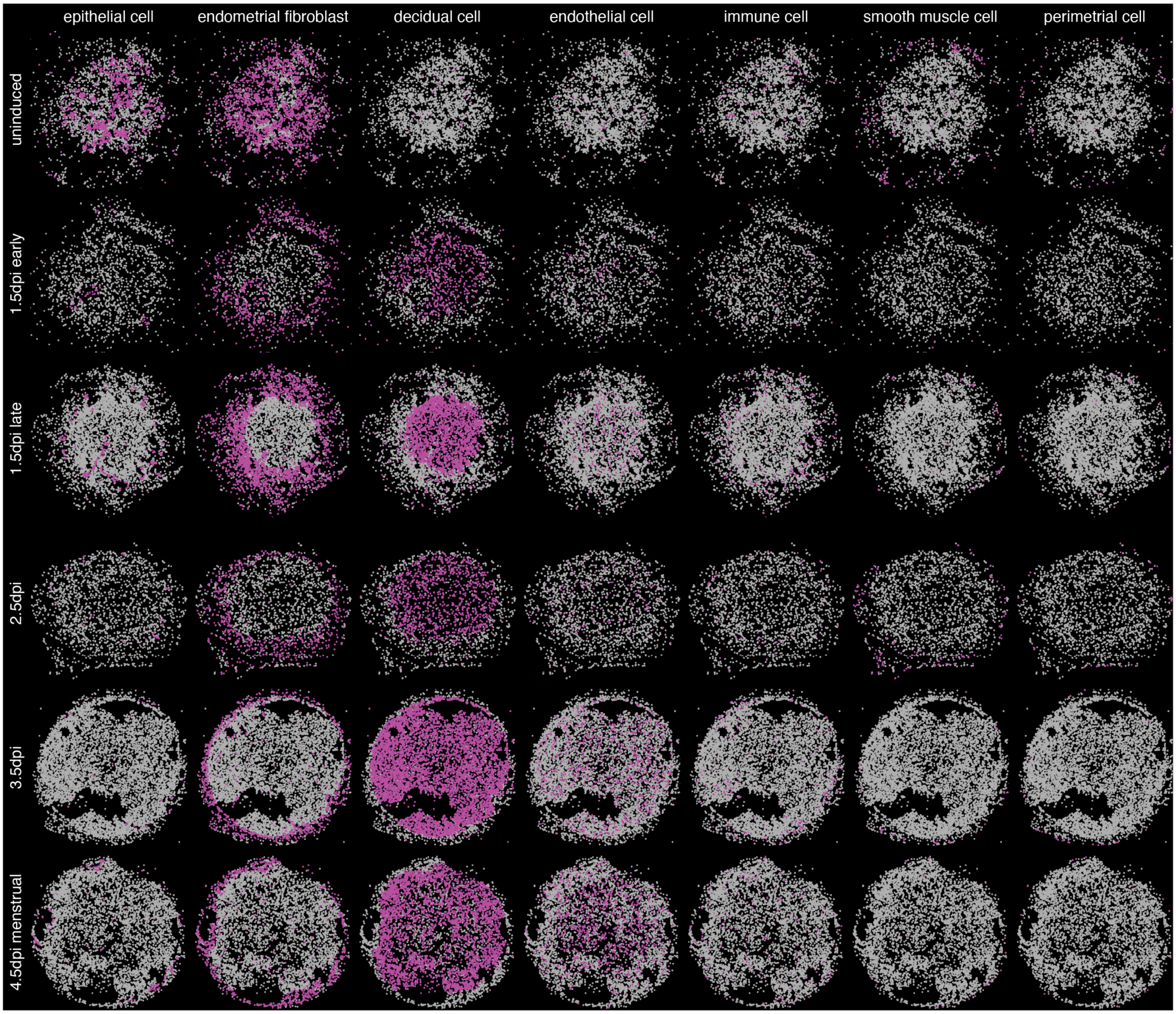


**Figure S13. Spatial mapping of the cell types in GsD X-Mens Slide-tags samples.**
Spatial distribution of major cell types (columns) in uterine sections from GsD X-Mens mice sampled at six stages (rows). Each panel highlights one cell type (magenta) overlaid on all other cells (gray). dpi: days post-induction.

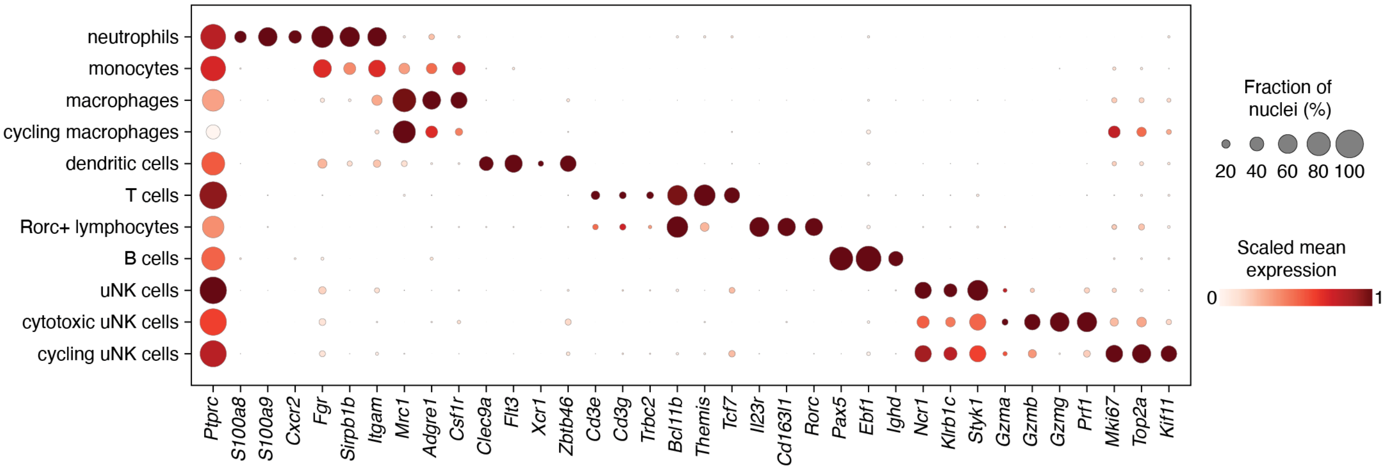


**Figure S14. Immune cell types in the X-Mens dataset.**
Dot plot of marker gene expression across immune cell subtypes. Dot size indicates the fraction of cells in which the gene was detected, and color indicates mean log normalized expression scaled from 0 to 1 across subtypes for each gene.

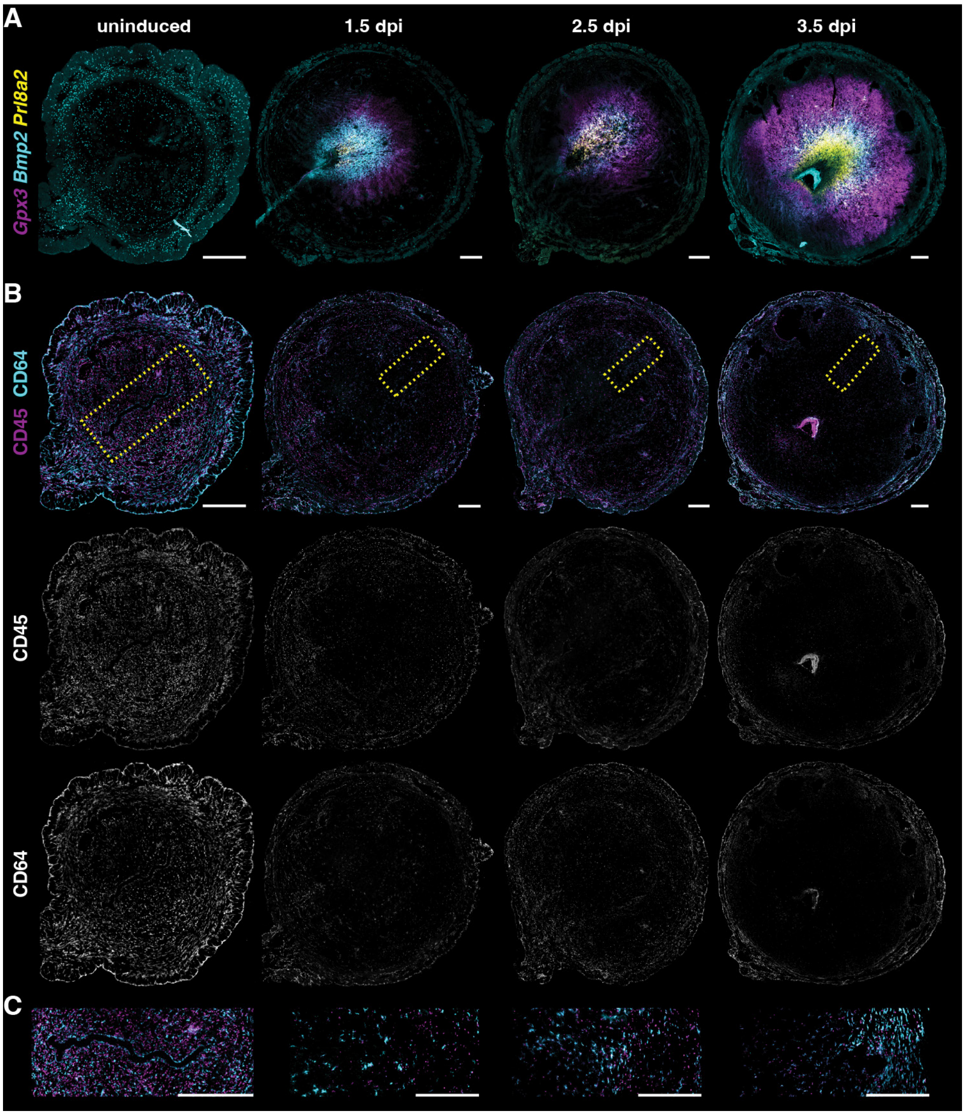


**Figure S15. Immune cell distribution in the endometrium of GsD X-Mens mice.**
**A.** Fluorescence in situ hybridization images of uterine cross sections from GsD X-Mens mice at the indicated stages. *Gpx3* (magenta), *Bmp2* (cyan) and *Prl8a2* (yellow), mark decidual cell populations. These images are provided as spatial references for decidual compartments relative to immune cells, which were stained in the adjacent serial sections and are shown in panel B. The GsD 3.5 dpi image is also shown in Figure 4B, and all images are reproduced in Figure S19. **B.** Immunofluorescence images of the uterine cross sections immediately adjacent to those shown in (A), stained for leukocytes (CD45, magenta) and macrophages (CD64, cyan). Composite images are shown in the top row, with the CD45 (middle) and CD64 (bottom) channels shown separately in grayscale. **C.** Insets showing the regions marked with yellow boxes on panel B. The images are oriented to position the mesometrial side on the left. Scale bars: 500 $\mu$m. dpi: days post-induction. Imaging settings, brightness, and contrast were adjusted independently to enable visualization of key structures.

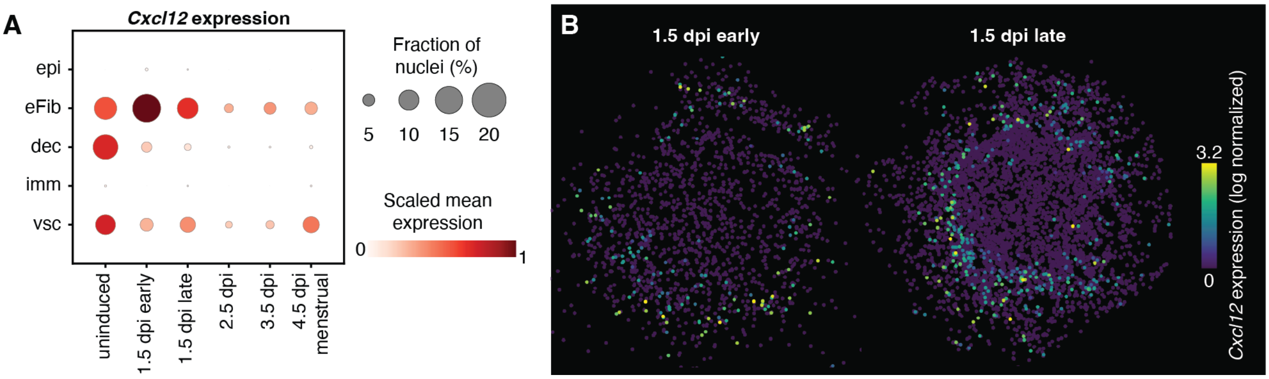


**Figure S16. *Cxcl12* expression during early decidualization in GsD X-Mens mice.**
**A.** Dot plot of *Cxcl12* expression across endometrial cell types. Dot size indicates the fraction of cells in which the gene was detected, and color indicates mean log normalized expression scaled from 0 to 1 across cell types. epi: epithelial cells; eFib: endometrial fibroblasts; dec: decidual cells; vsc: vasculature cells; imm: immune cells; smc: smooth muscle cells; pm: perimetrial cells. **B.** Spatial maps of *Cxcl12* expression at each timepoint. Color indicates log-normalized expression, ln(counts per 10,000 + 1), on a scale shared across all timepoints. The color scale ranges from 0 to 3.2, the 99.5th percentile among *Cxcl12*-expressing nuclei; nuclei above this value are shown at the maximum color. dpi: days post-induction.

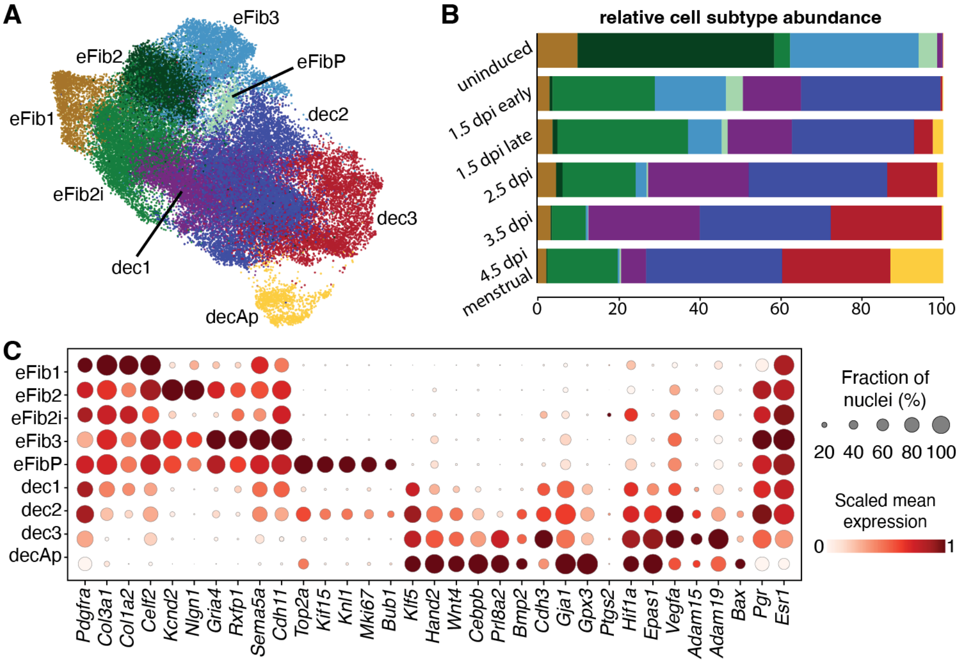


**Figure S17. Endometrial fibroblast and decidual subclusters across the X-Mens cycle.**
**A.** UMAP embedding of endometrial fibroblast and decidual cells, colored by subcluster. eFib1–3: endometrial fibroblast subclusters 1–3; eFib2i: induced endometrial fibroblasts 2; eFibP: proliferating endometrial fibroblasts; dec1–3: decidual cell subclusters 1–3; decAp: apoptotic decidual cells. **B.** Relative abundance of each subtype at the indicated stages, expressed as a percentage of all endometrial fibroblast and decidual cells recovered at that stage. Colored as in panel A. **C.** Dot plot of marker gene expression across subclusters. Dot size indicates the fraction of cells in which the gene was detected, and color indicates mean log normalized expression scaled from 0 to 1 across subclusters for each gene. dpi: days post-induction.

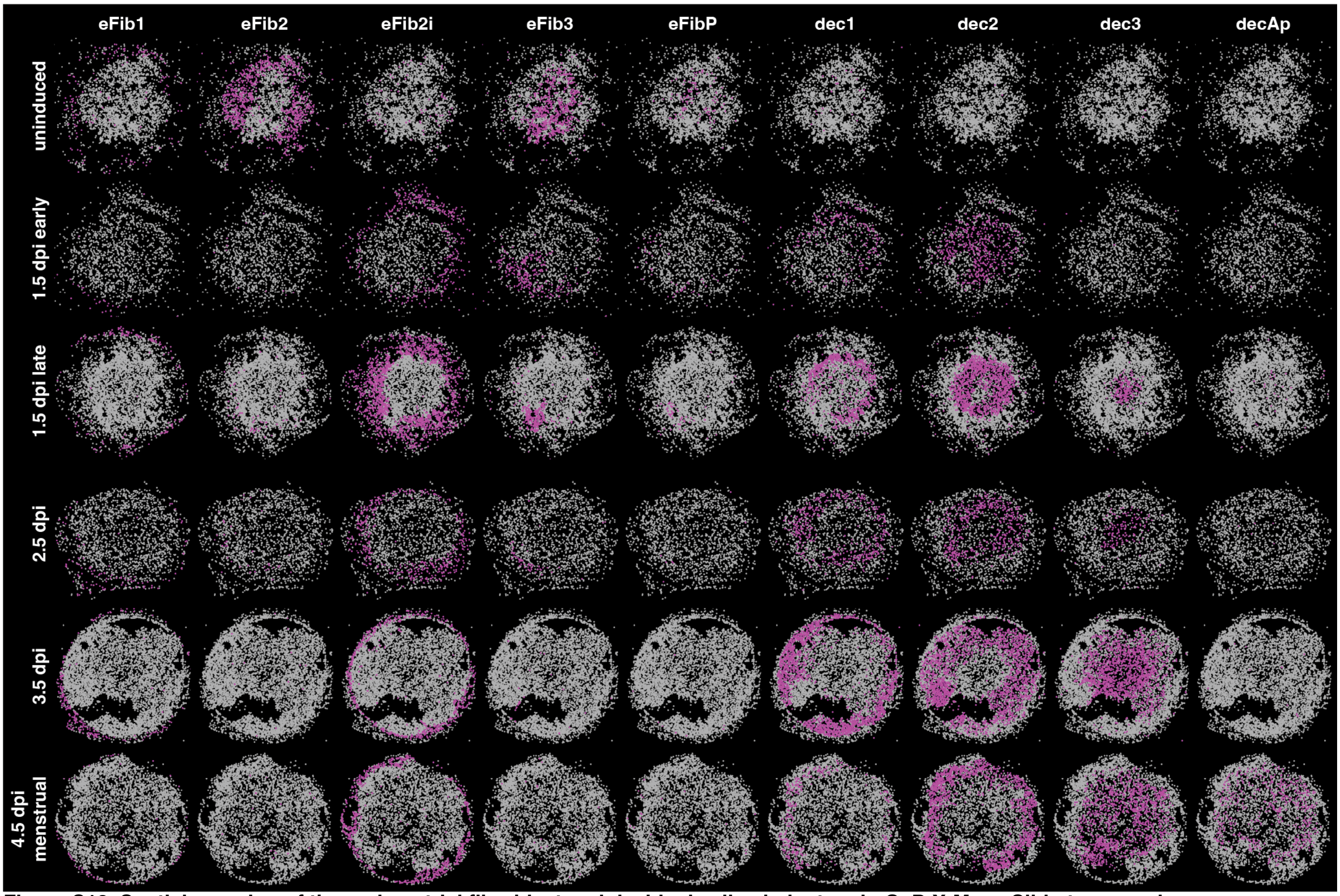


**Figure S18. Spatial mapping of the endometrial fibroblast and decidual cell subclusters in GsD X-Mens Slide-tag samples.**
Spatial distribution of endometrial fibroblast (eFib) and decidual (dec) cell subclusters (columns) in uterine sections from GsD X-Mens mice sampled at six stages (rows). Each panel highlights one cell subcluster (magenta) overlaid on all other cells (gray). dpi: days post-induction; eFib1–3: endometrial fibroblast subclusters 1–3; eFib2i: endometrial fibroblasts 2 induced; eFibP: proliferating endometrial fibroblasts; dec1–3: decidual cell subclusters 1–3; decAp: apoptotic decidual cells.

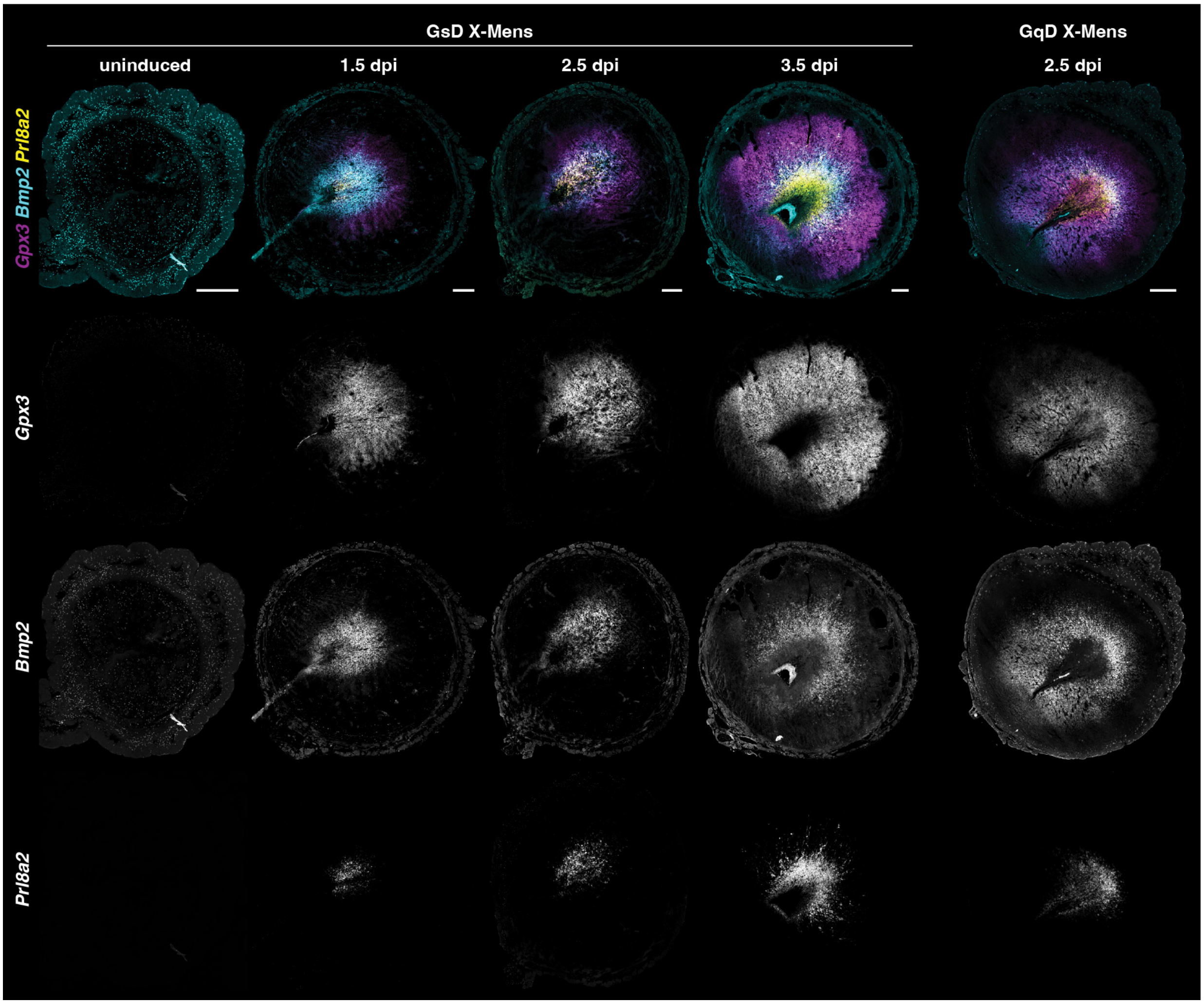


**Figure S19. Radial organization of decidual markers across decidualization in X-Mens mice.**
Fluorescence in situ hybridization images of uterine cross sections from GsD X-Mens mice at the indicated stages (left four columns) and from a GqD X-Mens mouse at 2.5 dpi (right column), showing *Gpx3* (magenta), *Bmp2* (cyan) and *Prl8a2* (yellow). The same GsD 3.5 dpi micrograph is also shown in Figure 4B. The composite images are the same as those shown in Figure S15A. Composite images are shown in the top row, with the individual channels shown below in grayscale. Scale bars: 500 $\mu$m. dpi: days post-induction. Imaging settings, brightness, and contrast were adjusted independently to enable visualization of key structures.

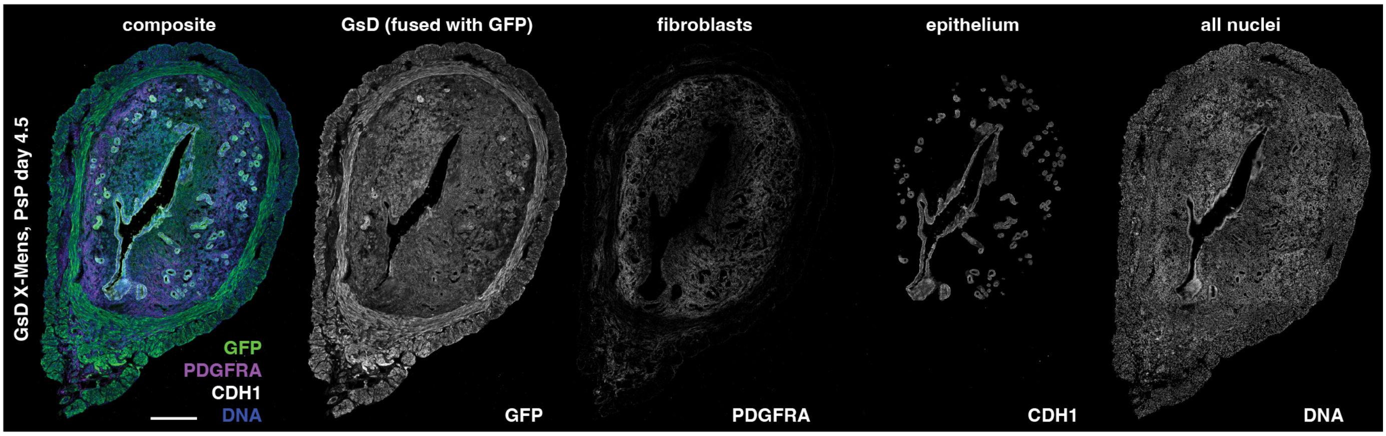


**Figure S20. Expression pattern of GsD in the uterus at pseudopregnancy (PsP) day 4.5.**
Representative cross-section of a GsD X-Mens mouse uterus stained for GFP (green; GsD-GFP fused protein), fibroblasts (PDGFRA, magenta), epithelium (CDH1, white). DNA is counterstained with DAPI (blue). The composite image is shown on the left, with individual channels shown in grayscale. Scale bar: 400 $\mu$m.

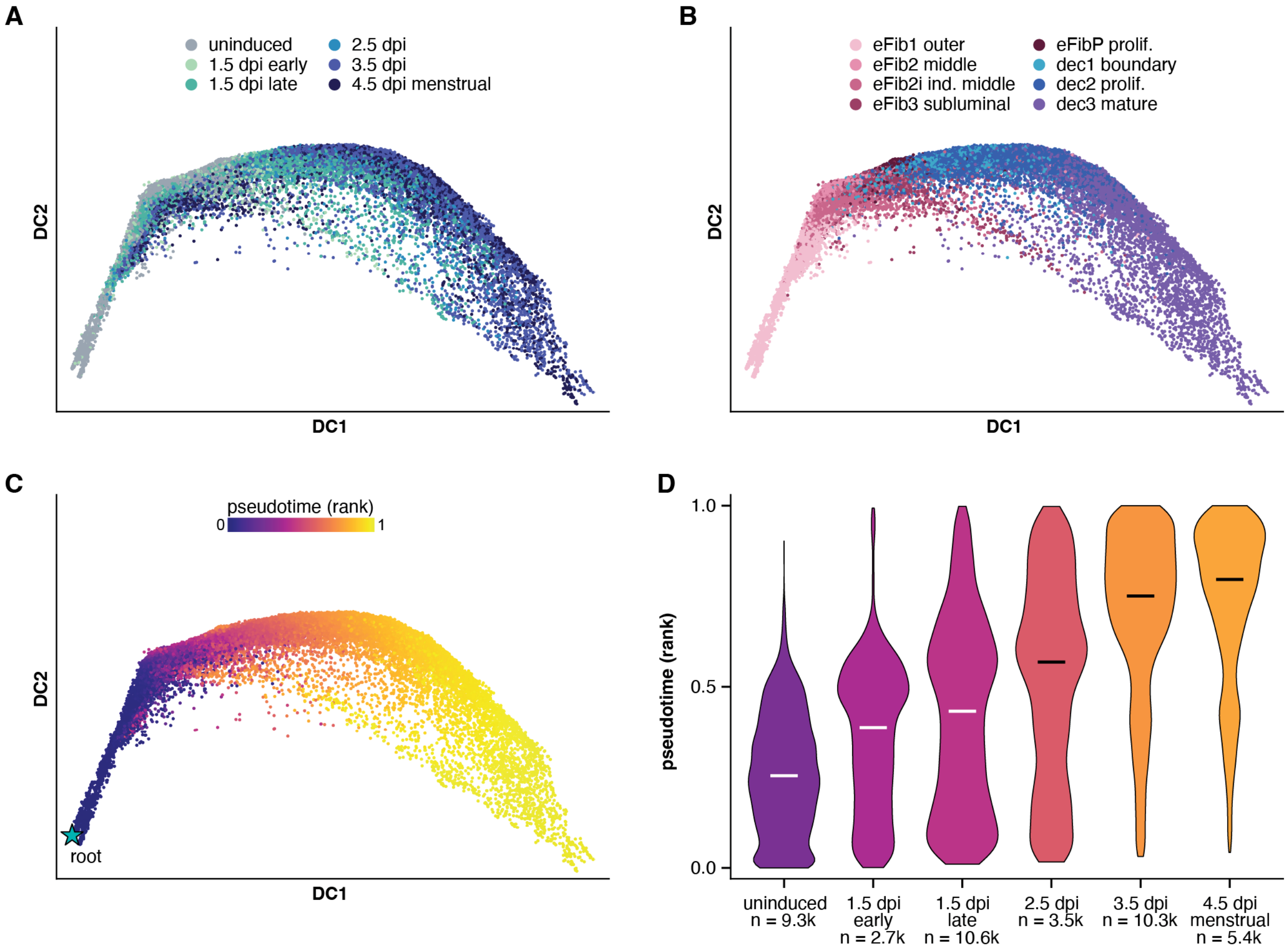


**Figure S21. Pseudotime analysis of GsD X-Mens endometrial fibroblast and decidual cells.**
**A-C.** Diffusion map (diffusion components 1 and 2) of endometrial fibroblast and decidual cells from all GsD X-Mens samples, excluding apoptotic decidual cells (decAp), colored by time point (A), colored by cell subcluster (B), colored by diffusion pseudotime value, rank-normalized to the range 0 to 1, with the root cell indicated by a star (C). The dataset comprises six animals, one per timepoint indicated in panel A. **D.** Distribution of pseudotime values for all cells in each sample. Horizontal bars indicate the median and fill color scales with the median value. n indicates the number of cells per sample. dpi: days post-induction; DC: diffusion component; eFib1–3: endometrial fibroblast subclusters 1–3; eFib2i: endometrial fibroblasts 2 induced; eFibP: proliferating endometrial fibroblasts; dec1–3: decidual cell subclusters 1–3.

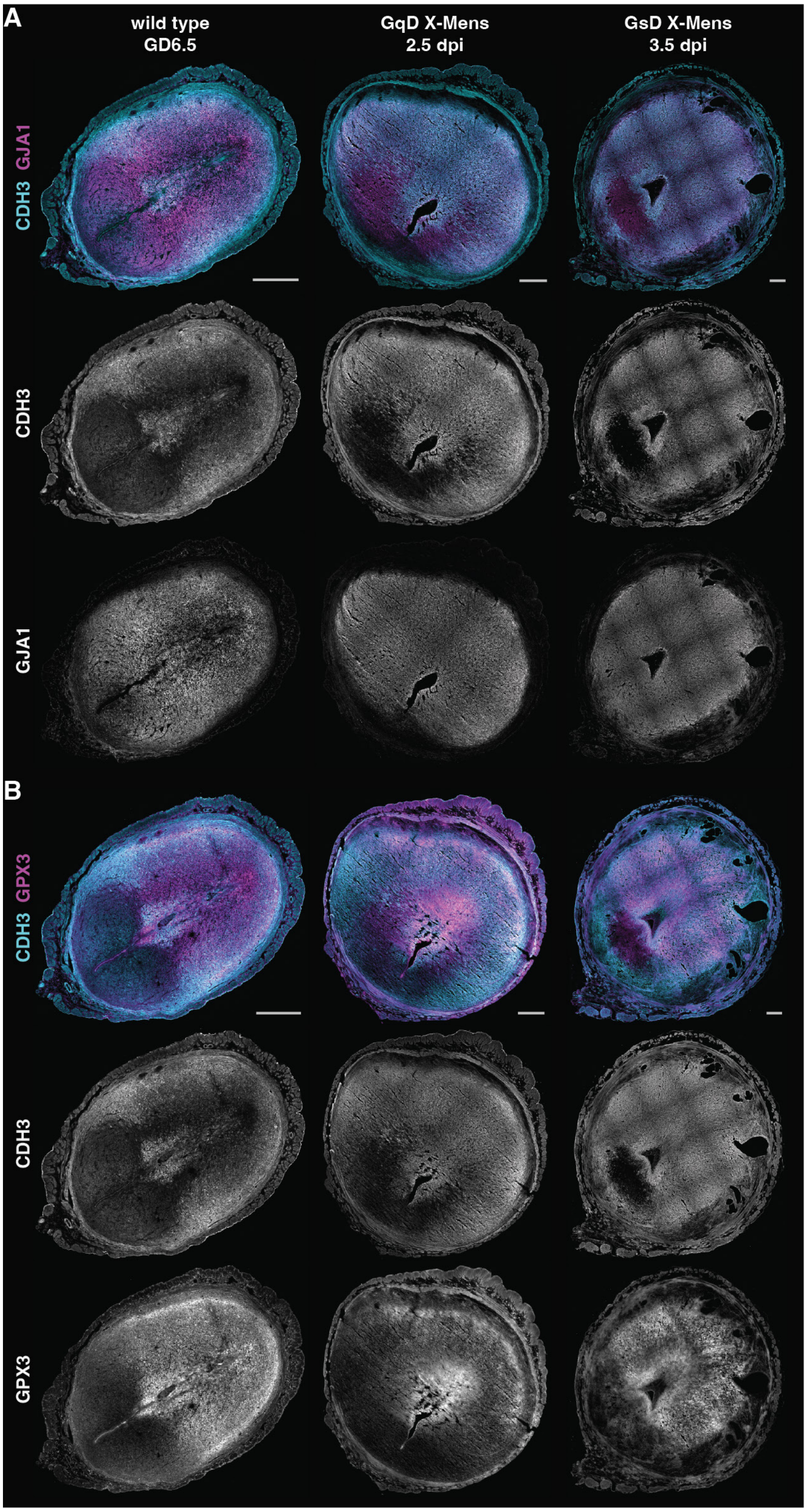


**Figure S22. Pan-decidual markers in X-Mens and pregnant uteri.**
**A.** Immunofluorescence images of uterine cross sections from a wild-type mouse at gestational day 6.5 (GD6.5, left), a GqD X-Mens mouse at 2.5 dpi (middle) and a GsD X-Mens mouse at 3.5 dpi (right), stained for CDH3 (cyan) and GJA1 (magenta). Composite images are shown in the top row, with the CDH3 (middle) and GJA1 (bottom) channels shown separately in grayscale. **B.** Uterine cross sections from the same animals as in panel A, stained for CDH3 (cyan) and GPX3 (magenta), shown as in panel A. Scale bars: 500 $\mu$m. dpi: days post-induction. Imaging settings, brightness, and contrast were adjusted independently to enable visualization of key structures.

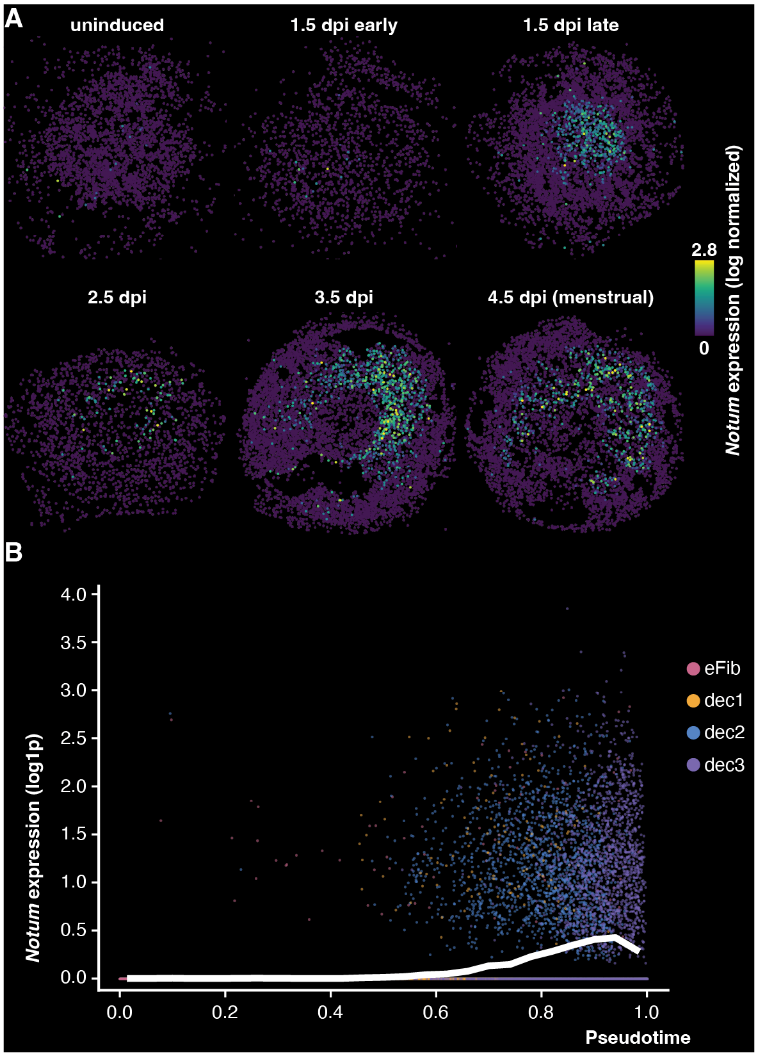


**Figure S23. Notum expression across X-Mens decidualization.**
**A.** Spatial mapping of Notum expression in GsD X-Mens Slide-tags samples at the indicated stages. dpi: days post-induction. **B.** Notum expression normalized as ln(counts per 10,000 + 1), against rank-normalized pseudotime in GsD X-Mens fibroblast and decidual cells, colored by cell subcluster (eFib: all fibroblast subclusters; dec1–3: decidual cell subclusters 1–3). The white line is the mean expression in 25 equal-width pseudotime bins. The bootstrap 95% confidence interval of each bin mean is drawn, but is narrower than the line. Apoptotic decidual cells, which have no pseudotime value, are not shown.

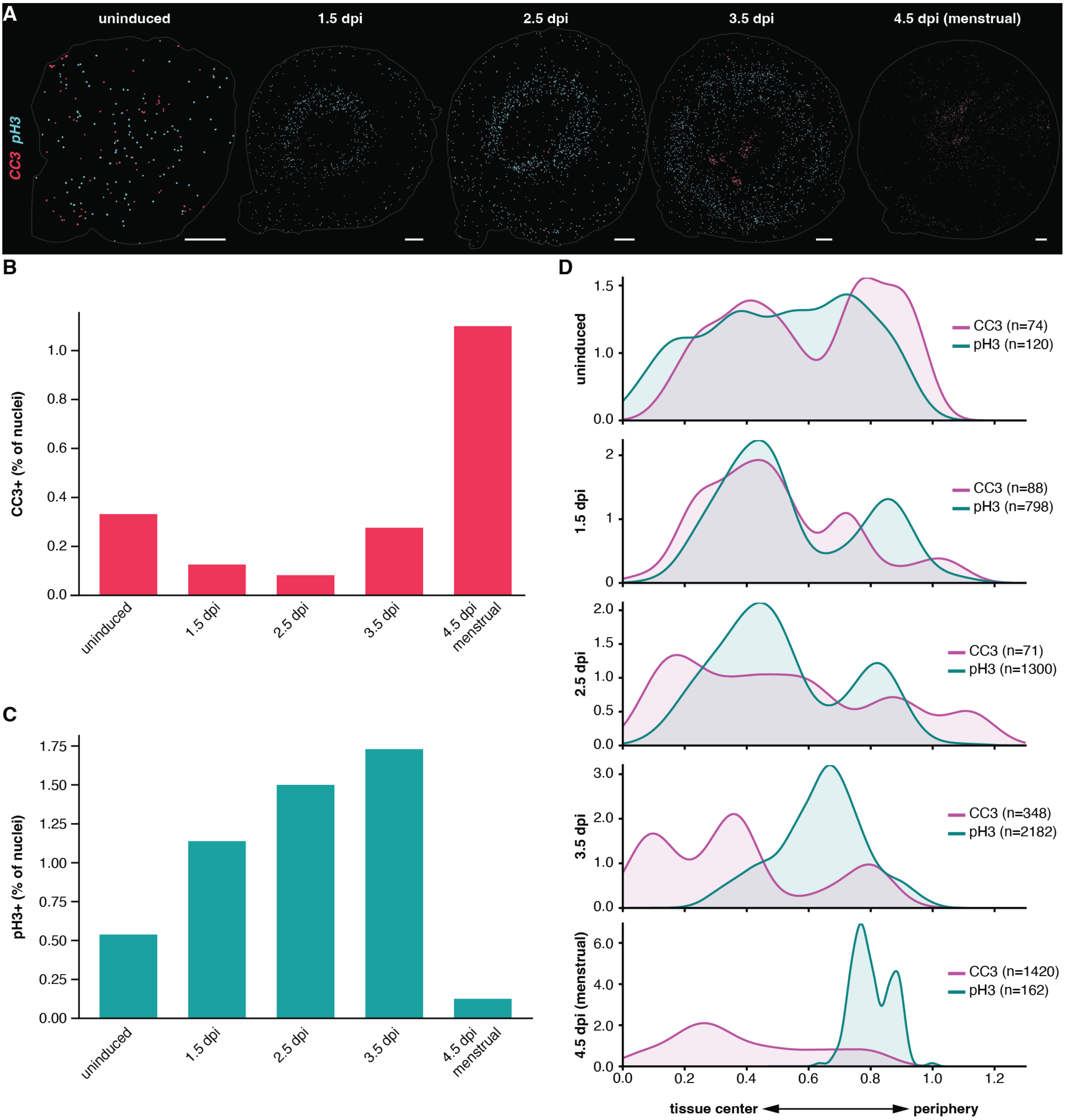


**Figure S24. Quantification of apoptosis and proliferation across X-Mens decidualization.**
Sections are adjacent to those shown in Figure S19, so the position of CC3- and pH3-positive cells can be compared directly with the decidual zones defined by in situ hybridization. **A.** Spatial distribution of cleaved caspase 3 (CC3, magenta) and phospho-histone H3 (pH3, cyan) in GsD X-Mens uterine cross sections at the indicated stages. Immunofluorescence images were segmented to identify CC3- and pH3-positive objects, which are displayed as dots enlarged for visibility. The gray outline indicates the tissue boundary. The 3.5 dpi image is reproduced from Figure 4C. Scale bars: 500 $\mu$m. **B.** Quantification of CC3-positive and **C.** pH3-positive objects at each stage, expressed as a percentage of total segmented nuclei in the cross-section. D. Distribution of CC3 (magenta) and pH3 (cyan) objects along the normalized radial axis of the uterine cross sections at each stage (normalized distance from section centroid; r/r_equiv), visualized using kernel density estimation (KDE). n indicates the number of objects. dpi: days post-induction.

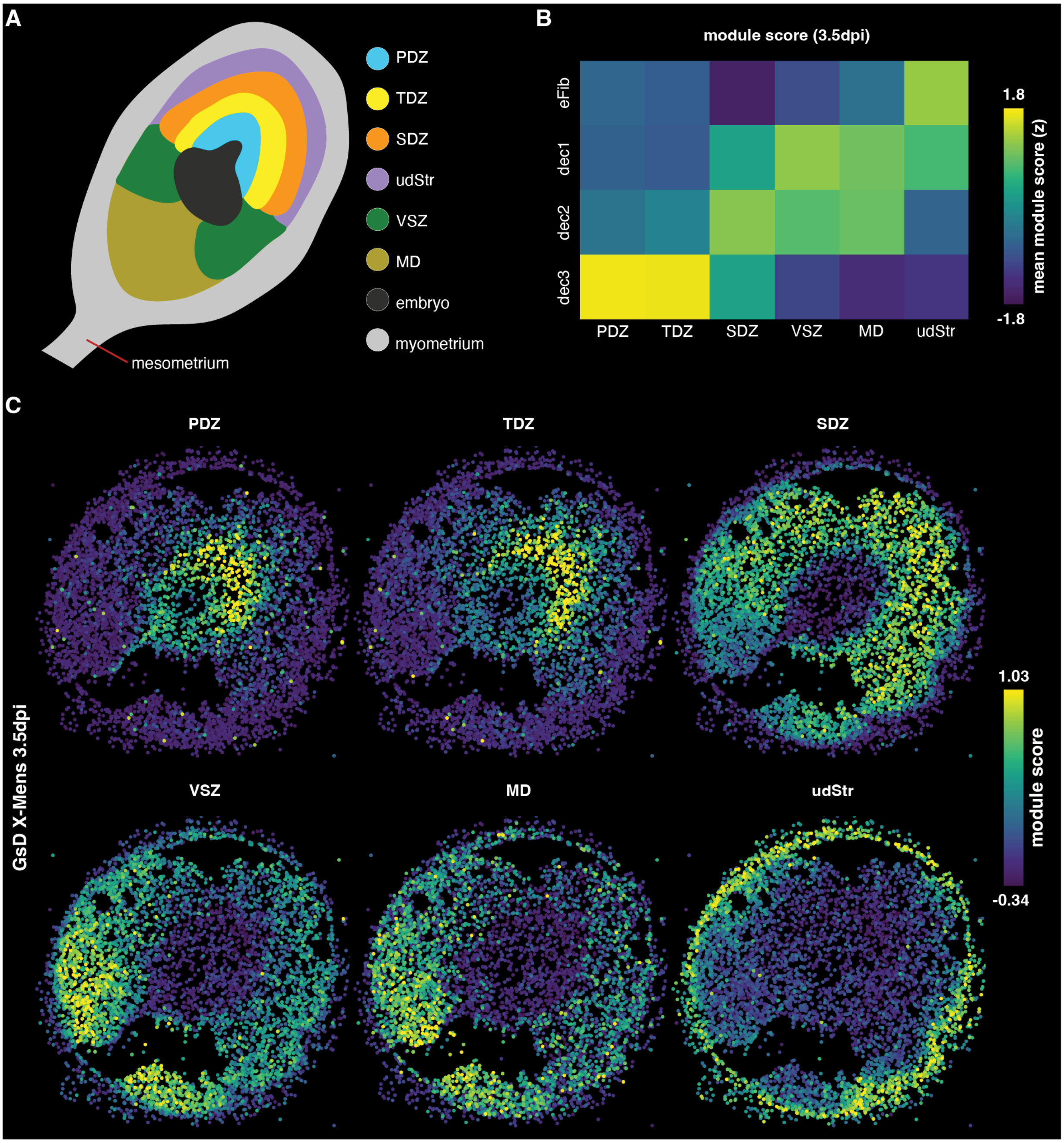


**Figure S25. Comparison of X-Mens decidual populations with uterine microenvironments of mouse pregnancy.**
Gene signatures for each uterine zone defined by spatial transcriptomics of the mouse uterus at gestational day 7.5 were scored in GsD X-Mens endometrial fibroblasts and decidual cells at 3.5 dpi. **A.** Schematic showing the decidual zones in the mouse uterus at gestational day (GD) 7.5, identified in (Li et al., 2022); the embryo and myometrium are also indicated. The uterine ligament (mesometrium), along which blood vessels enter the uterus, is shown at the bottom left. As the schematic demonstrates, the mesometrial-anti-mesometrial axis shapes the organization of decidual zones. **B.** Heatmap showing the module score for each gestational uterine zone signature (columns) in X-Mens endometrial fibroblasts and decidual cell subclusters (rows). Scores are averaged within each subcluster and z-scored across subtypes for each signature, so that color indicates which subcluster scores highest for a signature rather than the absolute score. The five endometrial fibroblast subclusters are pooled as eFib, and apoptotic decidual cells (decAp) were not scored. **C.** Spatial mapping of the module score for each gestational zone signature in the 3.5 dpi GsD X-Mens uterine section. The color range is shared across the six signatures and clipped to the 1st and 99th percentiles of the scored nuclei; nuclei without a score are shown in dark gray. PDZ: primary decidual zone; TDZ: transition decidual zone; SDZ: secondary decidual zone; VSZ: vascular sinus zone; MD: mesometrial decidua; udStr: undifferentiated stroma; eFib: endometrial fibroblasts; dec1–3: decidual cell subclusters 1–3; dpi: days post-induction.

A

| timepoint | section thickness | #10X lanes | number of cells | spatially mapped (%) |
|---|---|---|---|---|
| secretory | 20$\mu$m | 3 | 30,573 | 77.23 |
| menstrual | 20$\mu$m | 3 | 35,136 | 68.72 |

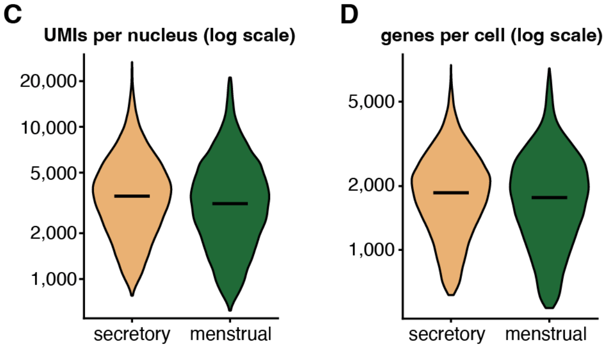


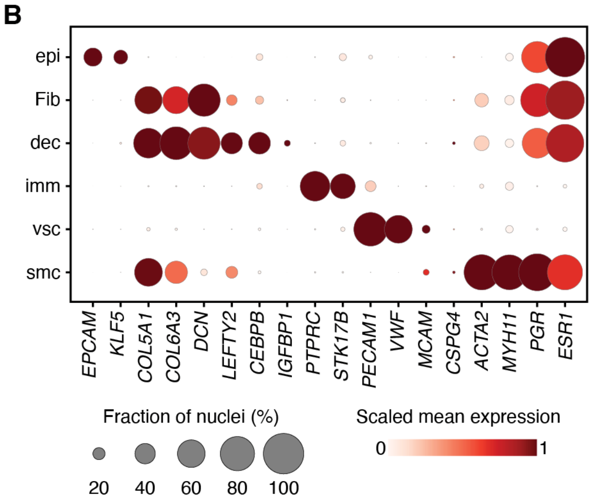


**Figure S26. Quality control and cell type annotation for Slide-tags datasets from human samples.**
**A.** Table summarizing the cycle phase, section thickness, number of 10X lanes, number of cells, and percentage of cells spatially mapped for each Slide-tags sample. **B.** Dot plot of marker gene expression across cell types. Dot size indicates the fraction of cells in which the gene was detected, and color indicates mean log normalized expression scaled from 0 to 1 across cell types for each gene. **C.** Violin plots showing the distribution of log-transformed counts per cell and **D.** genes per cell for each sample. epi: epithelial cells; Fib: fibroblasts; dec: decidual cells; imm: immune cells; vsc: vasculature cells; smc: smooth muscle cells.

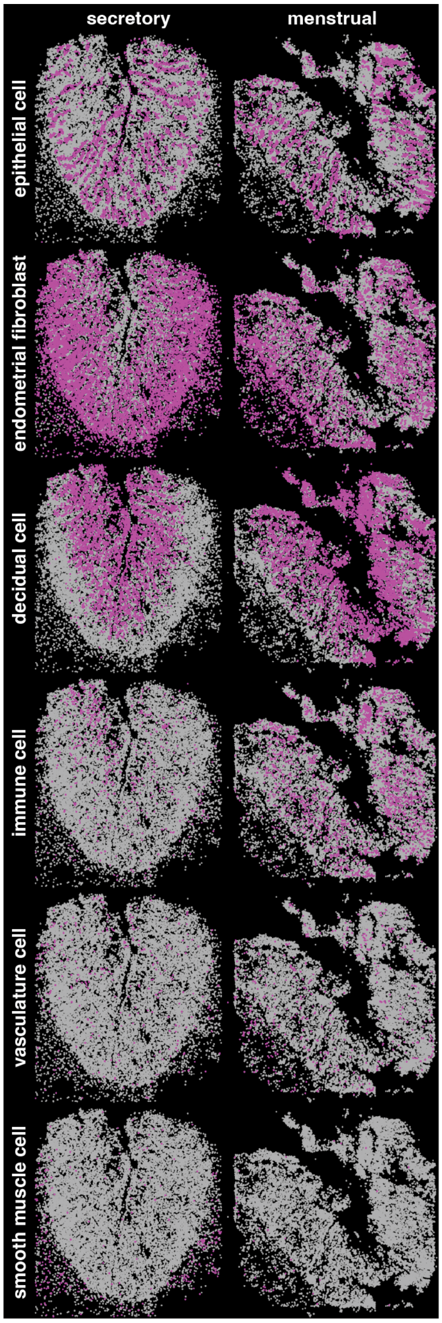


**Figure S27. Spatial mapping of the cell types in human Slide-tags samples.** Spatial distribution of major cell types (rows) in endometrial sections from the secretory and menstrual samples (columns). Each panel highlights one cell type (magenta) overlaid on all other cells (gray).

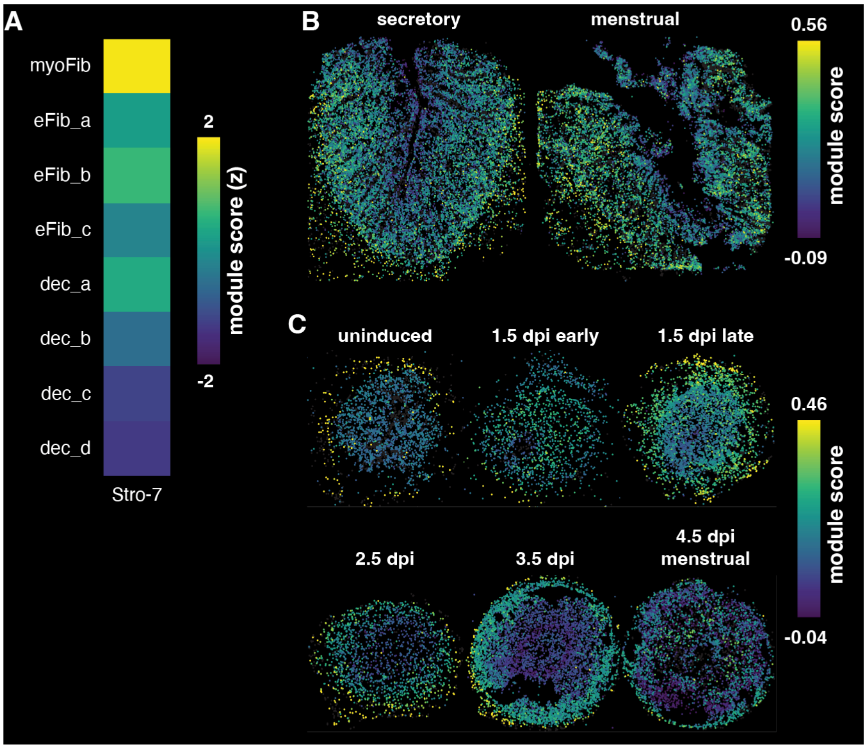


**Figure S28. Expression of the C7+ stromal (Stro-7) signature in human and X-Mens uteri.**
Each nucleus was scored for the 50 markers with the largest average log2 fold change of the Stro-7 stromal cluster of the human uterine atlas of (Ulrich et al., 2024). **A.** Heatmap showing the module score for the Stro-7 signature in each human endometrial fibroblast and decidual subcluster. Scores are averaged within each subcluster and z-scored across the eight subclusters, so that color indicates which subtype scores highest for the signature rather than the absolute score; both donors are pooled. **B.** Spatial mapping of the Stro-7 module score in the secretory and menstrual human Slide-tags datasets. The color range is shared between the two sections and clipped to the 1st and 99th percentiles of the scored nuclei; nuclei without a score are shown in dark gray. **C.** Spatial mapping of the Stro-7 module score in the GsD X-Mens Slide-tags datasets at the indicated stages, transferred from the human nuclei through the SAMap embedding (see Materials and Methods). myoFib: myometrial fibroblasts; eFib_a–c: endometrial fibroblast subclusters a–c; dec_a–d: decidual cell subclusters a–d; dpi: days post-induction.

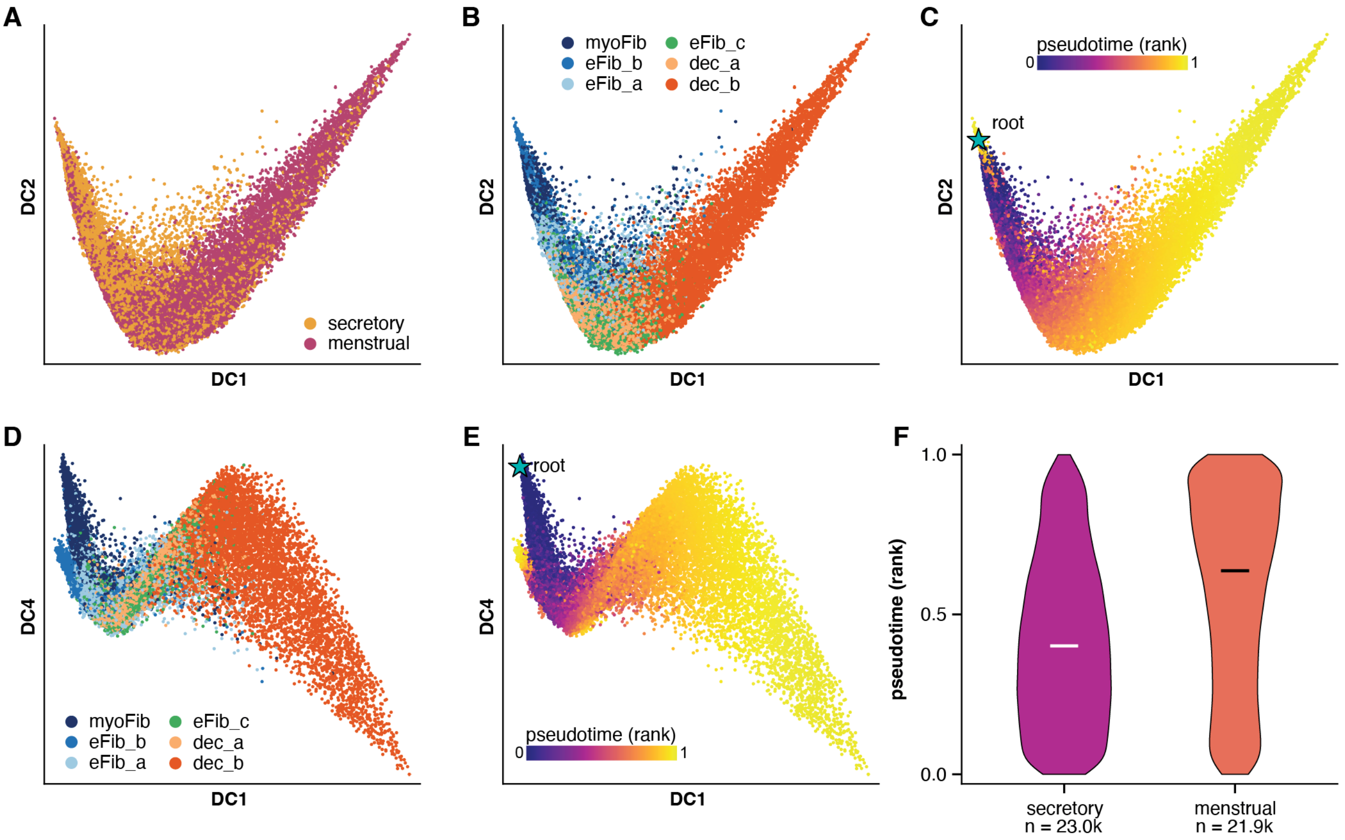


**Figure S29. Pseudotime analysis of human endometrial fibroblast and decidual cells.**
Diffusion map of endometrial fibroblast and decidual cells from both human Slide-tags samples, excluding apoptotic decidual cells (dec_c and dec_d). A-C. Diffusion components 1 and 2, colored by **A.** cycle phase, **B.** cell subcluster and **C.** diffusion pseudotime, rank-normalized to the range 0 to 1, with the root cell indicated by a star. D-E. Diffusion components 1 and 4, which separate myoFib from eFib-a cells, colored by **D.** cell subcluster and **E.** pseudotime. **F.** Distribution of ranked pseudotime values for all cells in each sample; horizontal bars indicate the median and fill color scales with the median value. n indicates the number of cells per sample. DC: diffusion component; myoFib: myometrial fibroblasts; eFib_a–c: endometrial fibroblast subclusters a–c; dec_a–b: decidual cell subclusters a and b.

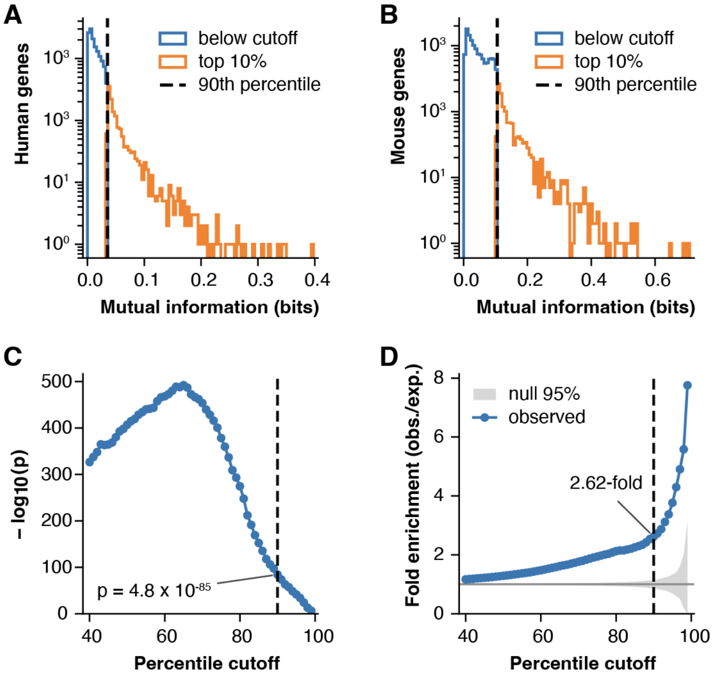


**Figure S30. Selection of pseudotime-associated genes based on mutual information.**
Distribution of estimated mutual information values, using the k-nearest-neighbor estimator of (Kraskov et al., 2004) as implemented in scikit-learn (Pedregosa et al., 2011), for **A.** human and **B.** mouse genes significant at a false discovery rate below 0.05. Genes above the 90th percentile (dashed line; genes highlighted in orange) were taken as pseudotime-associated. **C.** Significance of the overlap between the human and mouse top 10% pseudotime-associated gene sets (hypergeometric test against a background of 9,800 one-to-one orthologs), and **D.** fold enrichment of the observed overlap over that expected by chance, as a function of the percentile cutoff used to define the gene sets. The gray band in panel D indicates the 95% interval of the hypergeometric null distribution. Dashed lines indicate the 90th percentile cutoff used in this study, at which 379 genes are shared between the 1,231 human and 1,153 mouse genes (2.62-fold; $p = 4.8 \times 10^{-85}$).

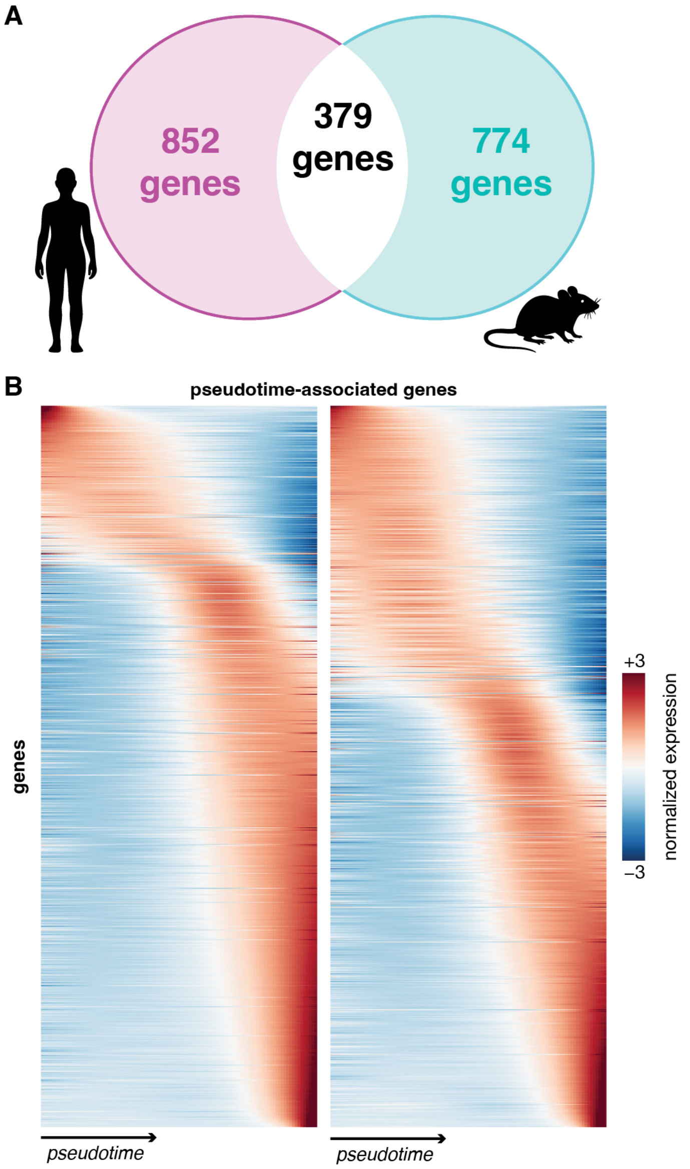


**Figure S31. Pseudotime-associated gene programs in human and X-Mens endometrium.**
**A.** Overlap between the top 10% of genes most strongly associated with pseudotime in human and X-Mens mice. Analysis was restricted to the 9,800 one-to-one orthologs that were detected in at least 1% of fibroblasts and decidual cells. The number of top 10% of the genes that are significantly associated with pseudotime are shown in magenta and cyan, for human and X-Mens, respectively. The number of top pseudotime-associated genes that are shared between species are shown in white. **B.** Normalized expression of the top pseudotime-associated genes along pseudotime in human (left, n = 1,231 genes) and X-Mens mice (right, n = 1,153 genes). Rows are each species' own gene set, ordered independently by the pseudotime centroid of their expression. Color indicates per-gene z-scores, clipped at ±3.

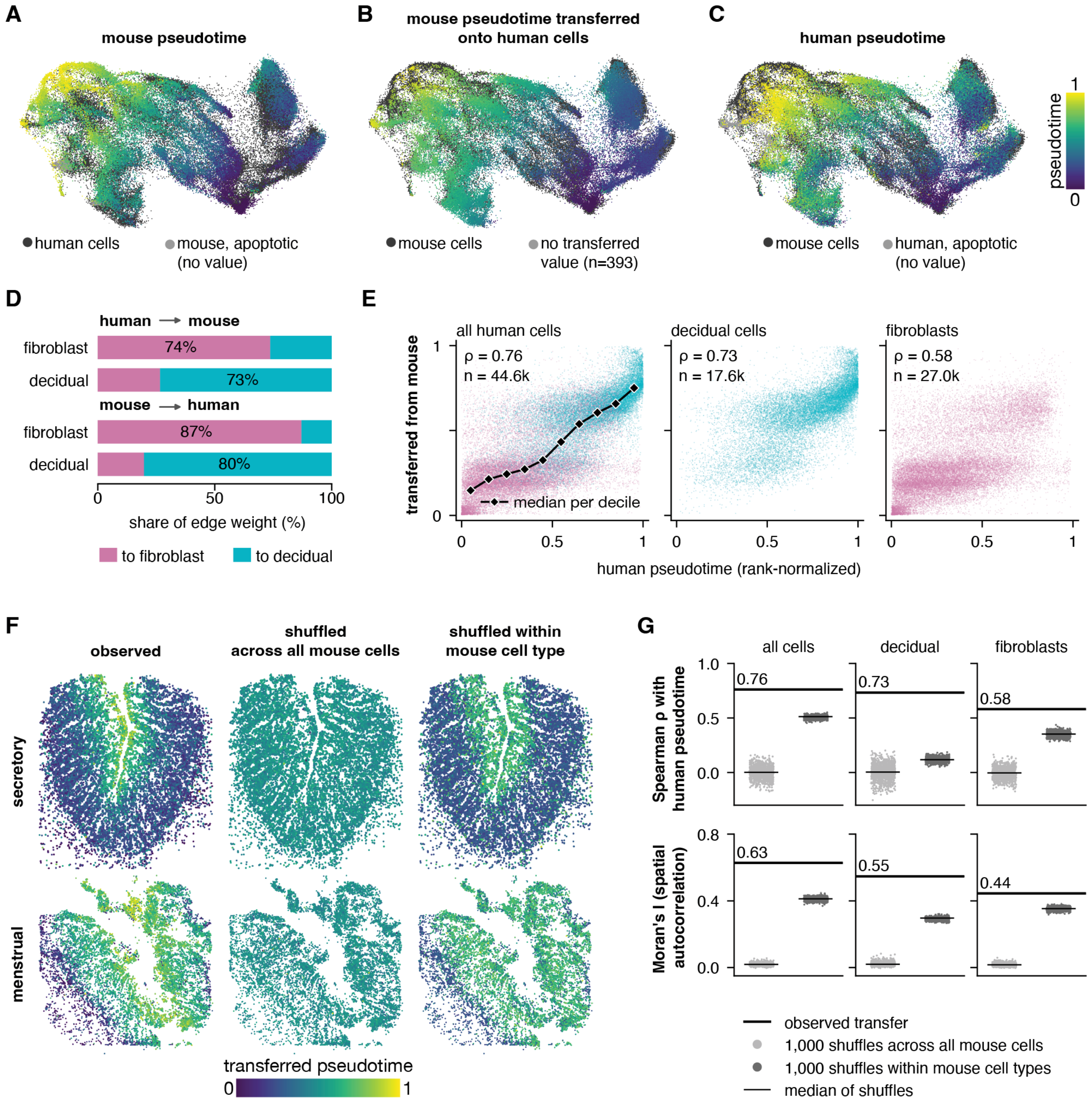


**Figure S32. Validation of cross-species integration and pseudotime transfer by SAMap.**
**A-C.** SAMap joint embedding of human and X-Mens endometrial fibroblasts and decidual cells. In each panel, cells of the other species are shown in dark gray, and cells without a pseudotime value are shown in light gray. **A.** Mouse cells colored by mouse pseudotime. **B.** Human cells colored by pseudotime transferred from their mouse neighbors in the SAMap graph. These data are reproduced from Figure 6C for comparison. **C.** Human cells colored by human pseudotime, computed independently of the X-Mens data. **D.** Distribution of cross-species SAMap edge weights from human to mouse cell populations (top) and from mouse to human cell populations (bottom). E. Comparison of mouse pseudotime transferred to human cells and pseudotime independently calculated from the human data, shown for all cells (left), decidual cells (middle) and fibroblasts (right), colored as in panel D. The black line indicates the median transferred pseudotime per decile of human pseudotime; n indicates the number of cells per panel, ρ: Spearman correlation. **F.** Spatial maps of transferred pseudotime on the secretory (top) and menstrual (bottom) human sections, showing observed transfer (left), transfer after shuffling mouse pseudotime across all mouse cells (middle), and transfer after shuffling within fibroblast and decidual compartments (right). The observed transfer shown at left is also shown in Figure 6E. **G.** Spearman correlation between transferred and human pseudotime (top) and spatial autocorrelation of transferred pseudotime (Moran's I, computed from the 10 nearest spatial neighbors; bottom), for all cells and separately within decidual cells and within fibroblasts.

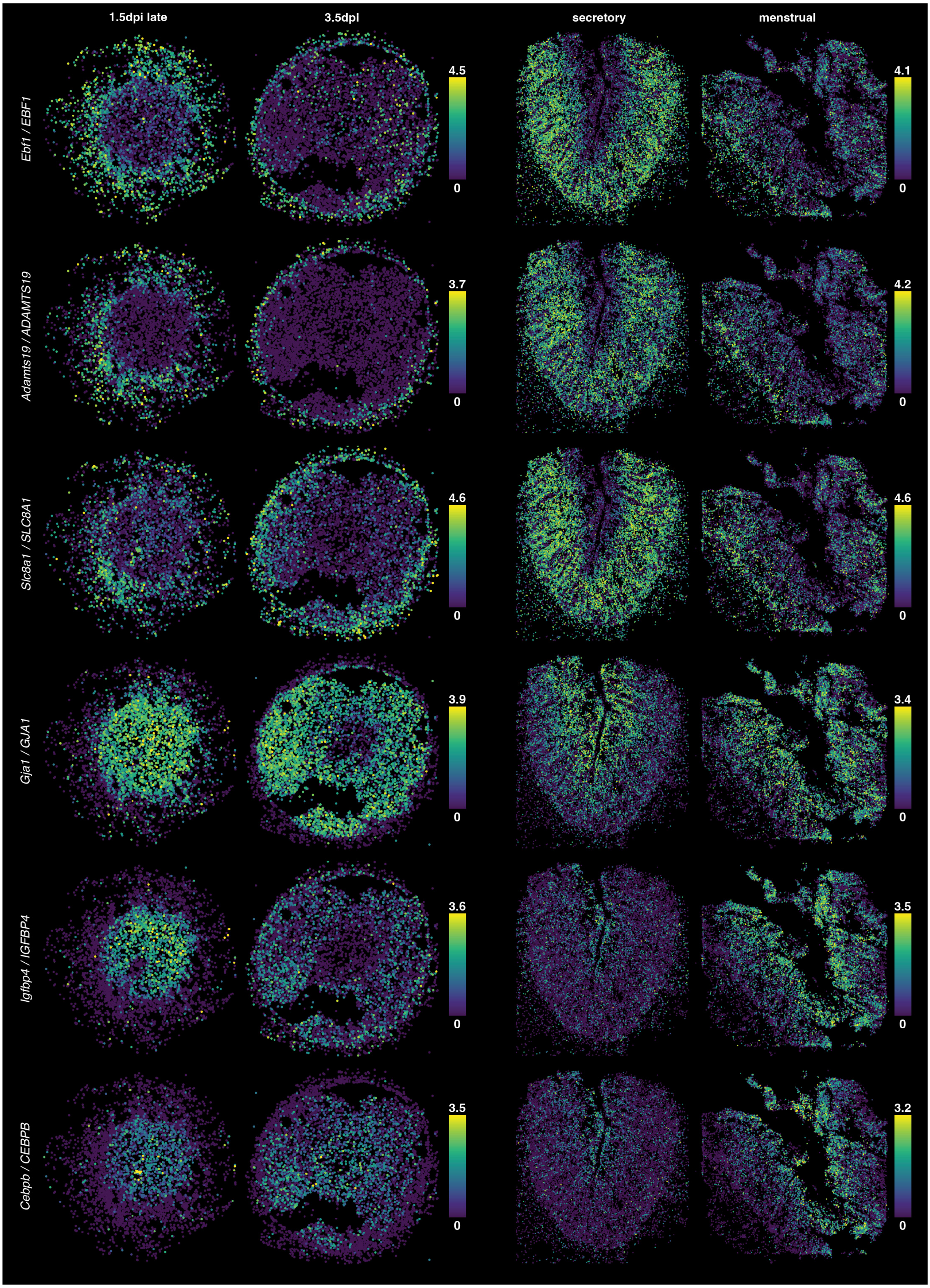


**Figure S33. Functionalis and basalis markers in human and X-Mens endometrium.**
Spatial maps of the expression of genes associated with the endometrial basalis (top three rows) or functionalis (bottom three rows), shown as orthologous pairs in GsD X-Mens uterine cross sections at 1.5 dpi late and 3.5 dpi (left two columns) and in human endometrial sections from the secretory and menstrual samples (right two columns). All cells of each section are shown. Color indicates log-normalized expression, ln(counts per 10,000 + 1), on a scale running from 0 to the 99.5th percentile among the expressing cells of that gene in that species; cells above this value are shown at the maximum color. The scale is therefore shared between the two sections of a species and independent between the two species, and its upper limit is given beside each color scale. dpi: days post-induction.

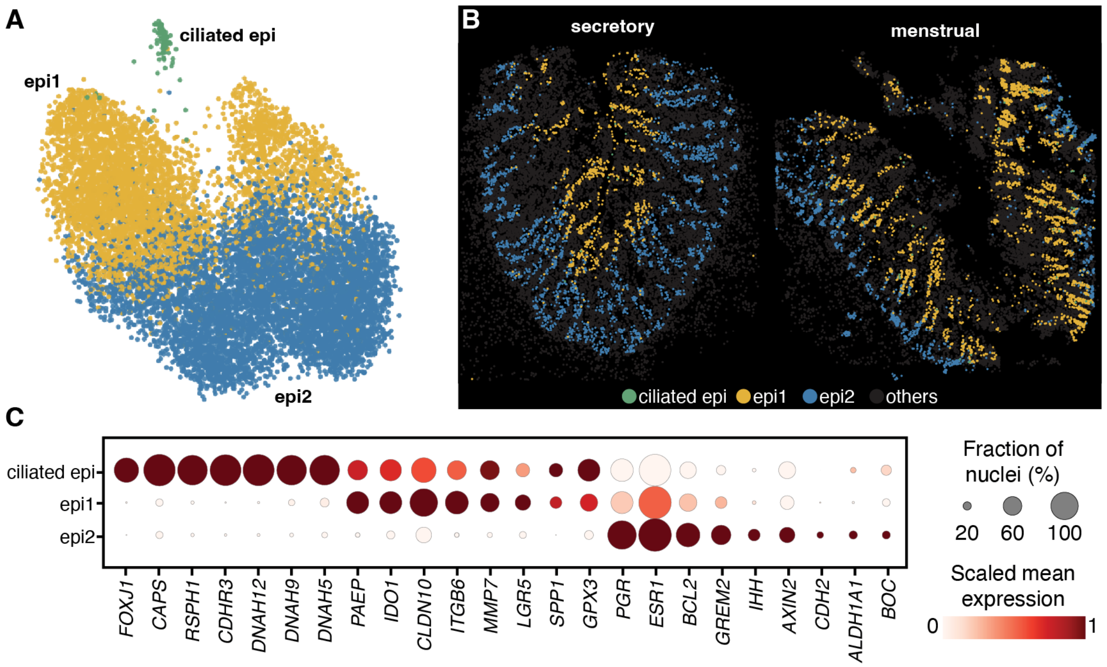


**Figure S34. Epithelial cell subclusters in the human endometrium.**
**A.** UMAP embedding of human epithelial cells, colored by subcluster. **B.** Spatial distribution of the epithelial subclusters in the secretory (left) and menstrual (right) sections, colored as in panel A and overlaid on all other cells (gray). **C.** Dot plot of marker gene expression across subclusters. Dot size indicates the fraction of cells in which the gene was detected, and color indicates mean log normalized expression scaled from 0 to 1 across subclusters for each gene. ciliated epi: ciliated epithelial cells; epi1 and epi2: epithelial cell subclusters 1 and 2.

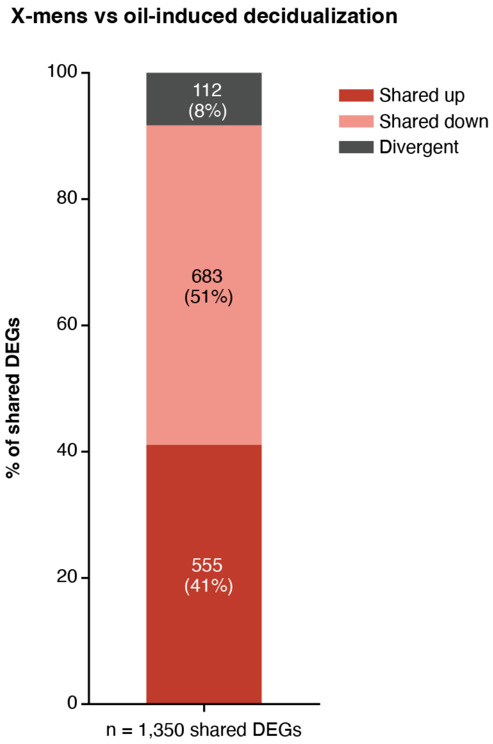


**Figure S35. Transcriptional agreement between X-Mens and oil-induced decidualization.**
Direction of change of the 1,350 genes called differentially expressed both in the published oil-induced decidualization dataset (Wang et al., 2020) and in the X-Mens comparison of decidual cells with endometrial fibroblasts (see Materials and Methods). Genes are classified as increased in both datasets (shared up), decreased in both (shared down), or changed in opposite directions (divergent); the number of genes and its percentage of the shared set are given on the bar.

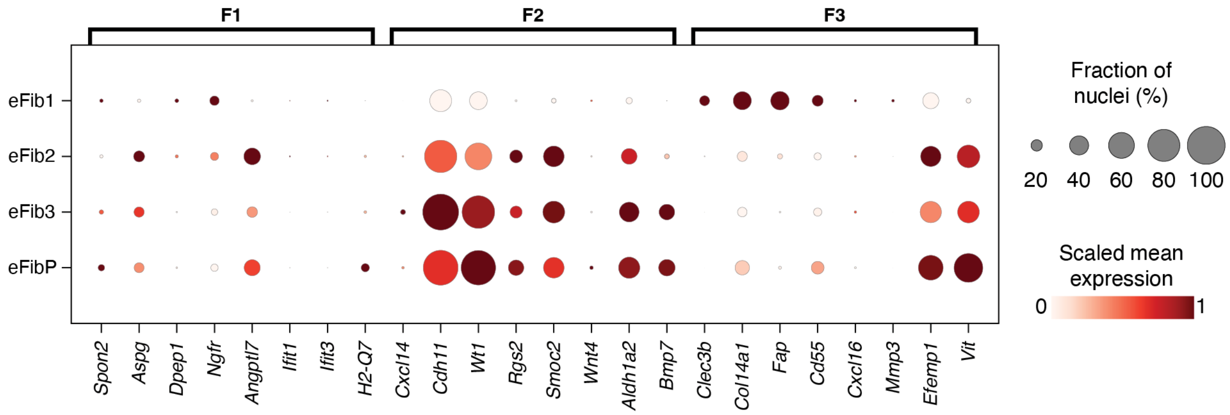


**Figure S36. Expression of published mouse endometrial fibroblast markers in the X-Mens fibroblast subtypes.**
Dot plot of the marker genes of the mouse endometrial fibroblast populations F1, F2 and F3 defined in (Kirkwood et al., 2021) (columns, grouped by population), across the endometrial fibroblast subtypes of the uninduced GsD X-Mens sample (rows); only the subtypes present in that sample are shown. Dot size indicates the fraction of cells in which the gene was detected, and color indicates mean log normalized expression scaled from 0 to 1 across subtypes for each gene. eFib1–3: endometrial fibroblast subtypes 1–3; eFibP: proliferating endometrial fibroblasts.

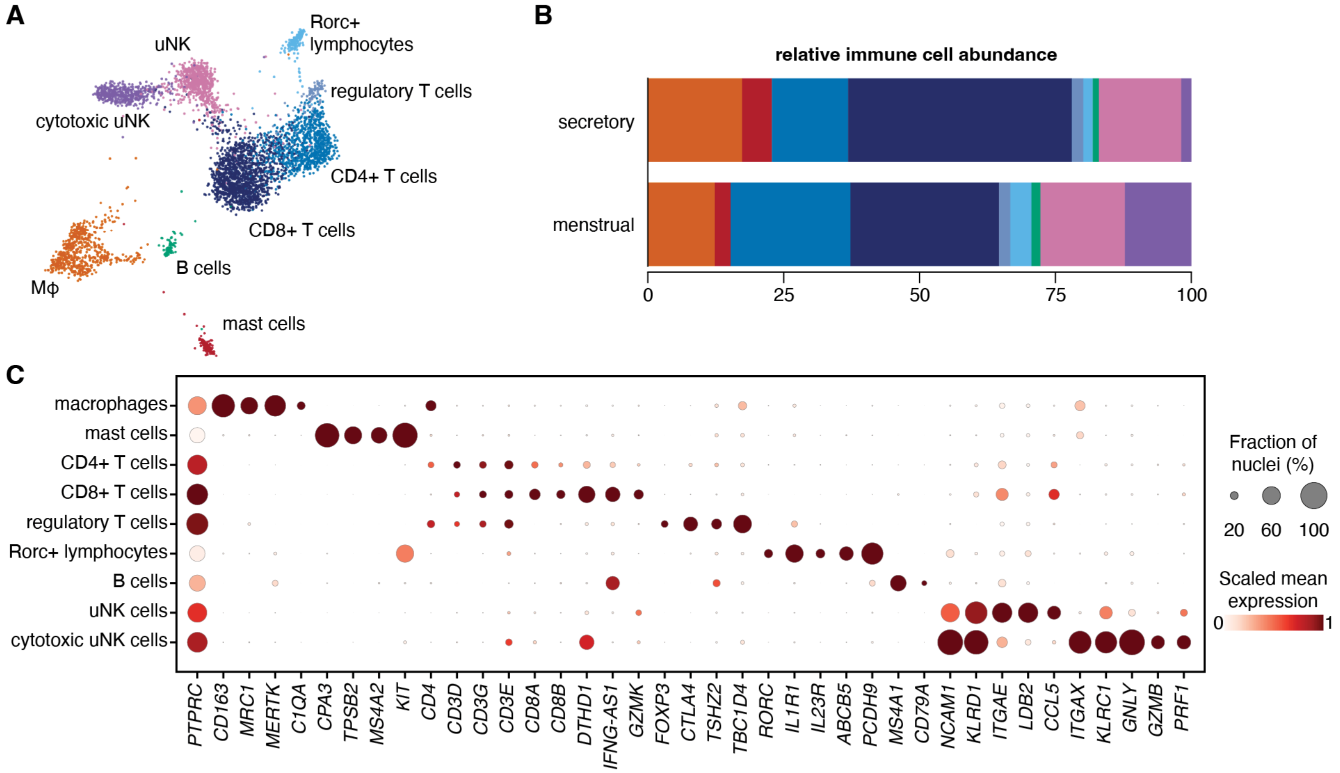


**Figure S37. Immune cell subtypes in the human endometrium.**
**A.** UMAP of immune cells subtypes from the human Slide-tags datasets, colored by immune cell subtype. Mϕ, macrophages; uNK, uterine natural killer cells. **B.** Abundance of each immune subtype, as a fraction of all immune nuclei, at the two sampled stages. Colors as in panel A. **C.** Dot plot of marker gene expression across the immune cell subtypes of the human Slide-tags samples. Dot size indicates the fraction of cells in which the gene was detected, and color indicates mean log normalized expression scaled from 0 to 1 across subtypes for each gene. uNK: uterine natural killer cells.
